# Evolution avoids a pathological stabilizing interaction in the immune protein S100A9

**DOI:** 10.1101/2022.05.09.490804

**Authors:** Joseph L. Harman, Patrick N. Reardon, Shawn M. Costello, Gus D. Warren, Sophia R. Phillips, Patrick J. Connor, Susan Marqusee, Michael J. Harms

## Abstract

Stability constrains evolution. While much is known about constraints on destabilizing mutations, less is known about the constraints on stabilizing mutations. We recently identified a mutation in the innate immune protein S100A9 that provides insight into such constraints. When introduced into human S100A9, M63F simultaneously increases the stability of the protein and disrupts its natural ability to activate Toll-like receptor 4. Using chemical denaturation, we found that M63F stabilizes a calcium-bound conformation of hS100A9. We then used NMR to solve the structure of the mutant protein, revealing that the mutation distorts the hydrophobic binding surface of hS100A9, explaining its deleterious effect on function. Hydrogen deuterium exchange (HDX) experiments revealed stabilization of the region around M63F in the structure, notably Phe37. In the structure of the M63F mutant, the Phe37 and Phe63 sidechains are in contact, plausibly forming an edge-face π-stack. Mutating Phe37 to Leu abolished the stabilizing effect of M63F as probed by both chemical denaturation and HDX. It also restored the biological activity of S100A9 disrupted by M63F. These findings reveal that Phe63 creates a “molecular staple” with Phe37 that stabilizes a non-functional conformation of the protein, thus disrupting function. Using a bioinformatic analysis, we found that S100A9 proteins from different organisms rarely have Phe at both positions 37 and 63, suggesting that avoiding a pathological stabilizing interaction indeed constrains S100A9 evolution. This work highlights an important evolutionary constraint on stabilizing mutations: they must avoid inappropriately stabilizing non-functional protein conformations.

**SIGNIFICANCE STATEMENT:** Protein stability constrains protein evolution. While much is known about evolutionary constraints on destabilizing mutations, less is known about the constraints on stabilizing mutations. We recently found a mutation to an innate immune protein that increases its stability but disrupts its function. Here we show, through careful biophysical and functional studies, that this mutation stabilizes a nonfunctional form of the protein through a direct interaction with another amino acid. We find that specific amino acids can be tolerated at each of the interacting positions individually, but not at both simultaneously. This pattern has been conserved over millions of years of evolution. Our work highlights an underappreciated evolutionary constraint on stabilizing mutations: they must avoid inappropriately stabilizing non-functional protein conformations.

## INTRODUCTION

Protein stability constrains protein evolution.^1–5^ This is intuitive for destabilizing mutations, as a mutation that disrupts the folded form of a protein will likely be deleterious. The evolutionary constraints on stabilizing mutations are, however, harder to understand. Under what circumstances does it matter that a mutation makes a protein *too* stable? Plausible arguments have been advanced for enzymes, where increasing stability decreases important dynamics related to catalysis or product release;^6–9^ however, the generic constraints on excess stability for non- enzymatic proteins remain poorly understood.

We recently identified a site in a protein that seemingly evolved to avoid deleterious stabilization,^10^ providing an excellent opportunity to probe how the requirement to avoid stabilizing mutations might constrain evolution. We isolated the mutation in question while studying the evolution of proinflammatory activity in mammalian S100A9 proteins. S100A9 proteins activate the receptor Toll-like receptor 4 (TLR4) by an as-yet poorly understood mechanism (Fig 1A).^11–14^ This activity evolved in the ancestor of all mammals when a gene shared by all amniotes (ancCG) duplicated and diverged into the last common S100A9 ancestor (ancA9) (Fig 1B).^10, 15^ Using Ancestral Sequence Reconstruction (ASR), we found that ancCG had negligible pro-inflammatory activity, ancA9 had moderate activity, and S100A9 proteins from placental mammals had high activity (Fig 1B).

**Fig 1.**
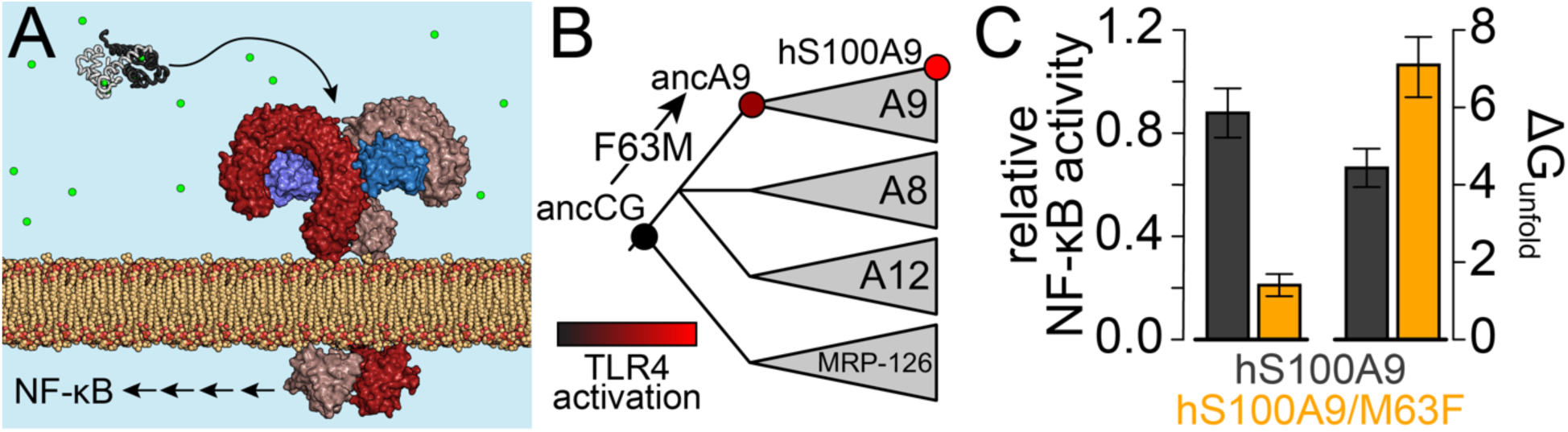
Reverting M63F in human S100A9 stabilizes the protein and disrupts proinflammatory activity. A) Schematic showing a calcium bound S100A9 dimer (white/black; PDB 1IRJ) in the calcium-rich extracellular space. S100A9 triggers dimerization of the TLR4 complex (red/blue; PDB 3FXI^16^) on the surface of immune cells, activating NF-κB and other pathways. B) Schematic phylogeny of S100A9 and its close paralogs S100A8, S100A12, and MRP-126. Wedges denote paralogs from mammals (A9, A8, and A12) or sauropsids (MRP-126). We previously characterized the indicated nodes (ancCG, ancA9, hS100A9).^10, 15^ The color of each node represents its relative ability to activate TLR4. The F63M mutation between ancCG and ancA9 was important for the evolution of S100A9’s pro-inflammatory activity. C) Reverting M63F in hS100A9 compromises its ability to activate NF-κB via TLR4 (left), while increasing unfolding free energy in 5 mM Ca^2+^ (right; experimental data shown in Fig 2).

This new pro-inflammatory function arose, in part, when Met replaced Phe at position 63 in S100A9. (Here, and throughout the manuscript, we identify amino acids by their numbering in the human sequence). Intriguingly, we found that reverting this position in human S100A9— hS100A9/M63F—both compromised the pro-inflammatory activity of the protein and increased its stability (Fig 1C).^10^ Relative to hS100A9, hS100A9/M63F required a higher concentration of urea to unfold, exhibited slower unfolding kinetics, and was much more resistant to proteolysis.^10^ We also showed that intact hS100A9, but not its proteolytic products, was the active TLR4 agonist, indicating that the mutation disrupts activity by altering the native functional state of the protein.

Here, we combine chemical denaturation experiments, structural biology, and hydrogen- deuterium exchange to understand the molecular mechanism for the stabilizing and functionally disruptive effects of this mutation. We find that a single position, Phe37, forms a “molecular staple” with Phe63. This interaction stabilizes a distorted, non-functional structure of the protein. A bioinformatic analysis reveals that S100A9 proteins frequently have Phe at either site, but rarely at both sites. Thus, for this family of immune proteins, evolution must avoid a pathological stabilizing interaction that compromises function by stabilizing non-functional protein conformations.

## RESULTS

In our previous work, we found that reverting Met63 to its ancestral Phe state in human S100A9 (hS100A9) both increases its stability and decreases its pro-inflammatory activity (Fig 1C).^10^ To better understand evolutionary constraints on stability, we set out to understand the origins of this altered stability and function.

### M63F stabilizes a calcium-bound conformation of the protein

We first hypothesized that the mutation disrupts function by stabilizing an inactive, calcium-free form of the protein. Like most S100 proteins, S100A9 forms a homodimer with two pairs of calcium binding sites.^17^ Upon calcium binding, S100A9 undergoes a conformational change that exposes a hydrophobic cleft (Fig 2A; shaded red).^17^ This cleft is the surface through which S100 proteins typically interact with target proteins.^18–22^ If the mutation stabilized an inactive, calcium-free conformation, this would explain how increased stability altered protein function.

**Figure 2.**
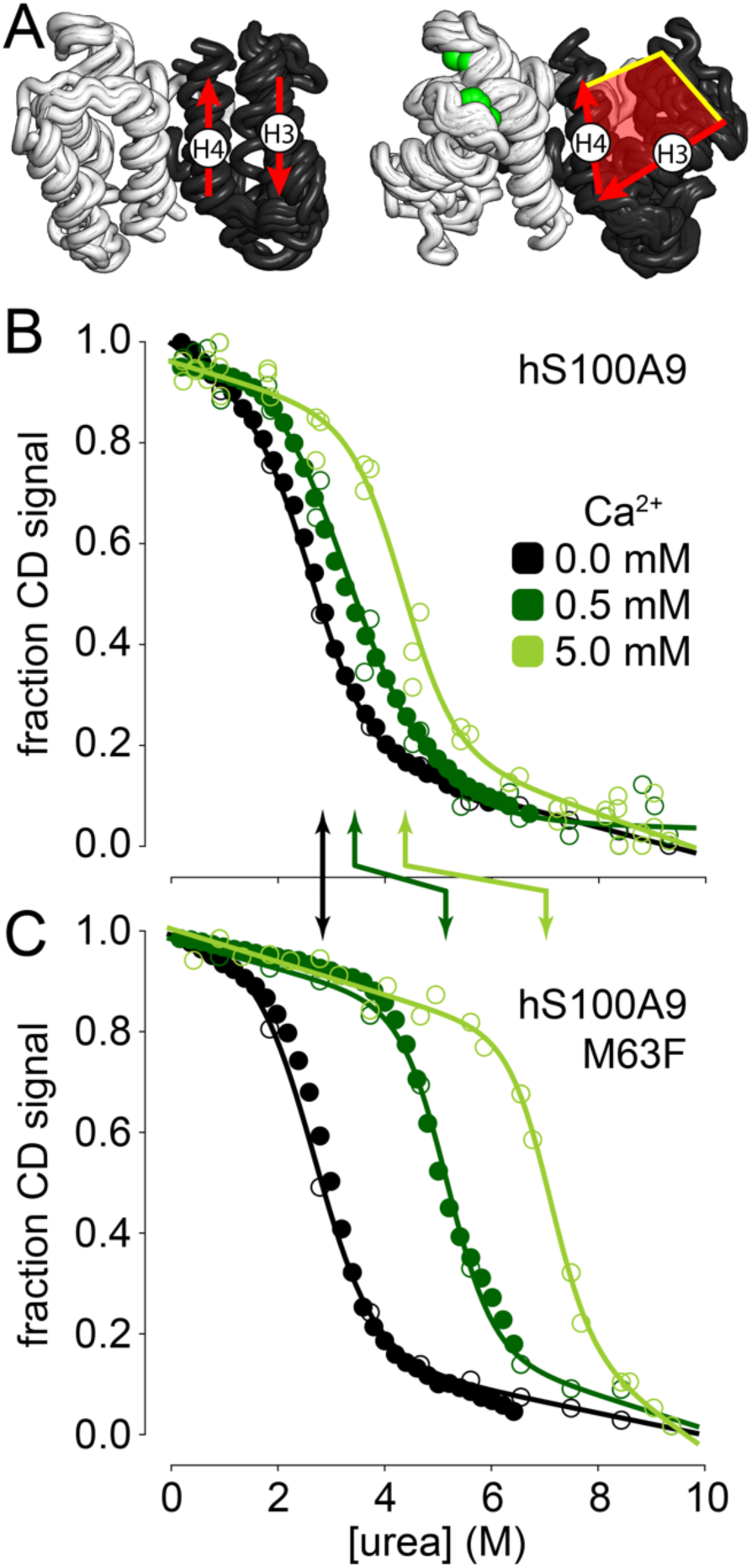
M63F stabilizes a calcium-bound form of hS100A9. A) Overlay of the structures of multiple S100 paralogs in the apo (n = 7) and calcium bound (n = 25) forms. See methods for PDB identifiers. Only residues corresponding to residues 5-85 of hS100A9 are shown; dimer chains are shown in white and black; calcium is shown as green spheres. The red arrows indicate the average orientation of helix 3 and helix 4. The red shaded area for the calcium bound structure indicates the exposed hydrophobic surface. B) Urea denaturation of hS100A9 in 0 (black), 0.5 (dark green), and 5 mM (light green) Ca^2+^. y-axis is circular dichroism signal at 222 nm, normalized from 0 to 1. Closed circles are automated titrations; open circles are manual titrations. Lines are the best fit model to all experiments for that calcium condition. C) Denaturation of hS100A9/M63F. Colors and symbols as in panel B. The arrows between panels B and C indicate the midpoint of denaturation (C_m_, in M) for the two proteins.

To test this hypothesis, we determined which conformation(s) of hS100A9 were stabilized by M63F. We measured the unfolding free energy (ΔG_unfold_) of hS100A9 and hS100A9/M63F in increasing concentrations of calcium: 0 mM, 0.5 mM, and 5 mM Ca^2+^. We measured equilibrium unfolding of the protein in urea by circular dichroism (CD) spectroscopy monitored at 222 nm, which reports on protein α-helical content. We fit a two-state model to the data, which had free parameters describing the unfolding free energy of the protein in water (ΔG_unfold_), the dependence of unfolding energy on urea concentration (m-value), and the native and denatured baselines.^23^. We analyzed the data in two ways. First, we fit a global m-value for all protein variant/calcium concentrations. This is a reasonable assumption for hS100A9 and hS100A9/M63F, as the proteins differ by a single amino acid and the m-value is strongly correlated with the amount of hydrophobic surface area that is exposed upon protein unfolding.^24^ These are the data presented throughout the text and figures. Second, we allowed the m-value to vary for each protein/calcium condition. The two approaches gave qualitatively similar outcomes (Fig S1, Table S1, Table S2); however, we note where they diverge in the text.

In the absence of calcium, we observed no significant difference in apparent stability between hS100A9 and hS100A9/M63F: 2.6 ± 0.5 vs. 2.4 ± 0.5 kcal/mol (Fig 2B,C). Adding calcium stabilizes both hS100A9 and hS100A9/M63F, as expected from Le Chatelier’s principle. The effect was, however, much more dramatic for the M63F mutant than hS100A9. At 5 mM Ca^2+^, hS100A9 and hS100A9/M63F have stabilities of 4.3 ± 0.5 vs. 7.0 ± 0.5 kcal/mol, respectively. This can be seen visually by the much larger calcium-induced increase in urea melting midpoint for the mutant compared to the wildtype protein (Fig 2B,C; see Table S1 for fit parameters).

Because the mutation had no effect in the absence of calcium, but exhibits an increasing effect size with increasing calcium, we concluded that the mutation preferentially stabilizes a calcium-bound form of the protein.

### M63F distorts the structure of hS100A9

It was surprising that a mutation compromised activity while stabilizing a calcium-bound form of hS100A9. hS100A9 activates TLR4 in the extracellular space, where the concentration of calcium is two orders of magnitude above the K_D_ of binding for hS100A9 to calcium^25^ (Fig 1A); therefore, the calcium-bound state is presumably the active form of the protein.^11–13^ Further, for most members of the S100 protein family, the calcium-bound conformation preferentially interacts with downstream target proteins.^18–21^ We also showed previously that the intact protein is the proinflammatory species of the protein.^10^ How could stabilization of a calcium-bound state disrupt activity?

To answer this question, we solved the structure of the calcium-bound form of hS100A9/M63F in the presence of 10 mM CaCl_2_ by NMR. We transferred and confirmed peak assignments for hS100A9/M63F from the calcium-loaded wildtype hS100A9 NMR structure^26^ using 3D NMR experiments and successfully assigned peaks for 91 out of 114 residues in hS100A9/M63F (Fig S2). The majority of unassigned peaks are within the S100A9 disordered C-terminal tail, most of which are also unassigned in the wildtype hS100A9 structure.^26^

The resulting ensemble of structures has a core set of residues (6-86) with an average pairwise backbone heavy atom RMSD of 0.9 Å. Detailed structural statistics are available in Table 1. The remaining residues (1-5 and 87-114) are poorly constrained by the NMR data (Fig 3A).

**Fig 3:**
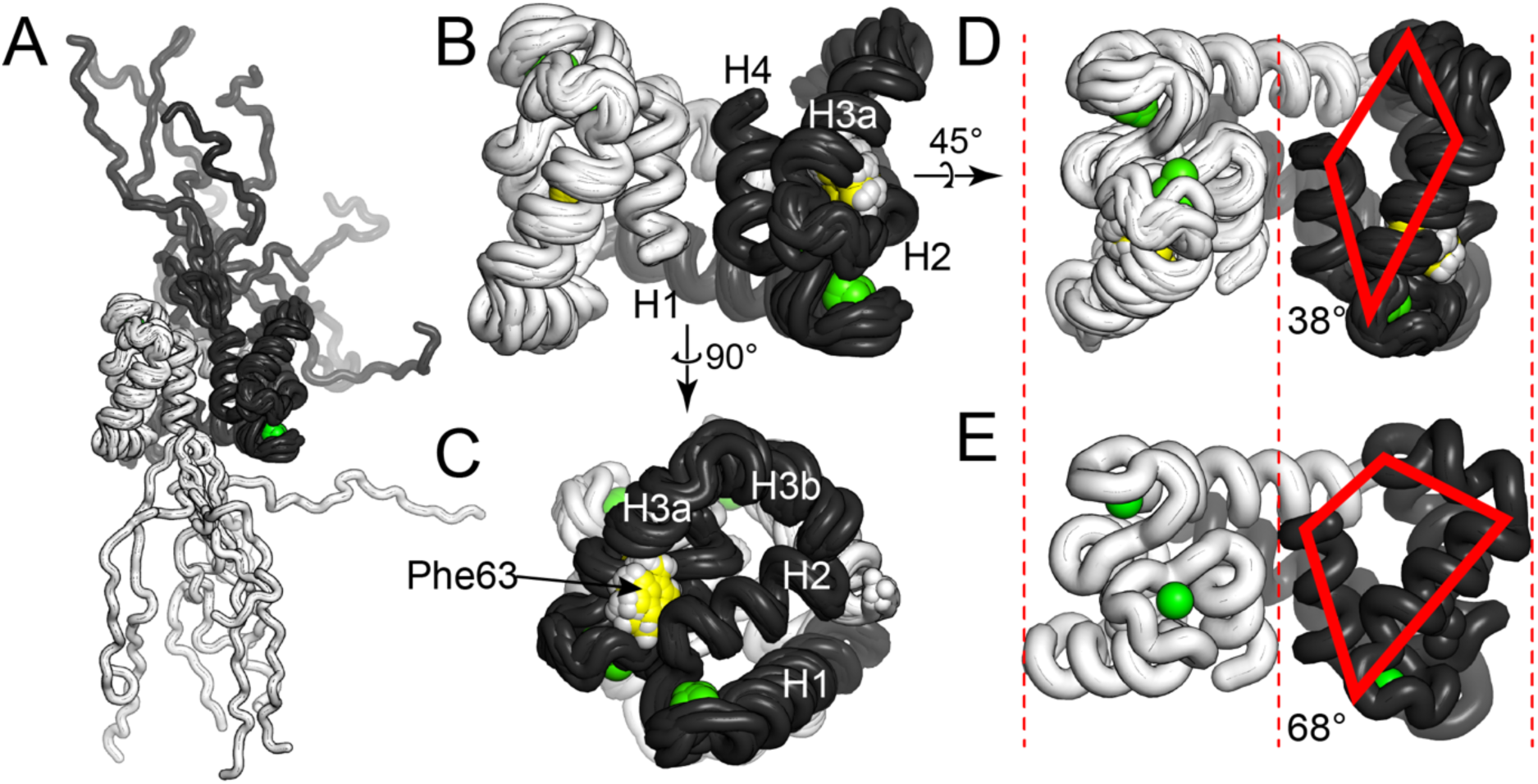
M63F distorts the structure of hS100A9. **A)** Overlay of 10 models from the NMR ensemble generated for hS100A9/M63F. The A and B chains are shown as white and black, calcium ions in green. Residues 1-5 and 87-114 are poorly constrained in the structure. B-C) Models for residues 6-86. Panel C is rotated 90° relative to B. Helices 1, 2, 3a, 3b, and 4 are indicated on the structure. Phe63 is highlighted in yellow. D) Structure shown in B, rotated 45° along indicated axis. The solid red lines highlight the S100A9 binding surface (as in Fig 2A). E) Crystal structure of hS100A9 (PDB ID: 1IRJ)^17^ in same orientation as structure in panel D. The solid red lines highlight the hydrophobic binding cleft. The red dashed lines highlight the increased compaction of hS100A9/M63F relative to hS100A9.

**Table 1:** NMR structure statistics

|  |  |  |
| --- | --- | --- |
| **Distance restraints** |  |  |
| Total NOE |  | 6136 |
| Unambiguous |  | 3562 |
| Ambiguous |  | 2574 |
| H-Bond Restraints |  | 132 |
| Total Dihedral Restraints |  | 330 |
| <sup>3</sup>J HNHA coupling constants |  | 90 |
| Total restraints |  | 6688 |
| **Structure statistics** |  |  |
| Violations |  |  |
| r.m.s.d of distance violation/constraint |  | 0.022 |
| r.m.s.d of dihedral angle violation/constraint |  | 0.294 |
| r.m.s.d from ideal geometry |  |  |
| Bond length |  | 0.002 |
| Bond angles |  | 0.510 |
| **Average pairwise r.m.s.d** |  |  |
| Ordered backbone |  | 0.9 |
| All backbone |  | 12.3 |
| Ordered heavy atoms |  | 1.3 |
| All heavy atoms |  | 12.3 |
| **Ramachandran statistics** |  |  |
| Most favored regions |  | 98.4% |
| Allowed regions |  | 1.6% |
| Disallowed regions |  | 0% |
| **Average Molprobit Clashscore** |  | 7.73 |
| **RCSB PDB ID** |  | 7UI5 |

This is consistent with previous work showing that the N- and C-termini of hS100A9 are disordered, both in solution^26^ and in a crystal structure of the protein.^17^ The well-ordered region of the protein adopts the same basic fold as observed in the crystal structure of wildtype hS100A9 (Fig 3B). The average pairwise backbone heavy atom RMSD between the hS100A9/M63F NMR ensemble and the hS100A9 crystal structure (PDB ID: 1IRJ) is 2.45 ± 0.2 Å.

In the hS100A9/M63F structure, Phe63 is in the same general orientation as Met63 in the wildtype structure (Fig 3C). This leads, however, to a change in the relative orientations of the helices that make up the protein binding surface of S100A9: H2, H3a, H3b, and H4. In the crystal structure of the wildtype protein, a relatively large hydrophobic surface is exposed (red trapezoid, Fig 3E). In contrast, this surface is significantly collapsed for hS100A9/M63F (Fig 3D). This change is accompanied by an overall compaction of the structure (red dashed lines, Fig 3D,E). The full magnitude of this change is more easily seen with a video morphing between the two structures (Video S1).

One way to quantify this change is by measuring the angle between H3b and H4. In the mutant NMR ensemble, this angle is 38.3° ± 2° (Fig 3D). In the wildtype crystal structure (pdb 1irj), this is 68.0° (Fig 3E). To put these numbers in context, we measured the angles between these helices for experimental structures of other members of the S100 protein family solved in both the absence and presence of calcium (shown in Fig 2A). For apo structures, we found an angle of 16.7° ± 4° (n = 7); for calcium-loaded structures, we found an angle of 78.6° ± 23° (n = 25) (Table S3). Thus, the M63F mutation pushes the structure toward the apo form even when calcium is bound, collapsing the hydrophobic protein-interaction patch typically exposed upon binding of calcium.

### Phe63 stabilizes an adjacent helix

We next sought to understand why this mutation causes this structural effect. To do so, we employed hydrogen-deuterium exchange measurements monitored by mass spectrometry (HDX-MS).^27, 28^ In this experiment, the relative rates of deuterium uptake for backbone amides report on the local stability and solvent accessibility of those positions. HDX–MS monitors the exchange of amide hydrogens in the peptide backbone with solvent hydrogens (in this case, deuterium). An individual amide’s ability to undergo exchange is thus directly related to its solvent-accessibility and local stability.^29, 30^ We monitored HDX on hS100A9 and hS100A9/M63F, quenching the reaction at various timepoints (lower pH and temperature). We digested each quenched sample using a combination of porcine pepsin and aspergillopepsin inline under acidic conditions and quantified the number of deuterium atoms per peptide by mass spectrometry (Table S4). We then calculated the percent of backbone amides that exchanged hydrogen for deuterium for each peptide at each timepoint. For details of the HDX experiment, see methods.

Fig 4A shows a plot of the difference in deuterium uptake for peptide fragments of hS100A9 and hS100A9/M63F after one hour of exchange, where most peptides are in the transition and not yet fully exchanged. The M63F mutation causes a striking shift towards increased protection across the entire protein sequence. 60% of peptides exhibit lower exchange in the mutant relative to wildtype, while no position in the mutant exhibits statistically significant greater exchange than wildtype (histogram in Fig 4A). When mapped to the NMR structure (Fig 4B), the HDX results point to increased stabilization of H1, H2, H3a, and H4. The most striking stabilization occurs between residues 30-39 in H2 (arrow, Fig 4A). This region is directly adjacent to Phe63 in the NMR structure. Peptides containing Phe37, in particular, are highly stabilized in the mutant relative to wildtype. In the NMR structure, the Phe37 sidechain forms an apparent edge-face π-stack with Phe63. This interaction bridges H2 and H3b and could conceivably be the proximate cause for the distorted orientation between these two helices.

**Fig 4.**
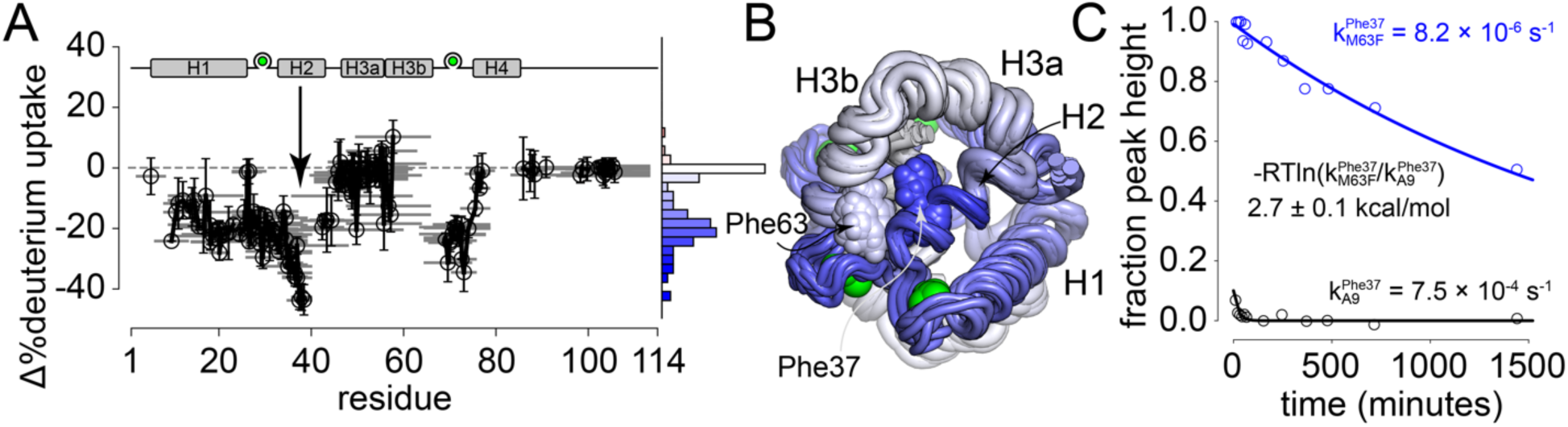
Phe63 stabilizes an adjacent helix. A) Difference in peptide deuterium uptake observed by mass spectrometry between hS100A9 and hS100A9/M63F after one hour. Gray lines indicate peptides observed. Circles indicate peptide centers. Errors represent uncertainty in difference in deuterium uptake for that peptide, as calculated by HDExaminer using default settings. Histogram on right indicates frequency of peptides with differences in deuterium uptake, colored slowest (blue) to fastest (red) uptake. B) Uptake values from panel A mapped to the NMR structure of hS100A9/M63F using color and tube radius. To assign a value to each site, we took the average change in deuterium uptake for all peptides containing a given site. Colors correspond to the histogram shown in panel A. C) Change in Phe37 backbone amide ^1^H/^15^N peak intensity when diluted into D_2_O. Lines show exponential decay model fit to the data. Colors denote protein: hS100A9 (black) and hS100A9/M63F (blue).

We next tested the hypothesis that Phe37 is, indeed, stabilized by the M63F mutation. We repeated our HDX experiment using NMR, which provides site-specific rather than peptide- specific information. Using this approach, we were able to quantify differences in exchange rate at 25 °C for 25 matched amide backbones in hS100A9 and hS100A9/M63F (Fig S3, Fig S4, Table S5). This includes five sites between residues 31 and 39 (31, 32, 37, 38, 39). All five sites exchanged between 60 and 600 times slower in the M63F mutant relative to hS100A9. As shown in Fig 4C, the Phe37 backbone amide exchanges 92-fold slower in hS100A9/M63F (8.2 × 10^-6^ s^-1^) relative to hS100A9 (7.5 × 10^-4^ s^-1^). This corresponds to an apparent stabilization of 2.7 ± 0.1 kcal/mol. Thus, consistent with the HDX-MS experiments, our HDX-NMR experiments reveal that M63F strongly stabilizes Phe37 and other residues on H2.

### Phe37 and Phe63 form a strong stabilizing interaction

Based on the NMR structure and our HDX data, we hypothesized that the Phe37/Phe63 interaction is the underlying cause of the stabilized, non-functional structure of the protein. We therefore constructed a mutant cycle measuring the effect of the Phe/Phe interaction on the stability and function of hS100A9. We constructed the cycle considering the evolutionary history of positions 63 and 37. The M63F mutation is a reversion to the ancestral state at position 63 (Fig 1B). Over the same evolutionary interval, an ancestral Leu at position 37 evolved to Phe.^15, 31^ We therefore constructed a mutant cycle looking at all combinations of Leu/Phe at position 37 and Met/Phe at position 63.

We used the protein hS100A9/F37L as the reference state for this cycle. This protein does not have phenylalanine at positions 37 or 63 (L/M). By using it as our reference, we could measure the effects of introducing phenylalanine at position 37 (F/M; hS100A9), at position 63 (L/F; hS100A9/F37L/M63F), or at both positions (F/F; hS100A9/M63F). This allowed us to measure any non-additive effect of the F/F pair on protein stability.

In the absence of calcium, we found that the stabilities of all four hS100A9 variants were indistinguishable within experimental uncertainty (Fig 5A, ∼3 kcal/mol). As a result, there is no interaction between the M63F and L37F mutations outside experimental uncertainty (Fig 5A inset, -0.2 ± 1 kcal/mol). With the addition of 5 mM Ca^2+^, a strong interaction emerged (Fig 5B). Introducing L37F or M63F alone mildly increased stability (0.9 ± 1 and 0.8 ± 1 kcal/mol, respectively); introducing them together had a much larger effect (2.7 ± 1 and 2.8 ± 1 kcal/mol). This works out to a coupling energy of 1.9 ± 1 kcal/mol for the introduction of the new Phe37/Phe63 interaction (Fig 5B inset; key energy terms underlined). All fit parameters are reported in Table S1. (When we re-analyzed our results with individual m-values for each protein/calcium condition, we obtain an even higher coupling energy: 4.8 ± 2 kcal/mol; Fig S5, Table S2).

**Fig 5.**
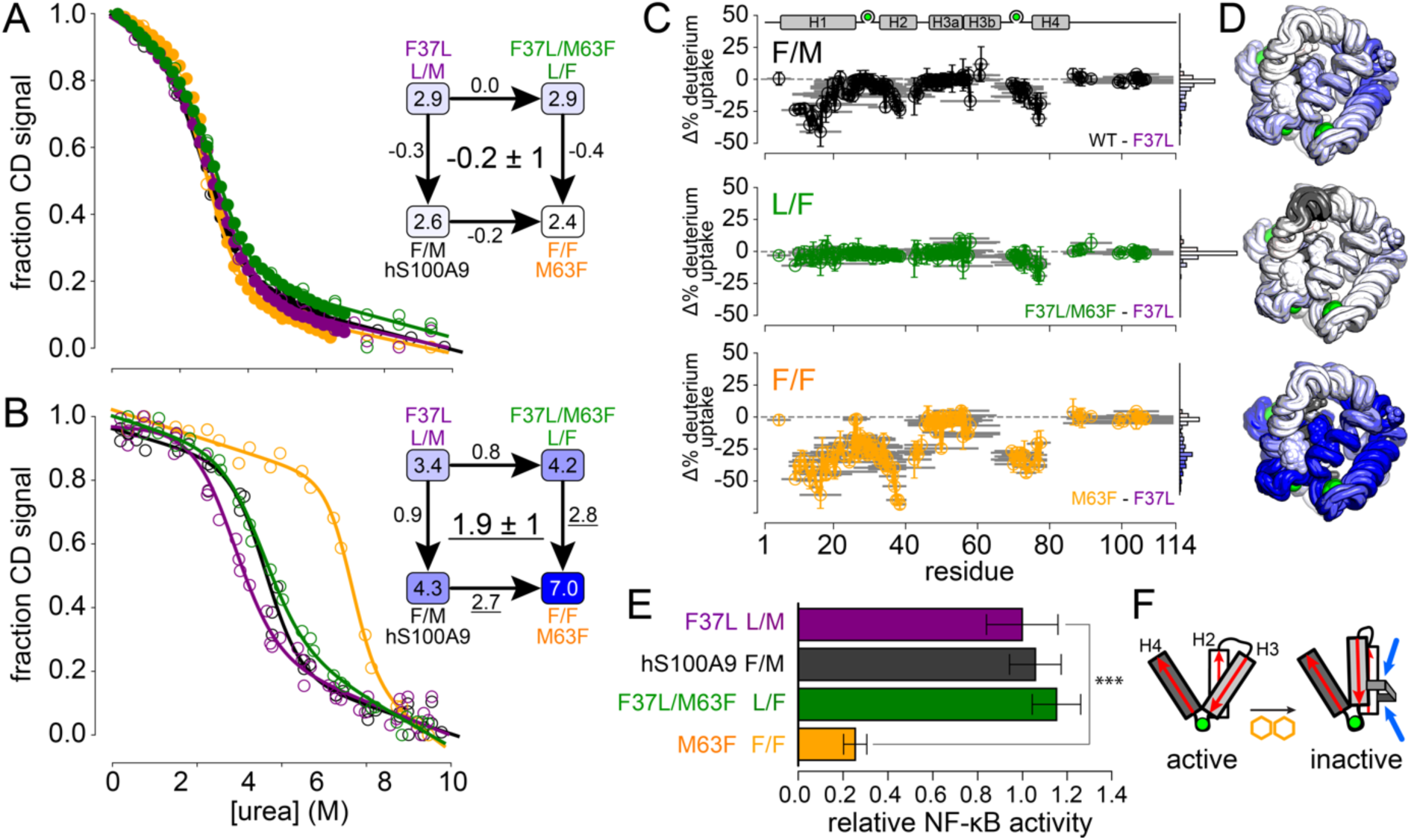
Phe37 and Phe63 form a stabilizing and functionally disruptive interaction. A) Urea denaturation of S100A9 proteins in 0 mM Ca^2+^. Colors represent the variant: hS100A9 (black), hS100A9/F37L (purple), hS100A9/M63F (orange), and hS100A9/F37L/M63F (green). Closed circles are automated titrations; open circles are manual titrations. Lines show best fit model to all experiments for each variant at this calcium condition. The inset shows a mutant cycle based on the ΔG_unfold_ values extracted from fits shown (ΔG values in kcal/mol). Values along arrows are the effect of each mutation (ΔΔG_unfold_). The number in the center indicates the coupling between Phe at each site. B) Unfolding curves and mutant cycle for unfolding at 5 mM Ca^2+^. Visual elements are as in panel A. Stabilizing interactions noted in text are underlined. C) Effects of Phe mutations on deuterium uptake after one hour, calculated as the difference in the uptake of the indicated variant relative to hS100A9/F37L. Colors as in panel A. Graphical elements described in Fig 4A. D) Change in deuterium uptake mapped to the NMR structure of hS100A9/M63F for each variant in panel C. Colors follow histograms in panel C. E) Protein variants (2 μM) activating an NF-κB luciferase reporter in HEK-293T cells transiently transfected with human TLR4, MD-2, CD14, and reporter plasmids. Bar lengths are the means of at least five biological replicates; error bars are the standard errors on the mean. “***” indicates a p-value < 0.001 (t-test). Scale is normalized such that the activity of hS100A9/F37L is 1.0. F) Schematic showing relative orientations of helices 2-4 for a single chain in an the active versus inactive structure. The aromatic/aromatic pair (yellow) forms a molecular “staple” (blue) that stabilizes the inactive conformation of the protein.

We next set out to validate the structural effect of the Phe/Phe interaction. We performed HDX-MS experiments on hS100A9/F37L and hS100A9/M63F/F37L, and then calculated the change in deuterium uptake across sites relative to the hS100A9/F37L background. Fig 5C shows the differences in uptake induced by F37 alone (black), F63 alone (green), or F37/F63 together (orange). When mapped to the structure, we found that residues 30-39 (H2) and residues 70-80 (H4) are massively more stabilized by the Phe/Phe pair than either Phe introduced alone (Fig 5C- D). As with the ΔG_unfold_ data, there is a strong synergistic stabilizing effect when F37 and F63 are introduced together.

### The Phe37/Phe63 interaction directly disrupts activity

We next tested whether this interaction was responsible for the loss of proinflammatory activity in hS100A9/M63F. We measured the ability of the four hS100A9 variants to activate NF-κB via TLR4 in a luciferase reporter assay. We found that the variant with no Phe (L/M) and the variants with a single Phe (F/M and L/F) activated TLR4 indistinguishably from one another (Fig 5E). In contrast, the F/F construct (hS100A9/M63F) had strongly compromised proinflammatory activity (Fig 5E).

Our work shows that it is not Phe at position 63 that disrupts function, but rather the combined effect of Phe at both positions 37 and 63. This leads us to the structural model shown in Fig 5F: the interaction between Phe/Phe creates a new “molecular staple” between H3 and H2. This causes H3 to rotate relative to H4, thus distorting the structure of the molecule and disrupting proinflammatory function.

### Evolving S100A9 proteins avoid having Phe at both position 37 and 63 simultaneously

Having identified a pair of amino acids that, together, stabilize a non-functional form of the protein, we looked for evidence that this shaped the evolution of S100A9 orthologs. We constructed an alignment containing 432 protein sequences of S100A9 and its close paralogs from across amniotes (Table S6). The proteins S100A9, S100A8, and S100A12 are found in mammals; MRP-126 is found in birds and reptiles (sauropsids). Although the relationship between these clades is difficult to establish with certainty, the most parsimonious scenario places MRP-126 as co-orthologous to S100A8, S100A9, and S100A12, which formed by serial duplication of the MRP-126 gene after the divergence of sauropsids and mammals (Fig 6A).^10, 15^

**Fig 6.**
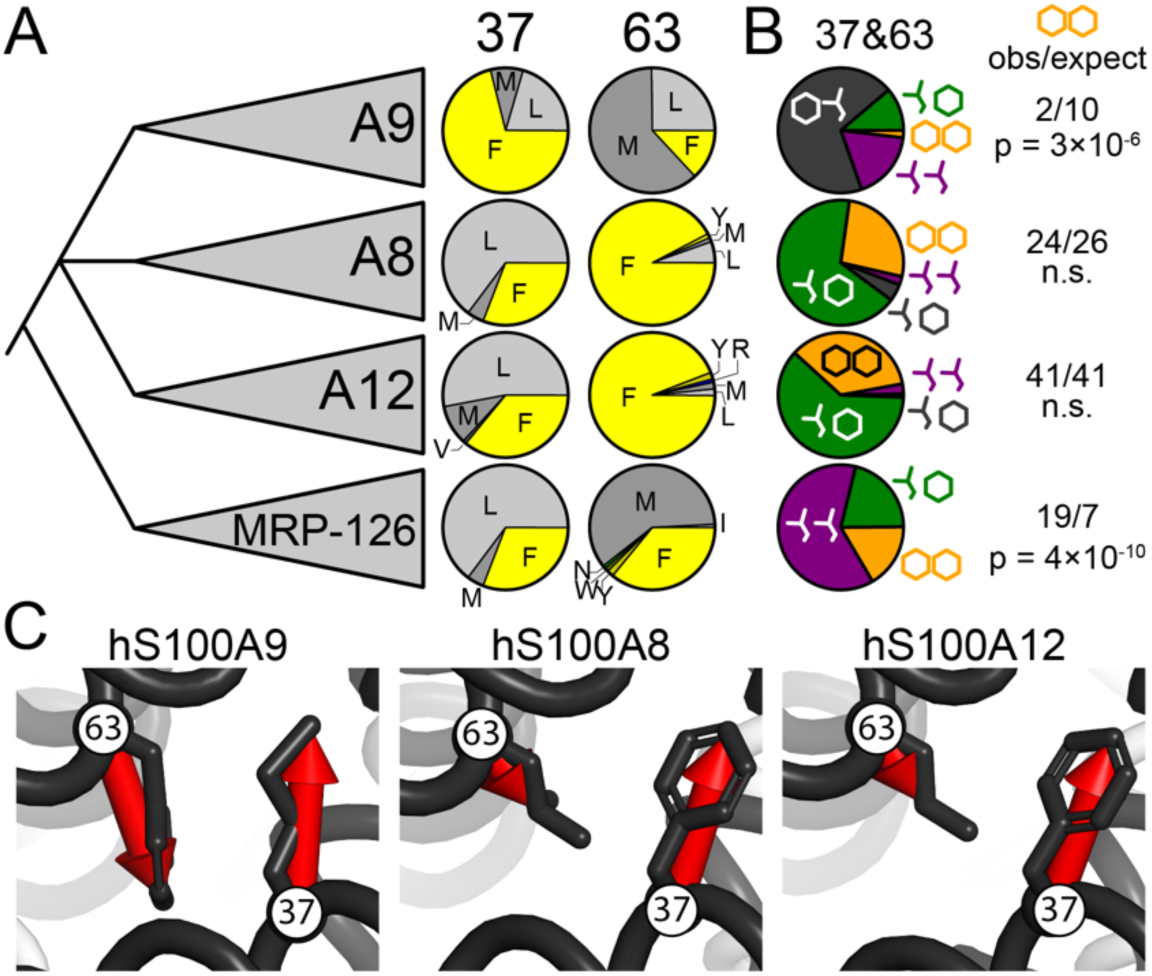
The Phe37/Phe63 state is rare in S100A9 proteins. A) Frequencies of amino acids at sites 37 and 63 extracted from an alignment of 432 sequences containing four clades of S100 proteins (labeled on left). Pie wedges are colored according to amino acid class: aromatic (yellow), non- aromatic/hydrophobic (gray), polar (green), or basic (blue). B) Joint frequencies of non-aromatic and aromatic residues at positions 37 and 63 in each clade. Schematic Leu sidechain represents hydrophobic (Met, Leu, Val, or Ile in alignment); hexagon represents aromatic (Phe, Tyr, or Trp in alignment). Colors follow Fig 5: non-aromatic/non-aromatic (purple), aromatic/non-aromatic (gray), non-aromatic/aromatic (green), and aromatic/aromatic (orange). Numbers to right indicate observed and expected number of aromatic/aromatic pairs observed in that clade given the aromatic frequency at each site; p-value is the result of a Fisher’s exact test. C) Relative orientations of residues 37 and 63 in experimental structures of hS100A9 (PDB: 1IRJ),^17^ hS100A8 (1MR8),^32^ and hS100A12 (1E8A).^33^ Red arrows connect the C_α_ and the most distal atom of the sidechain.

Both positions 37 and 63 strongly prefer Leu, Met, or Phe to the exclusion of other amino acids in all four paralogs (Fig 6A). S100A9 has noticeably different preferences than S100A8 or S100A12. Of the 107 S100A9 sequences in the alignment, 71% have Phe at position 37 and 13% have Phe at position 63. This preference is reversed relative to S100A8 and S100A12, which have ∼30% Phe at position 37 and ∼90% Phe at position 63 (Fig 6A).

We next asked whether we saw patterns in the co-occurrence of Phe at positions 37 and 63 by measuring the frequencies of all four possible combinations of non-aromatic and aromatic residues at these two sites (Fig 6B). This is directly analogous to our mutant cycles shown in Fig 5. For S100A9, we found that aromatic/non-aromatic—e.g. Phe37/Met63—is the most common state. This is followed by non-aromatic/non-aromatic and non-aromatic/aromatic. Only 2 of the 107 sequences have aromatic/aromatic at both sites. If aromatic residues occurred independently at these sites, we would expect 10 sequences to have an aromatic/aromatic pair (71% × 13% × 107 sequences = 9.8). This depletion is statistically significant (p = 3 × 10^-6^; Fisher’s exact test). Thus, it appears that evolving S100A9 proteins can readily tolerate Phe at either position but avoid placing Phe at both positions simultaneously.

Such depletion is not observed for the other paralogs. S100A8 and S100A12 show no significant deviation in observed aromatic/aromatic pairs from what is expected given the frequency of aromatic residues at sites 37 and 63 (Fig 6B). For MRP-126, we see relative enrichment of Phe/Phe pairs: we observe 19 pairs, but only expect 7. Thus, it appears that the S100A9 clade has a unique evolutionary constraint to avoid placing Phe at both position 37 and 63 simultaneously. Given our experimental observations, a plausible constraint is avoiding the pathological stabilization of a non-functional conformation of the protein.

## DISCUSSION

This work reveals how reverting a single residue to its ancestral amino acid in hS100A9 decreases its proinflammatory activity. The mutation creates a new stabilizing interaction between Phe37 and Phe63. This “molecular staple” reorients a helix, thus occluding the hydrophobic surface on S100A9 that is typically exposed upon calcium binding and thought to be critical for function.^12, 13, 34^ We found that Phe can be tolerated at either site individually, but not at both sites together. This biochemical and functional constraint has left a detectable evolutionary footprint in the sequence preferences of evolving S100A9 proteins.

This evolutionary rule against Phe/Phe pairs appears to be specific to S100A9, but not other closely related proteins (Fig 6B). Its closest paralogs—S100A8 and S100A12—exhibit no aversion for Phe/Phe. An earlier-diverging protein—MRP-126—even prefers Phe/Phe over other amino acid pairs at these positions. This differential preference for Phe/Phe can be understood structurally. Fig 6C shows the equivalent 37/63 positions in the structures of hS100A9, hS100A8, and hS100A12.^17, 32, 33^ In hS100A9, these two residues are directly parallel and adjacent, allowing formation of the Phe/Phe contact. In hS100A8 and hS100A12, these residues are slightly offset, presumably allowing Phe to be accommodated at both sites. While a structure of MRP-126 has not yet been solved, the observed preference for Phe/Phe in MRP-126 (Fig 6C) indicates that these proteins can structurally accommodate both aromatic residues. This suggests that the MRP-126 structure might more closely resemble S100A8 and S100A12 structures at positions 37 and 63.

This change in structure and amino acid preference occurred on the same evolutionary interval over which S100A9 evolved potent proinflammatory activity (Fig 1B).^10, 15^ It is unclear if this structural change was important for the acquisition of proinflammatory activity; however, once established, the new structure created a new constraint on sequence evolution: Phe is allowed at either site in S100A9s, but not at both together. This is an example of evolutionary entrenchment.^35–40^ The altered structure of S100A9 relative to its close paralogs makes a previously allowable Phe/Phe pair functionally deleterious, thus preventing reversion to the ancestral state of the protein.

We started this work asking what evolutionary constraints exist on stabilizing mutations. We can now answer this question for S100A9: the constraint at position 63 is not to avoid stabilizing the native conformation, but rather to avoid stabilizing a non-native conformation. In hS100A9, the most stable structure is the functional structure. In hS100A9/M63F, the most stable structure is a distorted, non-functional structure. A single interaction, between Phe37 and Phe63, is sufficient to stabilize this non-functional conformation. The fact that this can be achieved with a single mutation indicates that the non-functional conformation in wildtype S100A9 is only slightly less stable than the functional conformation in hS100A9—likely only a few kcal/mol given the apparent interaction energy between F37 and F63 (Fig 4C, 5B).

From one perspective, the existence of an energetically similar but non-functional conformation is surprising. Selection for protein stability can lead to both positive design constraints (optimizing the native structure) and negative design constraints (deoptimizing non- native structures).^41, 42^ In an ideal case, evolution would find robust protein sequences for which no single mutation is sufficient to switch the protein from its native to a non-native structure.^42–46^ S100A9 may be less robust than expected because the protein undergoes a conformational change in response to calcium (Fig 2A). The same protein sequence must therefore be compatible with two different structures. As a result of this degeneracy, the energy landscape for the protein may be interspersed with energetically similar, but non-functional, conformations.^47–51^ In this scenario, evolution cannot purge these structures from the energy landscape without disrupting the ability of the protein to respond to calcium.

Whatever their evolutionary origin, our work demonstrates that energetically similar, non- functional conformations constrain the evolution of protein sequences. Proteins exist as ensembles of structures, some highly populated, others rarely seen.^52–55^ The same mutation can have different effects on different structures within this energetic ensemble, redistributing their relative populations.^52, 53, 56^ This means that proteins must not only avoid mutations that destabilize native structures, but also avoid mutations that stabilize non-native structures. As a result, we need to think of constraints on protein evolution in terms of multiple conformations—not simply in terms of the native conformation. Or, in the memorable words of Gregorio Weber, we should think of evolving proteins not as “platonic” single structures, but as “kicking and screaming stochastic molecules.”^57^

## MATERIALS AND METHODS

Raw data and scripts to perform all analyses and generate all figures are posted as a freely downloadable repository on github: https://github.com/harmslab/stability-constraint-ms-figures.

### Protein expression and purification

We used a Cys-free variant of hS100A9 in a pETDuet-1 plasmid as our wildtype construct. This corresponds to Uniprot P06702 with a C3S mutation. We introduced the F37L and M63F mutations by site-directed mutagenesis (Agilent). We expressed the recombinant proteins in *E. coli* BL21 (DE3) pLysS Rosetta cells grown in luria broth (LB) containing both ampicillin and chloramphenicol. We streaked from glycerol stocks onto LB plates, inoculated overnight cultures from single colonies, and grew overnight cultures at 37°C/250 rpm. The next morning, we inoculated 1.5 L of media with 10 mL of overnight culture, then allowed this to grow at 37°C/250 rpm to an OD_600_ between 0.6 – 1.0. To produce isotopically labeled protein samples for NMR, we grew 2 × 1.5 L cultures to an OD_600_ between 0.6 – 1.0 as above. Cells were then pelleted at 3,000 rpm for 20 min, resuspended in 750 ml of 1X M9 minimal media, and grown at 37°C/250 rpm for 45 minutes. M9 media was prepared as follows: 1L of 5X M9 stock solution was prepared by adding 30 g of NA_2_HPO_4,_ 15 g of KH_2_PO_4,_ 2.5 g NaCl to 1L of ddH_2_O, and then autoclaved for 15 mins. 750 mL of 1X M9 media for labeling was prepared by combining 150 mL of 5X M9 stock solution, 15 ml of 50% glucose (not included for ^13^C labeled samples), 7.5 mL of Basal Medium Eagle Media (Thermo Fisher Scientific), 1.5 mL of 1M MgSO_4_, 75 μL of 1M CaCl_2_, 0.75 g of ^15^NH_4_Cl (Sigma Aldrich) for ^15^N labeling (and 0.75 g of ^13^C glucose (Sigma Aldrich) for ^15^N/^13^C double-labeling), and 750 μL of 1000X antibiotic stocks (ampicillin and chloramphenicol), filled to 750 mL with ddH_2_0. We induced expression using 1 mM IPTG and allowed the cultures to grow overnight at 16°C (or 4 hours at 37°C/250 rpm for isotopically labeled samples). Cells were then pelleted at 3,000 rpm for 20 min. After decanting the media, we stored the pellets at -20 °C. To lyse the cells, we resuspended the frozen pellets (3-5 g) in 25 mM Tris, 100 mM NaCl, pH 7.4, incubated for 20 min at 25 °C with DNAse I and lysozyme (ThermoFisher Scientific), and then sonicated on ice. We isolated the soluble fraction by centrifugation at 15,000 rpm for 20-60 min at 4°C.

We purified the proteins using an Äkta PrimePlus FPLC. In its calcium bound form, hS100A9 naturally forms a high-affinity nickel binding site. We exploited this to purify the protein. We brought the soluble lysate to 2 mM CaCl_2_ and 25 mM imidazole and then ran it over a 5 mL HisTrap FF nickel-affinity column. We eluted the protein using a 50 mL gradient from 25-1,000 mM imidazole in 25 mM Tris, 100 mM NaCl, 2 mM CaCl_2_, pH 7.4. We pooled the peak elution fractions and dialyzed overnight against 4 L of 25 mM Tris, 100 mM NaCl, pH 8.0 at 4°C. We then ran the sample over a Q HP anion exchange column and eluted with a gradient of 100-1,000 mM NaCl. We checked for purity on an SDS-PAGE gel. If trace contaminants remained, we performed an additional Q HP anion exchange at pH 6, using the same base buffers and elution strategy as in the previous anion exchange step. All purified proteins were dialyzed overnight at 4 °C into 25 mM Tris, 100 mM NaCl, pH 7.4. We removed divalent ions by placing 2 g/L Chelex-100 resin (Biorad) into the dialysis container. We flash froze the proteins, dropwise, in liquid nitrogen and stored at -80 °C. All proteins had purity >99% as assessed by SDS-PAGE.

### Biophysical and biochemical characterization

For all experiments, we thawed protein aliquots on ice freezer stocks and then either dialyzed in the appropriate experimental buffer overnight at 4°C or exchanged 3x into experimental buffer using 3K microsep spin concentrator columns (Pall Corporation). We filter-sterilized all samples using 0.1 μm spin filters (EMD Millipore. Thawed aliquots were used for no more than one week before discarding. We measured concentration by placing the protein into 6 M guanidinium HCl and measuring absorbance at 280 nm. We used an extinction coefficient of 6,990 M^-1^ cm^-1^, calculated using the ExPASy ProtParam tool.

### CD Spectroscopy and chemical denaturation studies

We performed equilibrium unfolding experiments using 2.5 μM S100A9 dimer, corresponding to 10 μM calcium binding sites. To equilibrate each protein sample fully at each measured urea concentration, we did at least one manual folding/refolding experiment where we pre-equilibrated samples in 0 or 9 M urea, mixed these samples at different ratios, and then allowed the mixtures to equilibrate overnight. We measured our final urea concentrations for each sample using their refractive index. For many variants, we complemented these manual titrations with automated titrations, in which we used an automatic titrator to gradually increase urea. For high calcium concentrations and some of the mutants, equilibration took longer than was feasible for an automated titration; therefore, we relied solely on manually titrations.

We did these experiments in 25 mM spectroscopy-grade Tris, 100 mM NaCl, pH 7.4. Manual unfolding and refolding curves were constructed by making concentrated 100 μM protein stocks in either buffer or 10 M urea at room temperature (25 ℃), and then preparing dilutions at various concentrations of urea—causing the protein to either unfold or fold over equilibration. Samples were left to equilibrate in denaturant overnight at room temperature (>16 hours). We measured the circular dichroism (CD) at 222 nm using a Jasco J-815 CD spectrometer. Unfolding/refolding equilibration was confirmed by comparing unfolded vs. refolded protein CD signal at the same concentration. CD signal was quantified at 222 nm in a 1 mm cuvette using a 1 nm bandwidth, standard sensitivity, and 2 second D.I.T. HT voltage was < 600 V for all measurements.

We fit a two-state unfolding model to the data to extract thermodynamic parameters:^23^

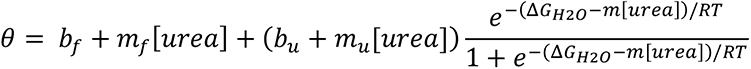

where *θ* is the fraction of molecules folded, *b_f_*, *m_f_*, *b_u_*, and *m_u_* are the folded and unfolded baseline y-intercepts and slopes, *ΔG_H2O_* is the unfolding free energy in water, *m* is the m-value, *R =* 0.001987 J×K^-1^×mol^-1^ and *T =* 298.15 K. We used two fitting strategies to analyze the data. We first allowed all six model parameters to float during regression for each protein/calcium condition. This yielded m-values ranging from 0.5-1.5, often yielding unrealistically curved native baselines (Fig S1B,C, Table S2). To better constrain the fits, we next fit a single m-value across all variant/calcium conditions (Fig S1A). This yielded a global m-value of 0.95 kcal×mol×^-1^×M^-1^, with more consistent and reasonable native baselines for each protein/calcium condition (e.g. Fig S1B,C vs. Fig 2,C). The ΔG values extracted by the two methods were qualitatively similar (Fig S1D), while the C_m_ values almost identical (Fig S1F). The main text and display figures use the global m-value fit. Fits were done using lmfit.^58^

### NMR experiments

Samples were prepared in 25 mM Tris, 100 mM NaCl, 10 mM CaCl_2_, pH 7.4, 10% D_2_O. All protein concentrations ranged from 0.5 – 1.0 mM dimer. All NMR experiments were performed at 25 °C or 37 °C on an 800 MHz (18.8T) Bruker Avance IIIHD spectrometer at Oregon State University equipped with a triple resonance (HCN) cryoprobe. 2D ^15^N-TROSY spectra were collected with 32 transients, 1024 direct points with a signal width of 12820, and 256 indirect points with a signal width of 2837 Hz in ^15^N. Backbone assignments were made based on the wild type assignments (BMRB: 30017)^26^ and confirmed using a suite of 3D experiments including HNCACB, HNCOCACB, HNCACO, and HNCO. Side chain assignments were made using 3D HCCH-TOCSY, 3D CCH-TOCSY, 2D HDCB, 3D HECB. Distance restraints were obtained using a timeshared ^15^N-^13^C-NOESY-HSQC^59^ with mixing time of 160 ms. Three bond HN-HA coupling constants were measured with a 3D HNHA experiment. All 3D experiments were performed with non-uniform sampling percentages between 25 and 50%. All spectra were processed using NMRPipe;^60^ data were visualized using the CCPNMR or nmrviewJ analysis programs.^61^

The NMR structure was determined with Xplor-NIH^62^ using data from Talosn derived dihedral angles, HN-HA 3-bond J-couplings, ^15^N and ^13^C NOE’s, and H-bonds inferred from secondary structure assignments. NOE spectra were picked using automatic peak picking in nmrviewJ and obvious noise peaks removed manually. Preliminary structures were calculated using PASD from the Xplor-NIH software package, with initial probabilities set based on the crystal structure (pdb 1IRJ)^17^. Structures were further refined with the final NOE assignments using Xplor-NIH. Additional distance restraints were included in the refinement stage to define the sidechain interactions with calcium based on the crystal structure.

For hydrogen-deuterium exchange (HDX) experiments by NMR, protein samples were lyophilized and resuspended in 100% D_2_O immediately prior to beginning to measure 2D ^1^H-^15^N TROSY-HSQC spectra. Spectra were collected over 24 hours: every ∼11 minutes for the first hour, every hour up to ∼8 hours, and then a final spectrum after 24 hours. Peak intensities were quantified using Sparky. We placed each residue into one of three classes: exchanged too quickly to quantify, exchanged too slowly to quantify, and exchanged on a timescale we could quantify. To make these classifications, we fit a two-parameter exponential decay model and a linear model with slope of zero to the peak intensity versus time for each residue. We used the Akaike Information Criterion (AIC) to determine select the model that best fit the data. Those residues for which we used the exponential model were quantifiable. Residues for which we used the linear model were classified as either too fast (relative intensity < 0.1) or too slow to quantify. See Fig S3, S4.

### Hydrogen-deuterium exchange mass spectrometry studies

Hydrogen exchange mass spectrometry (HDX-MS) experiments were carried out using a LEAP PAL autosampler (LEAP Technologies, Carrboro, NC, USA). 10 μM protein stocks were maintained at 4°C. 3 μL of 10 μM protein sample was diluted into 27 μL of deuterated buffer at 20°C (90% final deuterium concentration) . After a varying labeling time, 27 μL of labeled sample was quenched with 27 μL of 3.5M GdmCl, 1.5M Glycine pH 2.4 at 1°C. 45 μL of quenched sample was then immediately injected into an Thermo UltiMate 3000 UHPLC system (Buffer A, Buffer B). For inline digestion, samples were flowed over two immobilized protease columns (Upchurch C130B), one with aspergillopepsin and one with porcine pepsin (conjugated to POROS 20 AL Aldehyde Activated Resin, Thermo Scientific), at a flow rate of 100 uL/min of buffer A. Digested protein was then desalted on a trap column (Upchurch C-128 with POROS R2 beads) for 2 minutes at a flow rate of 300 uL/min. An acetonitrile gradient (10% buffer B to 60% buffer B over 9 minutes) eluted peptides from this trap column onto an analytical C-8 column (Thermo 72205– 050565) for separation before injection into an ESI source for mass spectrometry analysis on a Thermo Q Exactive mass spectrometer operating in positive ion mode.

For every protein variant, a non-deuterated control was subjected to the same LC/MS protocol as the deuterated samples and the resulting peptides were identified using tandem MS (MS/MS). Peptide precursor spectra were acquired in data-dependent mode with the top 10 most abundant ions (charge state >= 2, <= 6) selected for fragmentation and product ion analysis. Following MS/MS acquisition precursor ions were excluded from further fragmentation for 4 seconds. The peptide identification software Byonic (Protein Metrics Inc.) was used to identify peptides from the tandem MS data. Following peptide identification deuterium incorporation for each peptide at each labeling time was determined by centroid analysis with the software HDExaminer (Sierra Analytics). Data were not corrected for back exchange. All data shown are comparisons between different solution conditions collected on the same instrument. Previous back exchange controls with this experimental setup had an average of 22% back exchange.^28^

### TLR4 activity assay and cell culture conditions

We purchased commercially distributed HEK293T cells from ATCC (CRL-11268). Because we are using this cell line as a host for heterologous transient transfections, the appropriate control for consistency between assays is the measurement of reporter output for a set of control plasmids and a panel of known treatments. Upon thawing each batch of cells, we ran a positive control for ligand-induced response. We transfected the cells with plasmids encoding human CD14, human MD-2, human TLR4, renilla luciferase behind a constitutive promoter, and firefly luciferase behind an NF-KB promoter. We then characterized the raw luciferase output for five treatments: 1) mock, 2) LPS, 3) LPS + polymyxin B, 4) hS100A9 + polymyxin B, and 5) hS100A9 + 1.25x polymyxin B. LPS is an endotoxin positive control that activates TLR4. Polymixin B sequesters endotoxin present in the environment. This experiment has a stereotypical pattern of responses in renilla luciferase (high for all) and firefly luciferase (low, high, low, high, high). To ensure that the cells maintained their properties between passages, we repeated the mock, LPS, and LPS + polymyxin B control on every single experimental plate. This assay has a built-in control for mycoplasma contamination. A high firefly luciferase signal in the absence of added agonist indicates another source of NF-kappa B output in the cells—most plausibly, contamination. We discarded any cells that exhibit high background values or reached 30 passages. All plasmids, cell culture conditions, and transfections for measuring the activity of S100s against TLR4s were identical to those previously described.^15, 31, 63^ Briefly, human embryonic kidney cells (HEK293T/17, ATCC CRL-11268) were maintained up to 30 passages in Dulbecco’s Modified Eagle Media (DMEM) supplemented with 10% fetal bovine serum (FBS) at 37°C with 5% CO2. Lipopolysaccharide *E. coli* K-12 LPS (LPS - tlrl-eklps, Invivogen) aliquots were prepared at 5 mg/ml in endotoxin-free water and stored at -20°C. Working solutions were prepared at 10 ug/ml and stored at 4°C to avoid freeze-thaw cycles. S100 proteins were prepared by exchanging into endotoxin-free PBS and incubating with an endotoxin removal column (Thermo Fisher Scientific) for 2 hours. S100 LPS contamination was assessed by measuring activity with and without Polymyxin B, an LPS chelating agent. 2 μM (dimer) S100 treatments were prepared by diluting stocks 25:75 endotoxin-free PBS:serum-free Dulbecco’s Modified Eagle Media (DMEM – Thermo Fisher Scientific). Polymyxin B (PB, 200 μg per 100 μL well) was added to limit activity due to background endotoxin contamination. Cells were incubated with treatments for 3 hours prior to assaying activity. We used the Dual-Glo Luciferase Assay System (Promega) to assay Firefly and Renilla luciferase activity of individual wells. Each NF-κB induction value shown represents the Firefly luciferase activity divided by the Renilla luciferase activity, background-subtracted using the LPS + PB activity for each TLR4 species and normalized to the activity of LPS alone for each TLR4 species to normalize between plates. All measurements were performed using three technical replicates per plate, a minimum of five biological replicates total, and a minimum of two separate protein preps.

## Supporting information

Combined supplemental material

## ACKNOWLEDGEMENTS

We would like to thank members of the Harms lab for helpful feedback during preparation of this manuscript. This research was funded by grants from the National Institutes of Health (NIH- 3R01GM117140-03S1, MJH; NIH-T32GM007413, JLH, NIH-GM050945, SM). MJH was a Pew Scholar in the Biomedical Sciences, supported by The Pew Charitable Trusts. The funders had no role in study design, data collection and analysis, decision to publish, or preparation of the manuscript. SM is a Chan Zuckerberg Biohub Investigator.

## AUTHOR CONTRIBUTIONS

JLH performed experiments, analyzed data, wrote, and edited the manuscript. PNR assisted with NMR data collection and solved the NMR structure of the protein. SMC collected data and analyzed the HDX mass spectrometry data and helped write up those results. GDW prepared samples and performed chemical denaturation experiments. SRP generated the sequence alignment used for the bioinformatics analysis. PJC prepared samples and performed chemical denaturation experiments. SM analyzed the HDX mass spectrometry data and helped write up those results. MJH managed the project, analyzed data, and helped write the manuscript. All authors provided input on the manuscript text and figures.

## DATA AVAILABILITY

The NMR structure of hS100A9/M63F has been deposited in the RCSB Protein Data Bank with accession number 7UI5.

