## Supplementary material for "Evolution avoids a pathological stabilizing interaction in the immune protein S100A9": Combined supplemental material

Harman et al.

- Figure S1: Comparison of individual and global m-value fit results.
- Figure S2: Overlaid $^{1}H/^{15}N$ HSQC spectra of hS100A9 and hS100A9/M63F
- Figure S3: Fits to HDX of amide backbone by NMR.
- Figure S4: Difference in amide backbone exchange rate measured for hS100A9/M63F.
- Figure S5: Calcium-dependent coupling between F37 and F63 is seen when unfolding curves are analyzed using either individual m-values or a global m-value.
- Table S1: Thermodynamic parameters extracted from chemical denaturation experiments using global m-value
- Table S2: Thermodynamic parameters extracted from chemical denaturation experiments using individual m-values
- Table S3: H3/H4 angles for published S100 structures
- Table S4: S100A9 peptides observed in the HDX-MS experiments
- Table S5: Residue-specific information from HDX-NMR experiments
- Video S1: Morph between the crystal structure of hS100A9 and the NMR structure of hS100A9/M63F

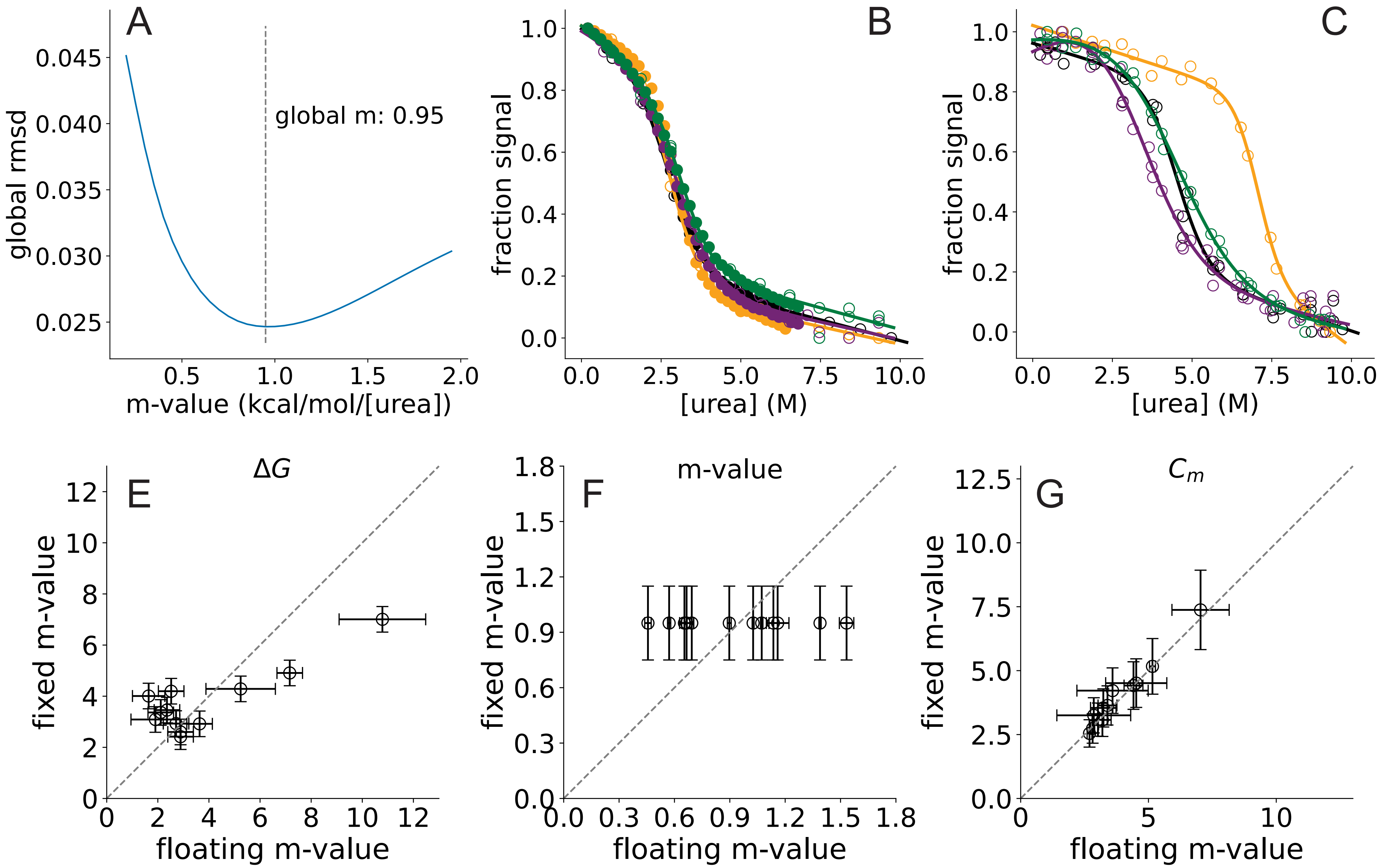


**Figure S1: Comparison of individual and global m-value fit results.** A) Global RMSD vs. m-value for simultaneous fits across all protein variants and calcium conditions. A value of 0.95 kcal/mol/M minimizes the RMSD. B and C) Unfolding curves at 0 mM and 5 mM CaCl2 fit with a floating m-value rather than a global m-value. (These curves correspond to those shown in Fig 5A and B). E) Comparison of estimated ΔG for each protein/calcium conditionfor a fixed m-value versus floating m-values. F) Comparison of the estimated m-value for each protein/calcium conditionfor a fixed m-value versus floating m-values. G) Comparison of the estimated Cm for each protein/calcium condition for a fixed m-value versus floating m-values.

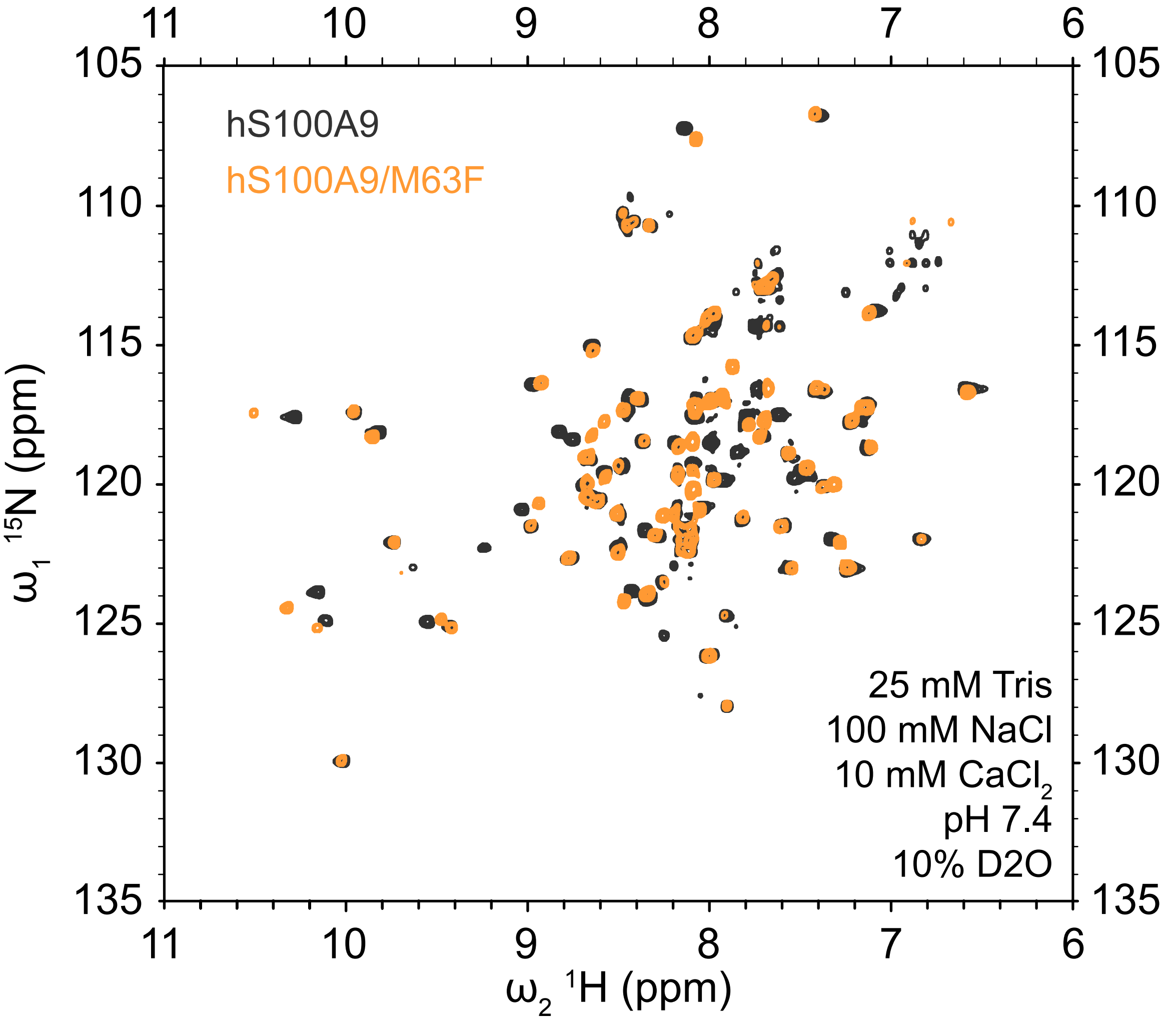


**Figure S2: $^{1}H/^{15}N$ HSQC spectra of hS100A9 and hS100A9/M63F give well resolved peaks.** HSQC spectra for 0.5-1.0 mM hS100A9 (black) and hS100A9/M63F (orange) under conditions indicated on the graph.

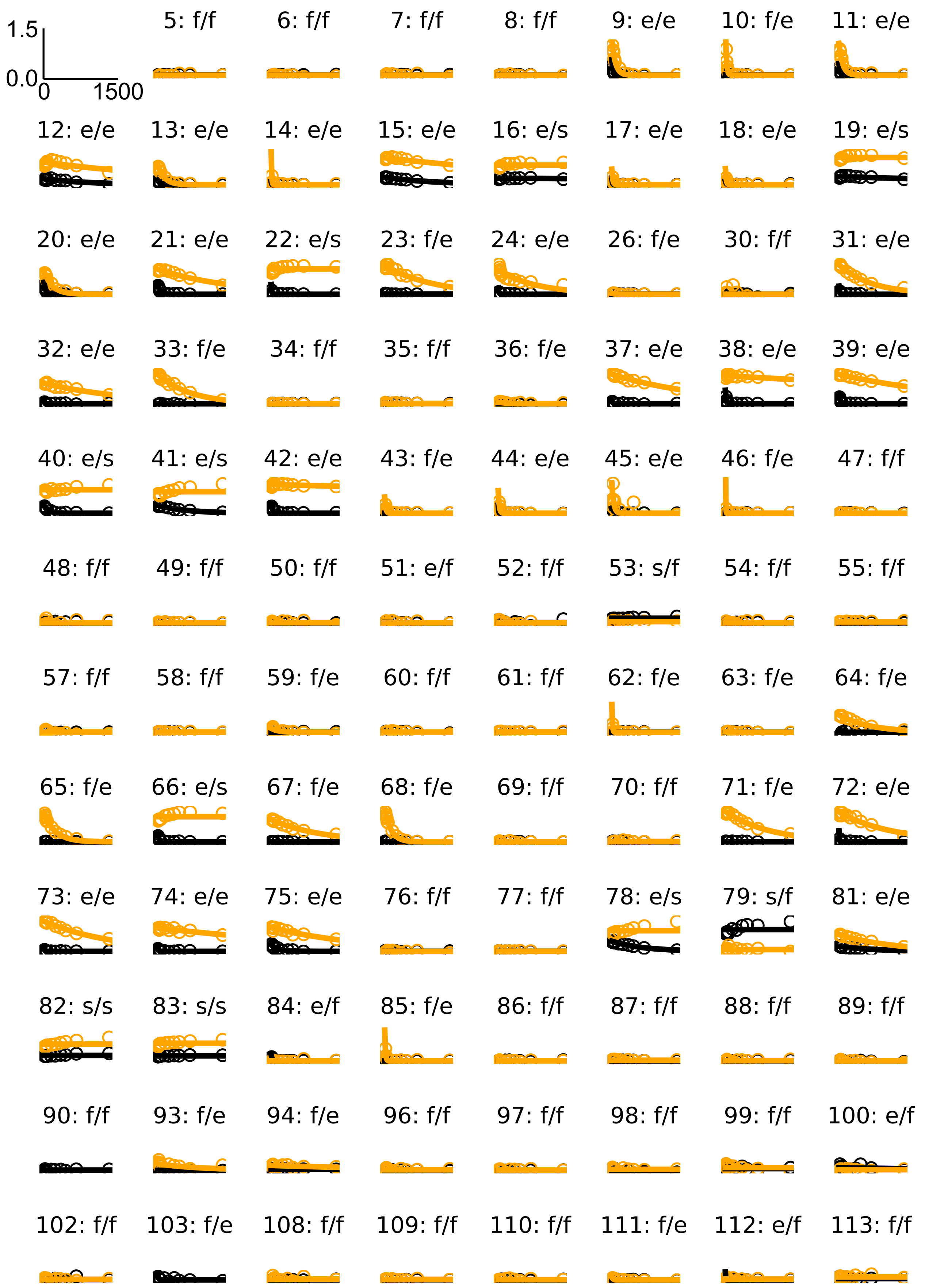


**Figure S3: Individual amide backbone fits.** Each subgraph is a residue (number indicated at top). Fit types are indicated by letter code: “e” (exponential fit), “f” (fast; exchange occurred too quickly for reliable fit); “s”: (slow, exchange occurred too slowly for reliable fit). x-axis is time (0-1500 min); y-axis is the peak intensity relative to intensity of the peak in an unexchanged protein (I/I0). Lines represent best fit to the data. Colors indicate hS100A9 (black) or hS100A9/M63F (orange).

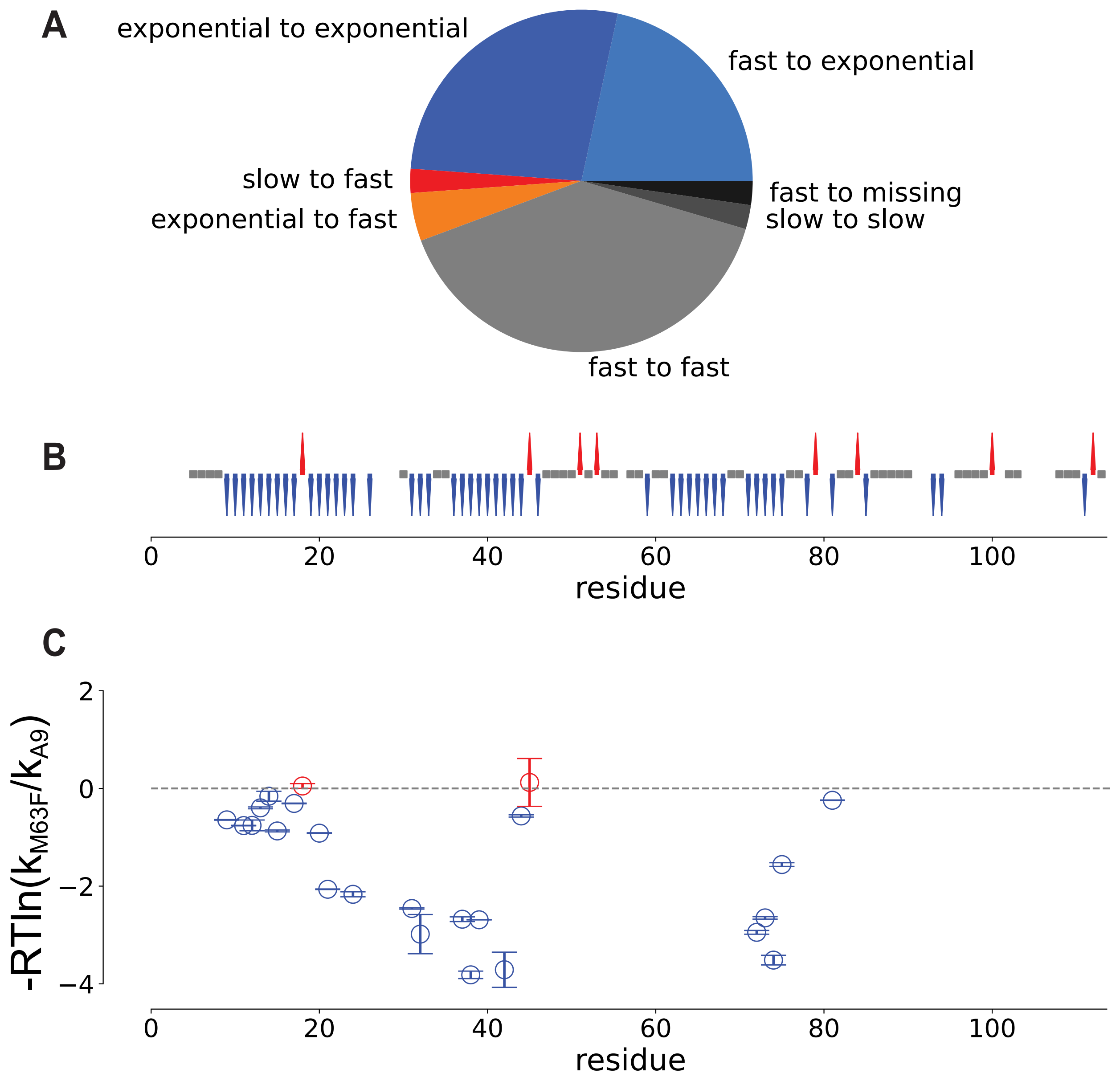


**Figure S4: HDX measured by NMR identifies backbone amides in hS100A9/M63F stabilized relative to hS100A9.** A) Pie chart summarizing fit types for residues under investigation. Text indicates change between hS100A9 and hS100A9/M63F. (For example, "fast to exponential" means the residue exchanged too quickly for a reliable fit in hS100A9, but had a reliable exponential in hS100A9/M63F). B) Qualitative changes in position stability. A blue arrow indicates sites more stable in hS100A9/M63F, either quantitatively ($k_{M63F} < k_{A9}$) or qualitatively (e.g. going from fast to exponential). Red arrows indicate sites more stable in hS100A9 using same criteria. Gray points indicate sites where the relative exchange rate of the sites could not be determined (e.g. both fast, both slow, or one missing). C) Relative rates of exchange for the 25 sites with exponential fits for both hS100A9 and hS100A9/M63F. Error bars are 95% confidence intervals on regression uncertainty. Colors indicate more stable in hS100A9/M63F (blue) or more stable in hS100A9 (red).

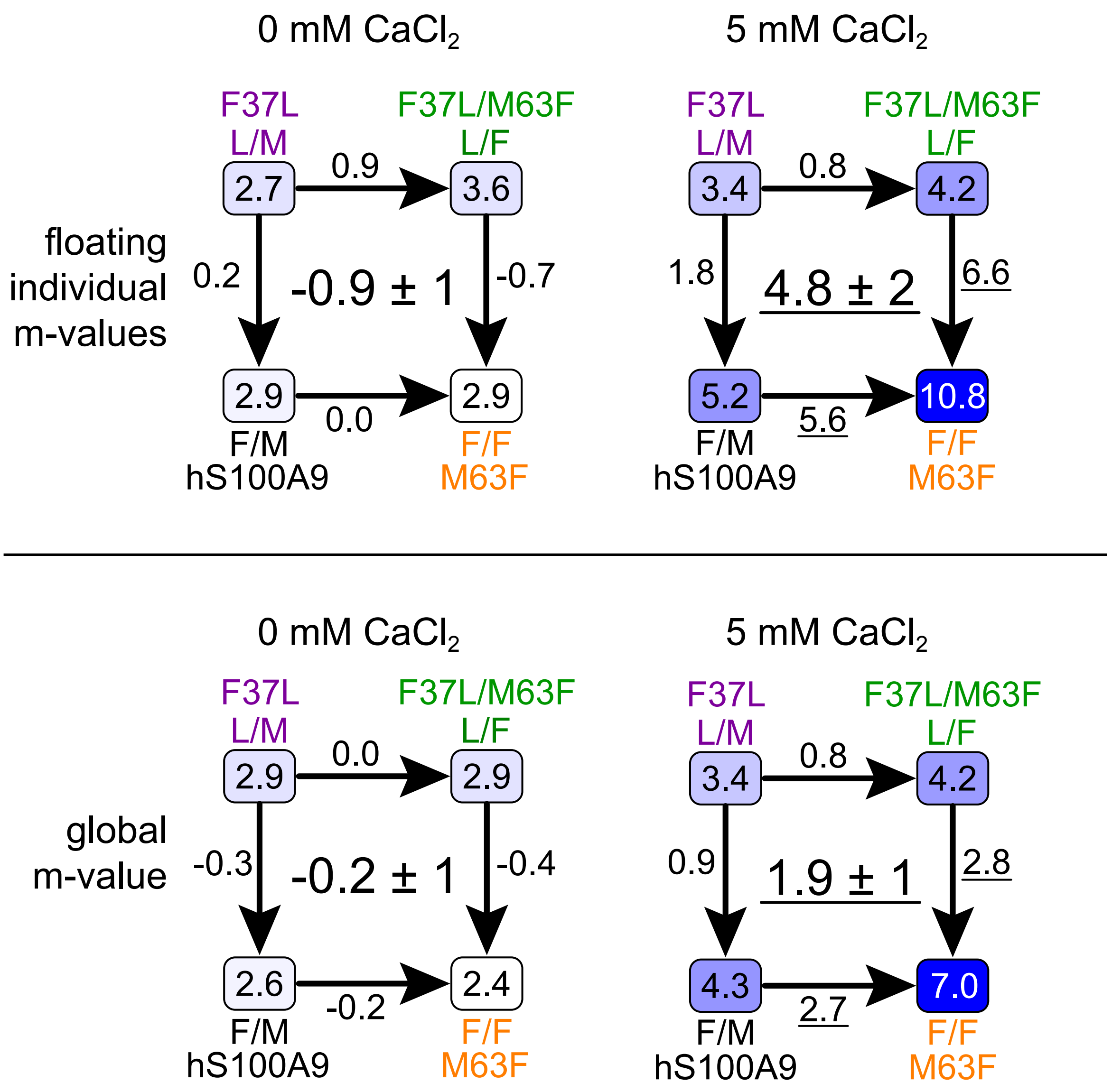


**Figure S5: Calcium-dependent coupling between F37 and F63 is seen when unfolding curves are analyzed using either individual m-values or a global m-value.** Top row: ΔG mutant cycles calculated from fits using individual m-values for reach protein/calcium condition. This corresponds to fits shown in Fig S1C,D, Table S2. Bottom row: ΔG mutant cycles calculated from fits using a global m-value (0.95 kcal/mol/M). This corresponds to fits shown in Fig 5A,B, Table S1).

**Table S1: Thermodynamic parameters extracted from chemical denaturation experiments using global m-value**

Available for download as a csv:
https://github.com/harmslab/stability-constraint-ms-figures/blob/88bf7710fef904b32799b47c6de52db543caeb31/figure_2bc_5ab_s1_s5_table_s1_s2_chemical-denaturations/table_s1.csv

Columns:

- variant: protein
- calcium_conc: calcium concentration in M
- dG: $\Delta G_{H2O}$ maximum likelihood estimate
- dG_err: uncertainty on $\Delta G_{H2O}$ fit parameter
- m: m-value maximum likelihood estimate
- m_err: uncertainty on m-value fit parameter
- Cm: [urea] concentration at which protein is half unfolded (M)
- Cm_err: uncertainty on Cm (propagated from fit uncertainty on $\Delta G_{H2O}$ and m-value)
- rmsd: root mean squared deviation between observed points and fit

**TABLE S1**

| variant | calcium_conc | dG | dG_err | m | m_err | Cm | Cm_err | rmsd |
|---|---|---|---|---|---|---|---|---|
| hS100A9 | 0.0000 | 2.60 | 0.50 | 0.95 | 0.20 | 2.74 | 0.58 | 0.0006 |
| hS100A9 | 0.0005 | 3.46 | 0.50 | 0.95 | 0.20 | 3.64 | 0.77 | 0.0020 |
| hS100A9 | 0.0050 | 4.29 | 0.50 | 0.95 | 0.20 | 4.51 | 0.95 | 0.0028 |
| hS100A9/M63F | 0.0000 | 2.42 | 0.50 | 0.95 | 0.20 | 2.54 | 0.54 | 0.0010 |
| hS100A9/M63F | 0.0005 | 4.91 | 0.50 | 0.95 | 0.20 | 5.17 | 1.09 | 0.0014 |
| hS100A9/M63F | 0.0050 | 7.01 | 0.50 | 0.95 | 0.20 | 7.38 | 1.55 | 0.0023 |
| hS100A9/F37L | 0.0000 | 2.93 | 0.50 | 0.95 | 0.20 | 3.08 | 0.65 | 0.0013 |
| hS100A9/F37L | 0.0005 | 3.09 | 0.50 | 0.95 | 0.20 | 3.25 | 0.69 | 0.0041 |
| hS100A9/F37L | 0.0050 | 3.36 | 0.50 | 0.95 | 0.20 | 3.54 | 0.75 | 0.0028 |
| hS100A9/F37L/M63F | 0.0000 | 2.92 | 0.50 | 0.95 | 0.20 | 3.07 | 0.65 | 0.0015 |
| hS100A9/F37L/M63F | 0.0005 | 4.01 | 0.50 | 0.95 | 0.20 | 4.22 | 0.89 | 0.0029 |
| hS100A9/F37L/M63F | 0.0050 | 4.20 | 0.50 | 0.95 | 0.20 | 4.42 | 0.93 | 0.0019 |

**Table S2: Thermodynamic parameters extracted from chemical denaturation experiments using individual m-values**

Available for download as a csv:
https://github.com/harmslab/stability-constraint-ms-figures/blob/5a4a1071f3500fe8eebf6d01d37f9f5cd71e237e/figure_2bc_5ab_s1_s5_table_s1_s2_chemical-denaturations/table_s2.csv

Columns:

- variant: protein
- calcium_conc: calcium concentration in M
- dG: $\Delta G_{H2O}$ maximum likelihood estimate
- dG_err: uncertainty on $\Delta G_{H2O}$ fit parameter
- m: m-value maximum likelihood estimate
- m_err: uncertainty on m-value fit parameter
- Cm: [urea] concentration at which protein is half unfolded (M)
- Cm_err: uncertainty on Cm (propagated from fit uncertainty on $\Delta G_{H2O}$ and m-value)
- rmsd: root mean squared deviation between observed points and fit

**TABLE S2**

| variant | calcium_conc | dG | dG_err | m | m_err | Cm | Cm_err | rmsd |
|---|---|---|---|---|---|---|---|---|
| hS100A9 | 0.0000 | 2.89 | 0.50 | 1.03 | 0.01 | 2.82 | 0.08 | 0.0006 |
| hS100A9 | 0.0005 | 2.36 | 0.50 | 0.69 | 0.01 | 3.40 | 0.37 | 0.0019 |
| hS100A9 | 0.0050 | 5.24 | 1.36 | 1.16 | 0.06 | 4.52 | 1.20 | 0.0027 |
| hS100A9/M63F | 0.0000 | 2.89 | 0.50 | 1.07 | 0.01 | 2.69 | 0.09 | 0.0010 |
| hS100A9/M63F | 0.0005 | 7.16 | 0.50 | 1.39 | 0.01 | 5.16 | 0.12 | 0.0010 |
| hS100A9/M63F | 0.0050 | 10.79 | 1.69 | 1.53 | 0.04 | 7.04 | 1.12 | 0.0018 |
| hS100A9/F37L | 0.0000 | 2.71 | 0.50 | 0.90 | 0.01 | 3.02 | 0.23 | 0.0013 |
| hS100A9/F37L | 0.0005 | 1.90 | 0.96 | 0.67 | 0.04 | 2.86 | 1.44 | 0.0040 |
| hS100A9/F37L | 0.0050 | 2.11 | 0.50 | 0.65 | 0.01 | 3.23 | 0.50 | 0.0027 |
| hS100A9/F37L/M63F | 0.0000 | 3.63 | 0.50 | 1.14 | 0.02 | 3.20 | 0.34 | 0.0015 |
| hS100A9/F37L/M63F | 0.0005 | 1.64 | 0.63 | 0.46 | 0.02 | 3.59 | 1.39 | 0.0025 |
| hS100A9/F37L/M63F | 0.0050 | 2.52 | 0.50 | 0.57 | 0.01 | 4.41 | 0.37 | 0.0016 |

**Table S3: H3/H4 angles for published S100 structures**

Available for download as a csv:
https://github.com/harmslab/stability-constraint-ms-figures/blob/5a4a1071f3500fe8eebf6d01d37f9f5cd71e237e/figure_2a_table_s3_s100-structures/calc-h3-h4-angles/table-s2_s100-h3-h4-angles.csv

Columns:

- pdb_id: RCSB PDB ID for structure analyzed
- S100: S100 paralog of structure
- calcium_bound: whether or not structure has calcium bound to S100
- target_bound: whether or not the structure has a peptide target bound to the S100
- h3_span: residues spanning helix 3
- h4_span: residues spanning helix 4
- h3_h4 angle: angle between helices 3 and 4

**TABLE S3**

| pdb_id | S100 | calcium_bound | target_bound | h3_span | h4_span | h3_h4 angle |
|---|---|---|---|---|---|---|
| 2wcf | S100A12 | FALSE | FALSE | (52, 63) | (70, 85) | 25.41 |
| 1k8u | S100A6 | FALSE | FALSE | (49, 62) | (68, 82) | 14.23 |
| 2wce | S100A12 | FALSE | FALSE | (50, 63) | (70, 85) | 19.44 |
| 2rgi | S100A2 | FALSE | FALSE | (51, 64) | (70, 84) | 15.86 |
| 2wcb | S100A12 | FALSE | FALSE | (50, 58) | (70, 90) | 14.77 |
| 1k9p | S100A6 | FALSE | FALSE | (49, 62) | (68, 81) | 13.94 |
| 2wc8 | S100A12 | FALSE | FALSE | (47, 58) | (70, 90) | 13.14 |
| 1qls | S100A11 | TRUE | TRUE | (51, 61) | (70, 92) | 45.34 |
| 3iqq | S100B | TRUE | TRUE | (50, 61) | (70, 84) | 73.38 |
| 4eto | S100A4 | TRUE | TRUE | (50, 61) | (70, 90) | 69.12 |
| 3cr5 | S100B | TRUE | FALSE | (50, 61) | (70, 89) | 77.90 |
| 3cr4 | S100B | TRUE | FALSE | (50, 61) | (70, 87) | 75.40 |
| 2q91 | S100A4 | TRUE | FALSE | (50, 61) | (70, 88) | 76.13 |
| 1e8a | S100A12 | TRUE | FALSE | (49, 60) | (72, 85) | 62.24 |
| 3gk1 | S100B | TRUE | FALSE | (49, 60) | (69, 86) | 83.00 |
| 3d0y | S100B | TRUE | FALSE | (50, 61) | (70, 89) | 74.95 |
| 1k9k | S100A6 | TRUE | FALSE | (48, 59) | (68, 88) | 58.80 |
| 3psr | S100A7 | TRUE | FALSE | (39, 49) | (70, 90) | 122.03 |
| 2wos | S100A7 | TRUE | FALSE | (39, 49) | (70, 90) | 124.21 |
| 2wor | S100A7 | TRUE | FALSE | (39, 49) | (70, 90) | 124.08 |
| 1mho | S100B | TRUE | FALSE | (49, 60) | (69, 87) | 80.21 |
| 3cr2 | S100B | TRUE | FALSE | (50, 61) | (70, 88) | 75.68 |
| 3iqo | S100B | TRUE | FALSE | (49, 60) | (69, 86) | 81.56 |
| 3hcm | S100B | TRUE | FALSE | (50, 61) | (70, 88) | 72.97 |
| 2h2k | S100A13 | TRUE | FALSE | (53, 61) | (70, 89) | 34.23 |
| 3cga | S100A4 | TRUE | FALSE | (48, 59) | (68, 86) | 52.71 |
| 1j55 | S100P | TRUE | FALSE | (46, 55) | (64, 86) | 64.25 |
| 1mr8 | S100A8 | TRUE | FALSE | (50, 58) | (70, 87) | 65.92 |
| 1irj | S100A9 | TRUE | FALSE | (56, 66) | (76, 86) | 68.08 |
| 2psr | S100A7 | TRUE | FALSE | (39, 49) | (70, 90) | 123.68 |
| 3gk2 | S100B | TRUE | FALSE | (49, 60) | (69, 86) | 82.79 |
| 2wnd | S100A7 | TRUE | FALSE | (39, 49) | (70, 90) | 121.62 |
| 3gk4 | S100B | TRUE | FALSE | (50, 61) | (70, 88) | 77.55 |
| 1k96 | S100A6 | TRUE | FALSE | (48, 59) | (68, 88) | 59.29 |
| 3d10 | S100B | TRUE | FALSE | (50, 61) | (70, 89) | 73.67 |

**Table S4: S100A9 peptides observed in the HDX-MS experiments**

Available for download as a csv:
https://github.com/harmslab/stability-constraint-ms-figures/blob/bfefe15a01e7a4ce0eb87b14a24f7f636e5448f8/figure_4ab_5cd_table_s4_hdx/table_s4_deterium-uptake-data.csv

Columns:

- Protein State: WT (hS100A9), M63F, F37L, or F37L/M63F
- Start: first residue in peptide:
- End: last residue in peptide
- Sequence: peptide sequence
- Deut Time (sec): how long deuteration ran before being quenched
- maxD: maximum number of deuterons possible for this peptide
- #D: number of deuterons observed
- %D: percent deuteration (#D/maxD)
- Stddev: estimated standard deviation on %D

**TABLE S4**

Table too large to display (~150 pages). Accessible via link.

https://github.com/harmslab/stability-constraint-ms-figures/blob/bfefe15a01e7a4ce0eb87b14a24f7f636e5448f8/figure_4ab_5cd_table_s4_hdx/table_s4_deterium-uptake-data.csv

**Table S5: Residue-specific information from HDX-NMR experiments**

Available for download as a csv:
https://github.com/harmslab/stability-constraint-ms-figures/blob/58c6ba864e96e9935167df70b70209980a8d510c/figure_4c_s3_s4_table_s5/table_s5.csv

Columns:

- residue: residue number
- qualitative: whether site is *unchanged* between hS100A9/M63F and hS100A9; *slower* in hS100A9/M63F than hS100A9; *faster* in hS100A9/M63F than hS100A9
- quantitative: $\ln(k_{ex,M63F}) - \ln(k_{ex,hA9})$ for exponential fits
- quant_err: 95% confidence interval propagated from regression uncertainty for value shown in quantitative column
- a9: fit type for hS100A9 experiment (fast, slow, exponential, missing)
- m63f: fit type for hS100A9/M63F experiment (fast, slow, exponential, missing)

**TABLE S5**

| residue | qualitative | quantitative | quant_err | a9 | m63f |
|---|---|---|---|---|---|
| 5 | unchanged | | | fast | fast |
| 6 | unchanged | | | fast | fast |
| 7 | unchanged | | | fast | fast |
| 8 | unchanged | | | fast | fast |
| 9 | slower | -1.0866092 | 0.01058853 | exponential | exponential |
| 10 | slower | | | fast | exponential |
| 11 | slower | -1.2838538 | 0.00882666 | exponential | exponential |
| 12 | slower | -1.2779697 | 0.18651498 | exponential | exponential |
| 13 | slower | -0.6715392 | 0.03179398 | exponential | exponential |
| 14 | slower | -0.2679822 | 0.17022828 | exponential | exponential |
| 15 | slower | -1.4699329 | 0.03267487 | exponential | exponential |
| 16 | slower | | | exponential | slow |
| 17 | slower | -0.5189272 | 0.01189698 | exponential | exponential |
| 18 | faster | 0.08382047 | 0.08164391 | exponential | exponential |
| 19 | slower | | | exponential | slow |
| 20 | slower | -1.5460689 | 0.01455658 | exponential | exponential |
| 21 | slower | -3.4857198 | 0.01074961 | exponential | exponential |
| 22 | slower | | | exponential | slow |
| 23 | slower | | | fast | exponential |
| 24 | slower | -3.6606302 | 0.08661899 | exponential | exponential |
| 26 | slower | | | fast | exponential |
| 30 | unchanged | | | fast | fast |
| 31 | slower | -4.1510996 | 0.02165643 | exponential | exponential |
| 32 | slower | -5.0372961 | 0.67804881 | exponential | exponential |
| 33 | slower | | | fast | exponential |
| 34 | unchanged | | | fast | fast |
| 35 | unchanged | | | fast | fast |
| 36 | slower | | | fast | exponential |
| 37 | slower | -4.5214009 | 0.08138629 | exponential | exponential |
| 38 | slower | -6.4446757 | 0.12925411 | exponential | exponential |
| 39 | slower | -4.5407726 | 0.00358419 | exponential | exponential |
| 40 | slower | | | exponential | slow |
| 41 | slower | | | exponential | slow |
| 42 | slower | -6.2680664 | 0.60822258 | exponential | exponential |
| 43 | slower | | | fast | exponential |
| 44 | slower | -0.9509807 | 0.03862213 | exponential | exponential |
| 45 | faster | 0.20787115 | 0.82697167 | exponential | exponential |
| 46 | slower | | | fast | exponential |
| 47 | unchanged | | | fast | fast |
| 48 | unchanged | | | fast | fast |
| 49 | unchanged | | | fast | fast |
| 50 | unchanged | | | fast | fast |

| | | | | | |
|---|---|---|---|---|---|
| 51 | faster | | | exponential | fast |
| 52 | unchanged | | | fast | fast |
| 53 | faster | | | slow | fast |
| 54 | unchanged | | | fast | fast |
| 55 | unchanged | | | fast | fast |
| 57 | unchanged | | | fast | fast |
| 58 | unchanged | | | fast | fast |
| 59 | slower | | | fast | exponential |
| 60 | unchanged | | | fast | fast |
| 61 | unchanged | | | fast | fast |
| 62 | slower | | | fast | exponential |
| 63 | slower | | | fast | exponential |
| 64 | slower | | | fast | exponential |
| 65 | slower | | | fast | exponential |
| 66 | slower | | | exponential | slow |
| 67 | slower | | | fast | exponential |
| 68 | slower | | | fast | exponential |
| 69 | unchanged | | | fast | fast |
| 70 | unchanged | | | fast | fast |
| 71 | slower | | | fast | exponential |
| 72 | slower | -4.9745863 | 0.06492935 | exponential | exponential |
| 73 | slower | -4.4756507 | 0.04229276 | exponential | exponential |
| 74 | slower | -5.9352058 | 0.1670115 | exponential | exponential |
| 75 | slower | -2.6314128 | 0.06436495 | exponential | exponential |
| 76 | unchanged | | | fast | fast |
| 77 | unchanged | | | fast | fast |
| 78 | slower | | | exponential | slow |
| 79 | faster | | | slow | fast |
| 81 | slower | -0.4100326 | 0.0105906 | exponential | exponential |
| 82 | unchanged | | | slow | slow |
| 83 | unchanged | | | slow | slow |
| 84 | faster | | | exponential | fast |
| 85 | slower | | | fast | exponential |
| 86 | unchanged | | | fast | fast |
| 87 | unchanged | | | fast | fast |
| 88 | unchanged | | | fast | fast |
| 89 | unchanged | | | fast | fast |
| 90 | unchanged | | | fast | missing |
| 93 | slower | | | fast | exponential |
| 94 | slower | | | fast | exponential |
| 96 | unchanged | | | fast | fast |
| 97 | unchanged | | | fast | fast |
| 98 | unchanged | | | fast | fast |
| 99 | unchanged | | | fast | fast |
| 100 | faster | | | exponential | fast |

| | | | |
|---|---|---|---|
| 102 | unchanged | fast | fast |
| 103 | unchanged | fast | missing |
| 108 | unchanged | fast | fast |
| 109 | unchanged | fast | fast |
| 110 | unchanged | fast | fast |
| 111 | slower | fast | exponential |
| 112 | faster | exponential | fast |
| 113 | unchanged | fast | fast |

**Table S6: S100 alignment**

Available for download as a csv:
https://github.com/harmslab/stability-constraint-ms-figures/blob/58c6ba864e96e9935167df70b70209980a8d510c/figure_6abc_table_s6/table_s6_site37-63-analysis.csv

Columns:

- paralog: paralog assignment for sequence
- site_37: state of site 37 (one letter amino acid code)
- site_63: state of site 63 (one letter amino acid code)
- class_site_37: amino acid classification for site 37
- class_site_63: amino acid classification for site 63
- sequence: aligned sequence of paralog

**TABLE S6**

| name | paralog | site_37 | site_63 | class_site_37 | class_site_63 | sequence |
|---|---|---|---|---|---|---|
| S100A8\|Chelonia mydas\|XP_027685838.2 | S100A8 | F | F | aromatic | aromatic | ---------------------------------------------------------------------------------------------------------MTTKLEKALNDIIEVYHEYS-ESECN-DDYLQKDEFKKLLKKEANSFLCYTVPRPGDEEKYIDNLFQKANKDKNEHLHFEEFVTTLAKLAIHAHNMSH |
| S100A8\|Chelydra serpentina\|KAG6923792.1 | S100A8 | F | F | aromatic | aromatic | ---------------------------------------------------------------------------------------------------------MTSKLEKALDDVIEVYHKYS-EKKPDSDDFLQKHEFKQLVKQHAKSFLKCTLPRGKTEDEYIEDLFKKADKNRNGFLHFEEFGTVLASLAIHAHNSSH |
| MRP-126\|Trachemys scripta elegans\|XP_034648305.1 | MRP-126 | F | F | aromatic | aromatic | ---------------------------------------------------------------------------------------------------------PTSKLEAALNDIIDVYHEYS-EKRPNSDDFLQKHEFKQLLKQHAKSFLACTIPRGKTEDEYIEDLFKKADKNKNGQLHFEEFVTTLAKLAIHAHDLSH |
| MRP-126\|Terrapene carolina triunguis\|XP_024055775.1 | MRP-126 | F | F | aromatic | aromatic | ---------------------------------------------------------------------------------------------------------MTSKLEKALNEVIEVYHKYS-EKKPNSDDFLQKHEFKQLLKQHAKSFLECTLPRGKTEDEYIEDLFKKADKNKNGHLHFEEFVTTLGKLAIHAHDISH |
| MRP-126\|Mauremys reevesii\|XP_039368194.1 | MRP-126 | F | F | aromatic | aromatic | ---------------------------------------------------------------------------------------------------------MTSKLENALNDVIEVYHQYS-EKRPNPDDFLQKHEFKQLLKQNAKSFLKCTLPHGSTEDKYIEDLFKKADKNGNGYLHFEEFVTTLAKLAIHAHNISH |
| S100A8\|Gopherus evgoodei\|XP_030398022.1 | S100A8 | F | F | aromatic | aromatic | ---------------------------------------------------------------------------------------------------------MTSKLEKALDDMIDVYHKYS-EKRPNSDDYLQKHEFRQLLKQNAKSFLKCTLPYGQTEDEYIDDLFKKADRNRNEYLHFEEFVTTLAKLAIHAHDISH |
| MRP-126\|Gavialis gangeticus\|XP_019379412.1 | MRP-126 | F | F | aromatic | aromatic | ----------------------------------------------------------------------------------------------------------KTQLEKELDCIVNIYHQYS-FRHPM-DDYLQKAEFKKLLREQASDFLKHTKPPNVTMEEYLDNLFKKADVNRDGRVKFTEFLITLASVAIDVHNTSH |
| MRP-126\|Empidonax traillii\|XP_027766643.1 | MRP-126 | F | F | aromatic | aromatic | ---------------------------------------------------------------------------------------------------------MKTDLELALECAVNIYHQYA-VQYPI-DDYLSKGEFSKLLKETAQPFLHNTKPPKASTDEYIAQLFAKADGNRDGRLKFTEFLTTLCLVAIDAHNRSH |
| MRP-126\|Molothrus ater\|XP_036253853.1 | MRP-126 | F | F | aromatic | aromatic | ---------------------------------------------------------------------------------------------------------MKTDLELALECAVNVYHRFA-AQRPV-DDYLSLGEFSRLLKETAEPFLKNTVPPKTTMDSYIKQLFTKADANHDGRLKFTEFLTTLSLVAIDAHNRSH |
| S100A8\|Catharus ustulatus\|XP_032938532.1 | S100A8 | F | F | aromatic | aromatic | ---------------------------------------------------------------------------------------------------------MKTDLELALECAVNVYHRYA-MRRPM-DDYLSRSEFSKLLRDTAEPFLRRTAPPNTNIDQYITQLFSKADANRDGRLKFTEFLSTLSLVAIDAHNRSH |
| S100A8\|Ficedula albicollis\|XP_016159506.1 | S100A8 | F | F | aromatic | aromatic | ---------------------------------------------------------------------------------------------------------MRTDLELALECAVNVYHRYA-LQRPL-DDFLSRSEFSKLLKEEAEPFLRNTAPPNTSTDTYINQLFTKADANRDGRLKFTEFLSTLSLVAIDAHNRSH |
| S100A8\|Pseudopodoces humilis\|XP_005533718.1 | S100A8 | F | F | aromatic | aromatic | ---------------------------------------------------------------------------------------------------------MKTDLELALECAINVYHRYA-ARRPI-DDYLSKNEFSKLLKETAEPFLRNTAPPNTSTDTYISQLFAKADANHDGRLKFTEFLTTLSLVAIDAHNRSH |
| S100A8\|Hirundo rustica\|XP_039943913.1 | S100A8 | F | F | aromatic | aromatic | ---------------------------------------------------------------------------------------------------------MKTDLELALECAVNVYHRYA-VQRPI-DDYLSRREFSKLLKETAEPFLRNTVPPNTSTDSYIAQLFAKADANRDGRLKFTEFLTTLSLVAIDAHNRFH |
| S100A8\|Lonchura striata domestica\|XP_021397802.1 | S100A8 | F | F | aromatic | aromatic | ---------------------------------------------------------------------------------------------------------MKTNLELALECVVNVYHRYA-MRHPM-DDYLSKNEFSKLLKETAEPFLKNTVPPNTGTDSYIQQLFTKADANHDGRLKFTEFLTTLSLVAIDAHNRSH |
| MRP-126\|Nipponia nippon\|XP_009469673.1 | MRP-126 | F | F | aromatic | aromatic | ---------------------------------------------------------------------------------------------------------MKTDLELALECVVNIYHQYA-IKYPI-DDYLSKTEFSELLKETAKPFLHNTMPPNTSVDDYISKLFTRADRNHDGRLKFTEFLTTLNLVAIDAHNRSH |
| MRP-126\|Anolis carolinensis\|XP_008123593.1 | MRP-126 | F | F | aromatic | aromatic | ---------------------------------------------------------------------------------------------------------MKTDMEKAMACVVDIFHQYC-ILSPI-DDYLQRAEFDKLMKEQAQPFLKKTQPPNLDQKAYLDQLFKKADKNHDAKLKFTEWATVLGLAMNEAHDRSH |
| MRP-126\|Gekko japonicus\|XP_015269604.1 | MRP-126 | F | F | aromatic | aromatic | ---------------------------------------------------------------------------------------------------------MKTKMEQAMECVVNVYHQYC-ILDPV-DDYLHPKEFQKLMKEQAQPFLKNTMPPSDDQNSYIEKLFQQADKDKNGYLKFTEWLYVIGKTLNEAHERSH |
| MRP-126\|Varanus komodoensis\|KAF7235498.1 | MRP-126 | F | F | aromatic | aromatic | ---------------------------------------------------------------------------------------------------------MKTHLEQAMECVVNVYHQYC-ILDPV-DDYLHLKEFSKLMKEQAQPFLKSTTPPNLDQDGYIKQLFEKADKDKTGYLKFTEFFCVLGLALNEGHDRSH |
| S100A8\|Notamacropus eugenii\|ACG69795.1 | S100A8 | F | F | aromatic | aromatic | -----------------------------------------------------------------------------------------------------MASTKYTPSEQDLNKIIDVYHKYS-RKAPD-DDALERKEFEGLIKQQAPEFSQ----YLQDHQISLDSVFKKADLNHDKHIVFPEFVTVLAQIAIEMHNHFH |
| MRP-126\|Trichosurus vulpecula\|XP_036610110.1 | MRP-126 | L | F | hphobic | aromatic | -----------------------------------------------------------------------------------------------------MASTKYTSSEENLNQIIAVFHEYS-HKSEE-EDALEFSELKGLISKEAPGFT----EHLKKENISVESVFKKADLDRDGHMFFPEFLLVLSLMGIEIHNKYH |
| MRP-126\|Phascolarctos cinereus\|XP_020847640.1 | MRP-126 | F | F | aromatic | aromatic | -----------------------------------------------------------------------------------------------------MASTKYTASEEQLNHIVEVFHKYS-RKSGE-EDALELKEFEGLIKEQAPEFF----EYLKK-NTSLASVFKKADLDNDKHMFFSEFVFVLALIAIEIHDKYH |
| S100A8\|Ornithorhynchus anatinus\|XP_028910543.1 | S100A8 | F | Y | aromatic | aromatic | -------------------------------------------------------------------------------------------------MSAAKVIQENTPAEKEMFVIIDVYHKYS-EKGGD-RDGLEKAEFEEMVIQEAPHFYK----KVVKNRDALNKIYKEANVDGDKKLCFKEFVMVLSRISIEAHDRSH |
| MRP-126\|Tachyglossus aculeatus\|XP_038624407.1 | MRP-126 | F | Y | aromatic | aromatic | -------------------------------------------------------------------------------------------------MSASRVSQVKTPTEKAMFDIIDVFHKFS-ERGGY-EDGLEKKEFEDMIHQEAPHFYK-----VVKTHENLEKIYKKANVDGNKWLCFKEVVMVLSSIANDSHNRSH |
| MRP-126\|Pithecopus nordestinus\|AFY11403.1 | MRP-126 | L | F | hphobic | aromatic | ---------------------------------------------------------------------------------------------------------MEIPIQDLMISMISAFNKYA-SGDGD-NNTLNKKELVNMACKEFPSLEK----HPQKD-KFLNDIFKITDRDGDNKINFEEFMTLMTCMTIALRKSGT |
| MRP-126\|Acanthopagrus latus\|XP_036949543.1 | MRP-126 | L | F | hphobic | aromatic | ----------------------------------------------------------------------------------------------------------HHDLKGALKTVKDFFYKYA-NQDGD-AHSITKKELKVLFQQELKMEL-----EGCEGKEKLDNIFKDLDDNKDNVVDFNEFCDLFVMILKISHFWLM- |
| MRP-126\|Struthio camelus australis\|XP_009684588.1 | MRP-126 | L | Y | hphobic | aromatic | ----------------------------------------------------------------------------------------------------------MSNLETAMMT-IAVFDKYA-KTQGN-PQTLTKAELKTLLEKELPNFL----SLNIENRTQKRHVYAKSQVNPEPPKACKH-AGQSKSLQLPVRSNFFA |
| MRP-126\|Phalacrocorax carbo\|XP_009503417.1 | MRP-126 | L | I | hphobic | hphobic | -------------------------------------------------------------------------------------------------------------MEHLLDNILKYFTQYA-LLEGS-TDTLSIGGLKNLLQNEFARYLK--------DAFSLENIIKSLDKNNDGRISLDEYVMLIKKITGTG----- |
| MRP-126\|Oryzias javanicus\|RVE62681.1 | MRP-126 | L | M | hphobic | hphobic | ---------------------------------------------------------------------------------------------------------AMARLDGAITNIVDVFMDYA-DDGGK-KRQLNKEELKKLFAQEIPE-LK-----DKISADDIEEAMEMLDKNHDGEVNFREFCRCVCVLATGYHKKIG |
| MRP-126\|Sebastes umbrosus\|XP_037639824.1 | MRP-126 | L | M | hphobic | hphobic | ----------------------------------------------------------------------------------------------------------MARLDLVITNIVDVFMDYA-DDEGK-KQQLNKEELKKMLEQEIQSPE------LKGHADDIGEAMEMLDKNHDGEVNFREFCRCVSVLAKCYHKKIG |
| S100A8\|Maylandia zebra\|XP_024660065.1 | S100A8 | L | L | hphobic | hphobic | --------------------------------------------------------------------------------------------------------TQMSKLEEAMEEVLGIFKQEC----TR-SKTMMKKDLKRTLRAELVNWLQ-----NCKDKTERDNILRELDENPHDCVNFEEFVYYVTRLMMCGHYFLQ |
| MRP-126\|Paroedura picta\|GCF56323.1 | MRP-126 | L | F | hphobic | aromatic | --------------------------------------------------------------------------------------------------------MPQTEMEQSIQTILKLFHESA-AQEGD-HDTLNKAELRGFINKNFPNFLK-----KLKDPIILENVFEQLDTNKDQEISFGEIMQFVTIVAILSHDRIH |
| MRP-126\|Polypterus senegalus\|XP_039608194.1 | MRP-126 | L | F | hphobic | aromatic | -------------------------------------------------------------------------------------------------------MAGQSDLQTAILSILNVFNKYA-KQEGS-KDKLSQSELKKLIEAELGPLSK-----KI-DASKVDELFKEMDKNHDGQVSLCEFCKFVSILTVGYHKSQC |
| MRP-126\|Anolis carolinensis\|XP_003229666.1 | MRP-126 | F | F | aromatic | aromatic | --------------------------------------------------------------------------------------------------------MTKTKMETVADNIIDMFHNYS-SKDSE-KDTLDPVEFKKMIEVELPHFVK-Q---VR-DPKALENFFAKVDGNKNKHINFEEFMKLFGEFVVLSHEKFH |
| MRP-126\|Podarcis muralis\|XP_028566962.1 | MRP-126 | F | F | aromatic | aromatic | ------------------------------------------------------------------------------------------------------MSKSKTQMQVLADGLIELFHKYS----GK-SDTIDKKEFKKMINEQFPDCVE---YP--QNQQGKDKLFKDLDTNKDNELSFEEWTRLVGSFLSCAHVKFH |
| MRP-126\|Zootoca vivipara\|XP_034954781.1 | MRP-126 | F | F | aromatic | aromatic | ------------------------------------------------------------------------------------------------------MSKTKTQMQILADQLIAVFHEYA----GK-SDTIDKKEFKKMINEQFPDCVE---YP--QNQQGKDKLFKDLDLNKDNQLSFEEWTRLLGSFLSCAHIKFH |
| S100A8\|Sarcophilus harrisii\|XP_003768081.1 | S100A8 | F | L | aromatic | hphobic | ---------------------------------------------------------------------------------------------------------MVTKLEGAVNCLVTVFHKYS-LASGH-YHTLSKDKFRKLLETEIPEVLK---NP--KDPKTVEKILKEVDSNKDGMINFEEFLFLATRLLVDAHDATH |
| S100A8\|Vombatus ursinus\|XP_027727273.1 | S100A8 | F | L | aromatic | hphobic | ---------------------------------------------------------------------------------------------------------MVTKFEGAINCMVDVFHKYS-LEHGH-PHAMNRKAFRKLIETECPELLT---NP--KDPKTVETLLKEVDSNKDDLMNFEEYLFMITRLMVDAHEASH |
| S100A8\|Phascolarctos cinereus\|XP_020847078.1 | S100A8 | F | L | aromatic | hphobic | ---------------------------------------------------------------------------------------------------------MVTKFEGAINCMVTVFHKYS-LGHGH-PHAMNRKSFRKLVETECPELLT---NP--KDPKTLEALLKEVDSNKDELMSFEEYLFLITRLMVDAHEASH |
| MRP-126\|Tachyglossus aculeatus\|XP_038624927.1 | MRP-126 | L | F | hphobic | aromatic | ----------------------------------------------------------------------------------------------------------MTVLEKAIDSLIDVFHKYS-LTEGH-YHSISKGVLKTLMEKELTTFLK-----SQKNPPTVDTLFKELDSNKDQQINFEEYMVLVTRVGVACHEEIH |
| S100A8\|Monodelphis domestica\|XP_001372154.1 | S100A8 | F | M | aromatic | hphobic | ---------------------------------------------------------------------------------------------------------MATKLECAINCLVEVFHKYS-LTGGH-PHALSREQFGKLLEKECSEFTK-------KSKKTVPEFMKELDINQDGFINFEEFLILTLKMVIEHHEDSH |
| S100A8\|Octodon degus\|XP_004629976.1 | S100A8 | L | F | hphobic | aromatic | ---------------------------------------------------------------------------------------------------------MMTELETALEDIINVYHKYS-IGVGH-YDVLYRGDLKRLLTNECPQYTK---------KRSADDWFKLLDVNEDGAVTFQEFLVLVTKLLIEEHKREP |
| S100A8\|Choloepus didactylus\|XP_037670203.1 | S100A8 | F | F | aromatic | aromatic | ---------------------------------------------------------------------------------------------------------MPTDLECALNNIIEVFHRYS-LVKGN-YHSLEKDDFKKLLENECRSFTK---------QKSVEEWFKELDINSDDAINFEEFLIMLVKMSVEAHKDTH |
| S100A8\|Erinaceus europaeus\|XP_007525454.1 | S100A8 | L | F | hphobic | aromatic | ---------------------------------------------------------------------------------------------------------MLTNLEKAINNIIDIFHKYS-LKTGN-YHSLYRNDLKSLMESECSMYLK---------EKNADTWFTELDINSDQAINFEEFLILVIKIAVAAHKASH |
| S100A8\|Meriones unguiculatus\|XP_021520244.1 | S100A8 | L | F | hphobic | aromatic | ---------------------------------------------------------------------------------------------------------MLSELEKAMANMIEAYHNYA-SLQGN-HHALYKEDLKKLVNTECPQYVK---------NKNAKTLFESLDINEDKAVNFEEFLVLMIKVGVEAHRDSH |
| S100A8\|Mus pahari\|XP_021052771.1 | S100A8 | F | F | aromatic | aromatic | ---------------------------------------------------------------------------------------------------------MPSEMEKALSSLIDVYHNYS-GIQGN-HHALYKDDFKKMVITECPQFGQ---------NKNNKNLFRELDINXDNAINFEEFLVMVIKVGVAYHKDSH |
| S100A8\|Mus musculus\|CAA61204.1 | S100A8 | F | F | aromatic | aromatic | ---------------------------------------------------------------------------------------------------------MPSELEKALSNLIDVYHNYS-NIQGN-HHALYKNDFKKMVTTECPQFVQ---------NINIENLFRDLDINSDNAINFEEFLAMVIKVGVASHKDSH |
| S100A8\|Grammomys surdaster\|XP_028609164.1 | S100A8 | F | F | aromatic | aromatic | ---------------------------------------------------------------------------------------------------------MLTELEKALNSMIDVYHNYS-TIKGN-HHAVYRDDFKKMATTECPQFVQ---------NKNTDNLFKELDVNSDNAVNFEEFLVLMIKMGVASHKDSH |
| S100A8\|Arvicanthis niloticus\|XP_034356824.1 | S100A8 | F | F | aromatic | aromatic | ---------------------------------------------------------------------------------------------------------MQSELEKALSNIIDVYHNYS-TIKGN-HHAVYRDDFKKMVTTECPQYVK---------NKNIENLFKEFDVNSDNAVNFEEFLVMTIRVGVAAHKDSH |
| S100A8\|Mastomys coucha\|XP_031231931.1 | S100A8 | F | F | aromatic | aromatic | ---------------------------------------------------------------------------------------------------------MPSELEKSLSNLIDVYHNYS-RLQGN-HHALYRDDFKKMVTTECPQFVK---------NKNTENLFRELDVNSDNAVNFEEFIVLLIRVGVAAHKDSH |
| S100A8\|Arvicola amphibius\|XP_038167988.1 | S100A8 | F | F | aromatic | aromatic | ---------------------------------------------------------------------------------------------------------MLSELEKSLENIVEVFHKYS-LVKGN-NHALYKDDFQKLVTTECPHYME---------KKNVETVFKELDINQDKVVNFEEFLVFLVKMGVAAHADSH |
| S100A8\|Microtus ochrogaster\|XP_005367318.1 | S100A8 | F | F | aromatic | aromatic | ---------------------------------------------------------------------------------------------------------MLSEMEKSLENMVEVYHRYS-LLKGN-NHALYKDDFKKLVTTECPHYMK---------KKNVETMFKELDINQDKAINFEEYLVLVVKMGVAAHADSH |
| S100A8\|Microtus oregoni\|XP_041512523.1 | S100A8 | F | F | aromatic | aromatic | ---------------------------------------------------------------------------------------------------------MLSELEKSLENMVEVYHRYS-LLKGN-NHALYKDDFKKLVTIECPQYIQ---------KKDIETVFKELDINQDKAINFEEYLVLLVKVGVAAHADSH |
| S100A8\|Cricetulus griseus\|XP_003511341.1 | S100A8 | L | F | hphobic | aromatic | ---------------------------------------------------------------------------------------------------------MPSELETALENIVTVYHRYS-EMKGN-HHALYRDDLQKLVTTECPQYMQ---------KKNAEALFKELDINQDNAVNFEEFLVLMIKIGVEAHKESH |
| S100A8\|Mesocricetus auratus\|XP_005080367.1 | S100A8 | L | F | hphobic | aromatic | ---------------------------------------------------------------------------------------------------------MLSELEKALENIVNVYHQYS-GTKGN-HHALYRDDLKKLLTTECPEFIQ---------KKNAETLFKELDVNQDNAVNFEEFLVLMIKVGLAAHEDSH |
| S100A8\|Peromyscus leucopus\|XP_028746685.1 | S100A8 | L | F | hphobic | aromatic | ---------------------------------------------------------------------------------------------------------MLSDLEKALESMVDVYHKYS-QVKGN-NHALYRDDLQKMLTTECPQYMQ---------RKKAEAVFRELDINQDNAINFEEFLVLVIKMGLAAHEDSH |
| S100A8\|Onychomys torridus\|XP_036047434.1 | S100A8 | L | F | hphobic | aromatic | ---------------------------------------------------------------------------------------------------------MLSDLEKALESMVEVYHKYS-EVKGN-HHALYRDDLQKLLTTECPQYVQ---------KKNAETIFKELDINKDNAVNFEEFLVLMIKVGVAAHEDSH |
| S100A8\|Jaculus jaculus\|XP_004667237.1 | S100A8 | M | F | hphobic | aromatic | ---------------------------------------------------------------------------------------------------------MLTELEKSLNAITEVFHKYS-LVKGN-YHALYKDDMKKLLSAECPQYMK---------KKDVDTWFRELDFNEDNAINFEEFLLFVTKMGVIAHRESH |
| S100A8\|Nannospalax galili\|XP_008853782.1 | S100A8 | L | F | hphobic | aromatic | ---------------------------------------------------------------------------------------------------------MLTDLEKSLNAIVDVFHKYS-LVKGN-HHALYRDDLKKLISNECPQYIQ---------KKDAETWFKELDFNEDGAVNFEEFLVLVIKVGVAAHRESH |
| S100A8\|Sorex araneus\|XP_004620092.1 | S100A8 | F | F | aromatic | aromatic | ---------------------------------------------------------------------------------------------------------MLTDLENSLNTLIDVFHKYS-LLKGN-NHAVDKNDFKKLIDTECPKYTK---------GKDSNTWFKELDINSDGAVNFEEYLILLVKIGVIAHKQSH |
| S100A8\|Artibeus jamaicensis\|XP_037015426.1 | S100A8 | F | F | aromatic | aromatic | ---------------------------------------------------------------------------------------------------------MLTELESALTNFVTVYHKYS-LLQGN-AHALYRNDFKKMLEAECPQYMK---------KKSADSWFRELDINSDKAINFEEFLILLVKIAVVAHKESH |
| S100A8\|Desmodus rotundus\|XP_024426406.1 | S100A8 | F | F | aromatic | aromatic | ---------------------------------------------------------------------------------------------------------MLTELESALSNFVTVYHKYS-LQQGN-AHALYKNDFKKMIEAECPQYLK---------KKNADAWFRELDINSDKAINFEEFLILVIKLGVKAHKESH |
| S100A8\|Dipodomys ordii\|XP_012870576.1 | S100A8 | L | F | hphobic | aromatic | ---------------------------------------------------------------------------------------------------------MPSDLECALDGVIDVYHRYS-LTQGN-YHSLYKDDLKKLLAAEVPNLLK---------NKDADTWFKVLDVNLDGAINFQEFLIFVIKMGVLAHEESH |
| S100A8\|Fukomys damarensis\|XP_010642045.1 | S100A8 | L | F | hphobic | aromatic | ---------------------------------------------------------------------------------------------------------MATELEKALNQVIEVYHKYS-LVHGN-PHALSKSDLKKLITTECPQYTK---------KKTATTWFKQLDINEDGAVNFEEFLHLVVKMGLRVHAESH |
| S100A8\|Heterocephalus glaber\|XP_004872792.1 | S100A8 | L | F | hphobic | aromatic | ---------------------------------------------------------------------------------------------------------METEMEKALNQVIEVYHKYS-LAQGN-PHALYKSDLKKLLTTECPQYTK---------KKTATAWFKQFDINEDGAVNFQEFLPLVMKVSLRAHEESH |
| S100A8\|Cavia porcellus\|XP_003478966.1 | S100A8 | L | F | hphobic | aromatic | ---------------------------------------------------------------------------------------------------------MLTELEKALDTIIDIYHKYS-LIKGN-PHALYKDDLKKLLTNECPQYTK---------KRNADAWFRQLDINEDGAVNFQEFLVFVVKVALKAHEESH |
| S100A8\|Chinchilla lanigera\|XP_005414659.1 | S100A8 | L | F | hphobic | aromatic | ---------------------------------------------------------------------------------------------------------MQTELEKALNNIITVYHKYS-LEKGN-HHAVYKDDLKKLLTTECPQYTK---------KKTADEWFKQLDVNEDGAINFQEFLVLVVKVALRAHEESH |
| S100A8\|Octodon degus\|XP_004632521.1 | S100A8 | L | F | hphobic | aromatic | ---------------------------------------------------------------------------------------------------------MPTELEKALDNVVSVYHKYS-EVKGN-HHAIYKDDLKKLLTKECPQYTK---------KKSAREWFRLLDINEDGAVNFQEFLVLVVKVALRAHEESH |
| S100A8\|Dasypus novemcinctus\|XP_004454077.1 | S100A8 | L | F | hphobic | aromatic | ---------------------------------------------------------------------------------------------------------MLTELETAITGLIKVYHDYS-LVKGN-YHSLYKDDLKKLLETESPKLVK---------NKSAETWFKELDINSDDAINFQEFLIVVIKMGVQMHKDSH |
| S100A8\|Tupaia chinensis\|ELW47339.1 | S100A8 | L | F | hphobic | aromatic | ---------------------------------------------------------------------------------------------------------MPTELECALNTVIDVFHKYS-LEKGS-HHAVYRDDLKKLLDVECPQYVK---------NKDADTWFKELDINKDGGVNFEEFLILVVKMGVAGHEESH |
| S100A8\|Ceratotherium simum simum\|XP_004436372.1 | S100A8 | L | F | hphobic | aromatic | ---------------------------------------------------------------------------------------------------------MLTDLEEAMDSLIDVYHKYS-LKKGN-YHSVYKDDLKRLLETECPHYTK---------EKDADTWFNELDVNRDGAINFEEFLILVIKVGLEAHRKSH |
| S100A8\|Equus caballus\|XP_001493557.2 | S100A8 | L | F | hphobic | aromatic | ---------------------------------------------------------------------------------------------------------MLTDLEKAMDSLIDVYHKYS-LVKGN-YHAVYKEDLKKLLETECPQYTK---------EKNADAWFKELDFTADGAINFEEFLILVIKVALAAHEASH |
| S100A8\|Condylura cristata\|XP_004693774.1 | S100A8 | L | F | hphobic | aromatic | ----------------------------------------------------------------------------------------------------------MTELEKALNSLINVYHKYS-LEKGN-YHALYSSDLKNLLEKECPDYVK---------KKSADVWFKELDINSDQAINFQEYLIFVVKIGVLGHKESH |
| S100A8\|Talpa occidentalis\|XP_037356156.1 | S100A8 | L | F | hphobic | aromatic | ----------------------------------------------------------------------------------------------------------MTDLERAMNSLINVYHKYS-LEKGN-YHALYDSDLKKLLEVECPDYVK---------RKSADAWFKELDINSDKAINFEEYLIFVIKIGVLGHKESH |
| S100A8\|Marmota marmota marmota\|XP_015358168.1 | S100A8 | L | F | hphobic | aromatic | ---------------------------------------------------------------------------------------------------------MPTDLEKALDSLIDVFHKYS-LEKGN-YHAIYRDDLRKLLTKECPHYTK---------NKDADTWFKELDVNADNAVNFEEYLVLVIKVGLAAHKESH |
| S100A8\|Ictidomys tridecemlineatus\|XP_005331308.2 | S100A8 | L | F | hphobic | aromatic | ---------------------------------------------------------------------------------------------------------MPTDLEKALDSLVCVFHKYS-LKKGN-YHALYRDDLKKMLTTECPHYTK---------NKDADTWFKELDVNADNAVNFEEYLVLVIKVGVEAHKLSH |
| S100A8\|Ochotona princeps\|XP_004588973.2 | S100A8 | L | F | hphobic | aromatic | ---------------------------------------------------------------------------------------------------------MPTDLENSLNSIISVYHKYS-LEKGN-YHALYRDDLRKLLATECPQYVK---------KKDADTWFRELDVNSDNAINFQEFLILVVKIGVIAHEDSH |
| S100A8\|Oryctolagus cuniculus\|XP_002715389.1 | S100A8 | L | F | hphobic | aromatic | ---------------------------------------------------------------------------------------------------------MPTDLENSLNSIISVYHKYS-LEKGN-YHALYGDDLKKLLATECPQYTK---------KKDADTWFKELDINSDGAINFQEFLILIVKIGVSAHEDSH |
| S100A8\|Castor canadensis\|XP_020041647.1 | S100A8 | L | F | hphobic | aromatic | ---------------------------------------------------------------------------------------------------------MPTELECSLNSLIDVYHKYS-LIKGN-YHAIYKDDLKKLLVNECPHYIK---------IKDADSWFKELDINLDGAINFQEYLIFVIKVGLAAHEDSH |
| S100A8\|Galeopterus variegatus\|XP_008565857.1 | S100A8 | M | F | hphobic | aromatic | ---------------------------------------------------------------------------------------------------------MLTDLENALDSFIDVYHKYS-LLQGN-YHSIYMDDMKKLLEKECPQFIK---------KKSAETWFKELDINSDGAINFEEFLVMVVKVGVEAHKEIH |
| S100A8\|Carlito syrichta\|XP_008062093.1 | S100A8 | L | F | hphobic | aromatic | ---------------------------------------------------------------------------------------------------------MPTDLEKSLNSVIDIYHKYS-LIKGN-YHAIYKDDLKKLLETECPQYTK---------RKDAESWFKDLDVNTDGAINFQEFLIFVVKIGVEAHKDSH |
| S100A8\|Otolemur garnettii\|XP_003802330.2 | S100A8 | L | F | hphobic | aromatic | ---------------------------------------------------------------------------------------------------------MPTDLECALNSVIDVYHKYS-LRKGN-YHAIYRDDLKSLLETECPQYMK---------KKDANTLFKELDINSDGAINFQEFLVLMIKVGVAAHEDSH |
| S100A8\|Propithecus coquereli\|XP_012514078.1 | S100A8 | L | F | hphobic | aromatic | ---------------------------------------------------------------------------------------------------------MLTDLENALDSFIDVYHKYS-LRQGN-YHAIYIDDLKQLLETECPQYMK---------KKDANTWFKELDINSDGAINFQEFLMFVIKLGVAAHEDIH |
| S100A8\|Microcebus murinus\|XP_012646628.1 | S100A8 | L | F | hphobic | aromatic | ---------------------------------------------------------------------------------------------------------MLTDLENALDSFIDVYHKYS-LRKGN-YHAIYKDDLKDLLETECPQYMK---------KKDANTWFKELDVNSDNAINFQEFLIFVTKMGVVAHEESH |
| S100A8\|Cebus imitator\|XP_017368781.1 | S100A8 | L | F | hphobic | aromatic | ---------------------------------------------------------------------------------------------------------MLTDLENALNSLIEVYHKYS-LIKGN-YHSIYRDDLKKLLETECPAYIN---------KKSAEAWFNELDINSDNAINFQEFLILVVKVGVAAHKESH |
| S100A8\|Aotus nancymaae\|XP_012293732.1 | S100A8 | L | F | hphobic | aromatic | ---------------------------------------------------------------------------------------------------------MLTDLEKALNSLIDVYHKYS-LIKGN-YHAIYRDDLKKLLETECPQYIK---------KKSAEAWFKELDVNSDNAINFQEFLILVIKVGVQAHKESH |
| S100A8\|Homo sapiens\|CAA68390.1 | S100A8 | L | F | hphobic | aromatic | ---------------------------------------------------------------------------------------------------------MLTELEKALNSIIDVYHKYS-LIKGN-FHAVYRDDLKKLLETECPQYIR---------KKGADVWFKELDINTDGAVNFQEFLILVIKMGWQPTKKAM |
| S100A8\|Nomascus leucogenys\|XP_030680359.1 | S100A8 | L | F | hphobic | aromatic | ---------------------------------------------------------------------------------------------------------MLTELEKALNSIIDVYHKYS-LIKGN-YHAVYRDDLKKLLETECPQYIR---------KKGADAWFKELDINTDGAVNFQEFLILVVKMGVAAHKKSH |
| S100A8\|Trachypithecus francoisi\|XP_033052956.1 | S100A8 | L | F | hphobic | aromatic | ---------------------------------------------------------------------------------------------------------MLTNLEKALDSIIQVYHKYS-LIKGN-YHAVYKDDLKKLLETECPQYIR---------KKGADAWFKELDINTDGAINFQEFLILVIKMGVAAHKESH |
| S100A8\|Myotis lucifugus\|XP_006095368.1 | S100A8 | F | F | aromatic | aromatic | ---------------------------------------------------------------------------------------------------------MLTEMENTLNNFIEIYHKYS-KLQGS-DHALYGNDFKKLLEDECPHFLQ---------NKNAATWFKELDINRDNAINFQEYLTLVIKVGLHAHDEIH |
| S100A8\|Eptesicus fuscus\|XP_008145468.1 | S100A8 | L | F | hphobic | aromatic | ---------------------------------------------------------------------------------------------------------MLTELESTLNNFIEIYHRYS-KLQGN-DHALYRDDLKKLLEAECPQYMK---------KKSADTWFRELDINEDNAINFQEYLTLVIKVGLAAHEEIH |
| S100A8\|Miniopterus natalensis\|XP_016075617.1 | S100A8 | L | F | hphobic | aromatic | ---------------------------------------------------------------------------------------------------------MLTEMENTLNNFIEIYHKYS-LAQGS-PHALYRDDLKKLLEAECPQYMK---------KKDANTWFRELDINKDNAINFEEFLIMVIKMGVEAHEDIH |
| S100A8\|Loxodonta africana\|XP_003415124.1 | S100A8 | F | F | aromatic | aromatic | ---------------------------------------------------------------------------------------------------------MLTDLERAIDSLIEVFHKYS-LLKGN-AHALYRDDFQKLLETEGRRYLR---------IKDANAWFKELDVNSDSAINFEEFLILVVKMGVIAHEDIH |
| S100A8\|Molossus molossus\|XP_036128612.1 | S100A8 | L | F | hphobic | aromatic | ---------------------------------------------------------------------------------------------------------MLTELESALNNFIDIYHKYS-QDRGN-VHALYRDELKRLLEAECPQYIK---------KKDADTWFKELDINKDNAINFEEFLVLVIKVGVASHEDFH |
| S100A8\|Pteropus alecto\|XP_006923667.1 | S100A8 | L | F | hphobic | aromatic | ---------------------------------------------------------------------------------------------------------MLTELESAMNNFIDIFHRYS-LEKGN-YHALYRDDLKKLLEAECPQYMK---------KKDADTWFKELDVNKDSAVNFEEFLALVVKVGVAAHEDIH |
| S100A8\|Hipposideros armiger\|XP_019482850.1 | S100A8 | L | F | hphobic | aromatic | ---------------------------------------------------------------------------------------------------------MLTELESAINNFIEIFHKYS-LEKGN-YHALYRDDLKKLLLAECPQYLK---------KKDVDTWFRELDVNQDNAVNFEEFLVLVVKLGIASHEDSH |
| S100A8\|Rhinolophus ferrumequinum\|XP_032949090.1 | S100A8 | L | F | hphobic | aromatic | ---------------------------------------------------------------------------------------------------------MLTELESAINNFIEIYHKYS-LETGN-YHALYRDDLKKLLHTECPQYMK---------KKDADTWFKELDINQDNAINFEEFLTLVVKVGITAHEDSH |
| S100A8\|Echinops telfairi\|XP_004717760.1 | S100A8 | M | F | hphobic | aromatic | ---------------------------------------------------------------------------------------------------------MLTDLEKALNQLIGVYHKYS-LLQGN-YHALYRNDMKKLLEVEGPRYLK---------IKSADAWFKELDINCDNAINFQEFLILVVKMGVIAHEDSH |
| S100A8\|Elephantulus edwardii\|XP_006895952.1 | S100A8 | M | F | hphobic | aromatic | ---------------------------------------------------------------------------------------------------------MLTELEQAIDSLIDVYHKYS-LKKGN-PHAIYRDDMKRLLETEANNYLK---------KKSADAWFNELDINSDRAINFQEFLIFVVKMGVIAHEDIH |
| S100A8\|Balaenoptera physalus\|KAB0396308.1 | S100A8 | L | F | hphobic | aromatic | ---------------------------------------------------------------------------------------------------------MLTDLESAMNCIVDVYHKYS-LVKGN-YHSIYRDDLKKLLQTECPKFLE---------KKDADTWFKELDVNEDDALNFREFLSLVIKVGLAAHEDIH |
| S100A8\|Physeter catodon\|XP_007124144.1 | S100A8 | L | L | hphobic | hphobic | ---------------------------------------------------------------------------------------------------------MLTDLECAMNSLIEVYHKYS-LLKGN-YHAIYQDDLKKLLETECPKFLE---------KKDVDTWLKELDINEDGALNFQEFLSLVIKVGLAAHEDIH |
| S100A8\|Ovis aries\|KAG5215317.1 | S100A8 | L | F | hphobic | aromatic | ---------------------------------------------------------------------------------------------------------MLTDLESAINSLIDVYHNYS-LLKGN-YHAVYRDDLKRLLETECPKFLK---------KKDADTWFKELDINQDGGINFEEFLVLVIKLTAQVSTSPT |
| S100A8\|Orycteropus afer afer\|XP_007946780.1 | S100A8 | L | F | hphobic | aromatic | ---------------------------------------------------------------------------------------------------------MLTDLENAMDSLIDVYHKYS-LVQGN-YHALYRDDLKKLLENEAPTYLK---------KKDADAWFKELDINTDGAINFQEFLIMVIKMGVIAHEDIH |
| S100A8\|Manis pentadactyla\|XP_036778081.1 | S100A8 | L | F | hphobic | aromatic | ---------------------------------------------------------------------------------------------------------MLTDLECALNSLVDVYHKYS-LLKGN-YHAIYKDDLKKLLETECPHFLK---------QKDADTWFKELDVNQDGGINFEEFIALVIKVGLAAHEAIH |
| S100A8\|Trichechus manatus latirostris\|XP_004388664.1 | S100A8 | L | F | hphobic | aromatic | ---------------------------------------------------------------------------------------------------------MLTDLENAMDSLITVYHKYS-LVKGN-AHALYKDDLKNLLQIEGPCYLK---------IKDADTWFKELDINSDNAINFQEFLILVIKMGVIAHHDIH |
| S100A8\|Chrysochloris asiatica\|XP_006861513.1 | S100A8 | L | F | hphobic | aromatic | ---------------------------------------------------------------------------------------------------------MLTDLESAMDSLIEVYHNYS-LVKGN-YHALYKDDLKKLLETEGPNYLK---------KKDADTWFKELDINSDKAINFEEFLILVIKLGVIAHEDSH |
| S100A8\|Halichoerus grypus\|XP_035922750.1 | S100A8 | L | F | hphobic | aromatic | ---------------------------------------------------------------------------------------------------------MLTELESAMNCLIEVYHNYS-LEKGN-YHALYRDDLKKLLETECPRYLK---------KKDADTWFKELDINSDGAINFEEFLILVIKVGVEAHKEIH |
| S100A8\|Vulpes vulpes\|XP_025852663.1 | S100A8 | L | F | hphobic | aromatic | ---------------------------------------------------------------------------------------------------------MLTELESAINSLIEVYHKYS-LVKGN-YHALYRDDLKKLLETECPQFMK---------KKDADTWFQELDINSDGAINFEEFLILVIKVGVASHKDIH |
| S100A8\|Crocuta crocuta\|KAF0880567.1 | S100A8 | L | F | hphobic | aromatic | ---------------------------------------------------------------------------------------------------------MLTELESAINSIIDVFHNYS-LEKGN-YHALYRDDLKKLLETECPRYMK---------KKDADTWFKELDINSDGAVNFEEFLILVIKVGVATHHEIH |

| | | | | | | |
|---|---|---|---|---|---|---|
| S100A8\|Suricata suricatta\|XP_029789123.1 | S100A8 | L | F | hphobic | aromatic | -----------------------------------------------------------------------------------------------------------------------------------------------------------------MLTDLETAINSIIDVYHKYS-LAKGN-YHAVYRDDLKKMLETECPRYMK---------KKDADTWFKELDVNSDDAINFEEFLILVIKVGVAAHHDIH |
| S100A8\|Panthera pardus\|XP_019286764.1 | S100A8 | L | F | hphobic | aromatic | -----------------------------------------------------------------------------------------------------------------------------------------------------------------MLTELETAINSLIDVYHKYS-LEKGN-YHALYRDDLKKLLETECPRYMK---------KKDADTWFKELDVNSDGAINFEEFLILVIKVGVATHEDIH |
| S100A8\|Sus scrofa\|NP_001153743.1 | S100A8 | L | F | hphobic | aromatic | -----------------------------------------------------------------------------------------------------------------------------------------------------------------MLTDLESAINSLIEVFHKYS-LEKGN-YHAIYADDLKRLLETECPKYMK---------KKDAETWFKELDINKDGAVNFEEFLILVIKVGVEAHEDIH |
| S100A8\|Camelus bactrianus\|XP_010952357.1 | S100A8 | L | F | hphobic | aromatic | -----------------------------------------------------------------------------------------------------------------------------------------------------------------MLTDLEGAINNIIQVYHNYS-LLKGN-HHAIYRDDLKKLLETECPQFLK---------KKNTDTWFKELDINEDGAVNFEEFLILVIKVGLAAHEDIH |
| S100A8\|Equus przewalskii\|XP_008541236.1 | S100A8 | L | F | hphobic | aromatic | -----------------------------------------------------------------------------------------------------------------------------------------------------------------MLTDLENAINSLIDVYHNYS-LLKGN-YHAIYRDDLKKLLETECPQYTK---------KKNADAWFKELDINEDGAVNFQEFLILVVKVGLLAHEDIH |
| S100A8\|Ceratotherium simum simum\|XP_014647225.1 | S100A8 | L | F | hphobic | aromatic | -----------------------------------------------------------------------------------------------------------------------------------------------------------------MLTDLENAINSLIDVYHKYS-LIKGN-YHAVYRDDLKKLLETECPQYLK---------KKDADTWFKELDINEDGAVNFEEFLILVIKVGLLAHEDIH |
| S100A8\|Bos taurus\|NP_001107197.1 | S100A8 | L | F | hphobic | aromatic | -----------------------------------------------------------------------------------------------------------------------------------------------------------------MLTDLECAINSLIDVYHKYS-LKKGN-YHAVYRDDLKQLLETECPKFMK---------KKDADTWFKELDINQDGGINFEEFLVLVIKVGLEAHEEIH |
| S100A8\|Cervus hanglu yarkandensis\|KAF4021311.1 | S100A8 | L | F | hphobic | aromatic | -----------------------------------------------------------------------------------------------------------------------------------------------------------------MLTDLESAINSLIEVYHRYS-LLKGN-YHAVYRDDLKKLLETECPKYMK---------KKDADTWFKELDINQDGGVNFEEFMVLVIKVGLATHEDIH |
| MRP-126\|Gekko japonicus\|XP_015265128.1 | MRP-126 | L | M | hphobic | hphobic | ----------------------------------------------------------------------------------------------------------------------------------------------------------------MAQTEMEKHCEGIINGFHKFS-IRVDH-HDMLSIGELKQLLEQQLPTYLK----K-NKD-AA-KVLMDELDKDKNKQLDFAEFMGVITRVLIFSHDNIH |
| MRP-126\|Pantherophis guttatus\|XP_034291533.1 | MRP-126 | F | F | aromatic | aromatic | ----------------------------------------------------------------------------------------------------------------------------------------------------------------MSQTQMEKCCECIIDIFHKYS-IQKEH-YDMLSKAEFAKMLKEQFPNFIQ-----GSKKPEDIDRLFKELDQNKDGQLSFEETMAFIGKFLIACHHKFH |
| MRP-126\|Pseudonaja textilis\|XP_026575966.1 | MRP-126 | L | F | hphobic | aromatic | ----------------------------------------------------------------------------------------------------------------------------------------------------------------MSQTQMEKCCECIIDVFHKYS-IQKEH-HDRLSKAELSKLLKEQLPNFIQ-----GSKDPAAIDRLFKELDQNKDQEVSFEEFMAFVSKVLIHCHNKFH |
| MRP-126\|Thamnophis elegans\|XP_032089922.1 | MRP-126 | F | F | aromatic | aromatic | ----------------------------------------------------------------------------------------------------------------------------------------------------------------MSQTQLEKCCQCIIDVFHKYS-KLTDH-HDRLSKAEFAKMLREQIPNFLK-----DNKNPDGVNRLFNELDQNKDRELSFEEFMTFLGRFLVQCHNEFH |
| MRP-126\|Protobothrops mucrosquamatus\|XP_015668886.1 | MRP-126 | F | F | aromatic | aromatic | ----------------------------------------------------------------------------------------------------------------------------------------------------------------MSKTQLEKCCDCIMDVFHKYS-TITEH-PDMLSKGEFANLLREQLPNYIQ-----STKNTEAMDRLFNELDKNKDQMLSFEEFMTLVGRFMIQCHDDFH |
| MRP-126\|Crotalus tigris\|XP_039219247.1 | MRP-126 | F | F | aromatic | aromatic | ----------------------------------------------------------------------------------------------------------------------------------------------------------------MSKTQLEKCCDCIMDVFHKYS-MPLAH-PDKLSKGEFAKMLKEQLPNFIQ-----STKNTEAMNRLFNELDQNKDGELSFEEWLTLVGRFLAQCHENFH |
| S100A9\|Trichosurus vulpecula\|XP_036613608.1 | S100A9 | M | M | hphobic | hphobic | ---------------------------------------------------------------------------------------------------------------------------------------------------------------MTQQCTLKNALQTFVDTFHKYS-VKYED-RDALNCKEMKQLLNKEMPNFIK-----NSKELKDIPNLMIELDTNLDKELSFNEFSVMLARVILESHEKMH |
| S100A9\|Monodelphis domestica\|XP_001372115.3 | S100A9 | M | M | hphobic | hphobic | ----------------------------------------------------------------------------------------------------------------------------------------------------------------MENCTMEKALDIIVNTFHHYS-TRVGN-PDTLVKGEMKQLITKELPNFIK-----NAKDLQDVKHLMQELDTNQNGQVDFKEFSMMMARLTMATHEKMH |
| S100A9\|Sarcophilus harrisii\|XP_003768080.1 | S100A9 | M | F | hphobic | aromatic | ----------------------------------------------------------------------------------------------------------------------------------------------------------------MESCTLKDCLKIIIDTFHQYS-GQGGD-PDALNKKEMKQLLTKEMPHFIK-----NSKDLQCVPHLFQELDTNQDHDINFKEYAVMIARLAMASHEKMH |
| S100A9\|Vombatus ursinus\|XP_027726505.1 | S100A9 | M | F | hphobic | aromatic | ----------------------------------------------------------------------------------------------------------------------------------------------------------------MEACSLKNSLKIFVDTFHKYS-NISGN-KDALNRREMKTLLNKEMPHFIK-----NSKDLKNVSHLFEELDTNQDADVDFKEFAVMIARVVMETHEKMH |
| S100A9\|Phascolarctos cinereus\|XP_020847685.1 | S100A9 | M | F | hphobic | aromatic | ----------------------------------------------------------------------------------------------------------------------------------------------------------------METCSLKSSLKIFVDVFHKYS-KISGN-LDALNRREMKQLLNKEMPHFIK-----NSKDLKDVQSLFEELDTNLDSDIDFKEFAVMIARVVMETHEKMH |
| S100A9\|Fukomys damarensis\|XP_010642044.1 | S100A9 | F | M | aromatic | hphobic | --------------------------------------------------------------------------------------------------------------------------------------------------------------MAGEMSVFERSLNSIVEAFHQYS-RREDN-NDTLNKKEFKLLMQKELPSYLK----KNEDGKDVVKDIMEDLDTNLDNELSFEEAIMLFSRVINASHETMH |
| S100A9\|Ornithorhynchus anatinus\|XP_028911801.1 | S100A9 | L | L | hphobic | hphobic | ---------------------------------------------------------------------------------------------------------------------------------------------------------------------MEKSLENIINVFHQYS-VRVGN-PDTLTKRELRQLINKELPNFLK-----DQQCPADVAKILEDLDSNQDSQLSFEEFVVLITRLTVASHNKMH |
| S100A9\|Tachyglossus aculeatus\|XP_038624912.1 | S100A9 | L | L | hphobic | hphobic | ---------------------------------------------------------------------------------------------------------------------------------------------------------------------MEKALGDVIDVFHQYS-VRVGN-PDTLTKRELRQLINKELPNFLK-----DQQCPAEVAKILEDLDSNQDFELSFEEFVVLITRLTVASHNKMH |
| S100A9\|Erinaceus europaeus\|XP_007525487.2 | S100A9 | F | M | aromatic | hphobic | --------------------------------------------------------------------------------------------------------------------------------------------------------------LAGTMSKMEQSLDTIIDTFHKYS-KELNH-PDTLTQGEFKKLVQRDLTNFLK----EQKRDEVAIGKIMEDLDSSADKQISFPEFIMLIARLAVASHEKMH |
| S100A9\|Heterocephalus glaber\|XP_004871186.1 | S100A9 | F | M | aromatic | hphobic | -----------------------------------------------------------------------------------------------------------------------------------------------------YLLGSECKKMTSEMSLLERSLDTTIKTFHRYS-TQEGN-LDTLSKKEFKQLVQRELPNFLK----NESRNEAAINDIMEDLDTNLDQQLSFEELTMLLARLINASHEKMH |
| S100A9\|Cavia porcellus\|XP_003478965.1 | S100A9 | F | M | aromatic | hphobic | --------------------------------------------------------------------------------------------------------------------------------------------------------------MAANKSQLERSLETIINTFHKYS-RLSGH-PDTLNKNEFKKLTETELANFLK----CENRKEDEINDIMQDLDTNQDQQLSFEEVIMLFARLINASHEKMH |
| S100A9\|Chinchilla lanigera\|XP_005414660.2 | S100A9 | F | M | aromatic | hphobic | -----------------------------------------------------------------------------------------------------------------------------------------------------CLLGSECEKTTAEMSLLERSLETIIDTFHQYS-TKAGH-PDTLNKREFRALAKTELPNFLK----C-----GDINDIMEDLDTNQDQQLSFEEVSMLFARLTNASHEKMH |
| S100A9\|Octodon degus\|XP_023572141.1 | S100A9 | F | L | aromatic | hphobic | -----------------------------------------------------------------------------------------------------------------------------------------------------CLLGSEYKKMAAEMSQLESSLNTIIDTFHHYS-TKEGH-KDTLNKKEFSGLVKTELPNFLK----CESRDEAEINDILEDLDTNQDQQLSFEEVIMLFAKLINATHEKMH |
| S100A9\|Carlito syrichta\|XP_008062090.1 | S100A9 | F | M | aromatic | hphobic | --------------------------------------------------------------------------------------------------------------------------------------------------------------MAGEKSLMEKNLETIIDTFHHYS-VRVGH-PDTLNKKEFKDLIKKELPNFLK----KVKKDDAEISHMMEDLDTNQDKQLSFEEFIIMIVRLIYASHEKMH |
| S100A9\|Nannospalax galili\|XP_008853793.1 | S100A9 | F | M | aromatic | hphobic | ------------------------------------------------------------------------------------------------------------------------------------------------------------------MSQLERSIETIIDTFHKHS-TKEGH-PDTLSQKEFKNVVNNDLPNFLK----KEKKEDKVINDIMEDLDTNQDKQLSFEEFTMLIAKLVHATHEKMH |
| S100A9\|Mesocricetus auratus\|XP_005080299.2 | S100A9 | F | M | aromatic | hphobic | -------------------------------------------------------------------------------------------------------------------------------------------------------------MATKTPSLLERSIETIINIFHKYS-VEGGH-PDTLSQGEFKKLVKTDLANFLK----KEKRKDELINDIMEDLDTDLNKQLSFEEFVILMAKMVHATHEKMH |
| S100A9\|Cricetulus griseus\|XP_035306499.1 | S100A9 | F | M | aromatic | hphobic | -------------------------------------------------------------------------------------------------------------------------------------------------------------MATKTPSLLEQSITTIIDTFHKYS-TQEGH-PDTLSQREFIKMVKTDLANFLK----KEKRKVNLINDIMEDLDTNLDKQLSFEEFSILMAKMVYATHEKMH |
| S100A9\|Peromyscus leucopus\|XP_028746688.1 | S100A9 | F | M | aromatic | hphobic | -------------------------------------------------------------------------------------------------------------------------------------------------------------MATKTPSVMERNINTVIDVFHQYS-RKGGN-PDTVRQQEFKEMVKTDLATFLK----KEKKDETLINDIMEDLDTNQDKQLSFEEYTMLIAKLIFATHEKLH |
| S100A9\|Peromyscus maniculatus bairdii\|XP_006976289.1 | S100A9 | F | M | aromatic | hphobic | -------------------------------------------------------------------------------------------------------------------------------------------------------------MATKTPSVMERNINTVIDVFHQHS-RKGGN-PDTLSQQEFKKLVETDLATFLK----EEKKDERVINDIMEDLDTNQDKQLSFEEFTMLIAKLIFATHEKLH |
| S100A9\|Microtus oregoni\|XP_041511029.1 | S100A9 | F | M | aromatic | hphobic | -------------------------------------------------------------------------------------------------------------------------------------------------------------MANQNPSLMERSIDTVIDVFHTNS-KMEGD-PDTLSQKEFKQMVKKDLANFMK----KEKRRDKIINDIMEDLDTNQDKQLSFEEFVMLVAKLVYATHEKMH |
| S100A9\|Onychomys torridus\|XP_036047206.1 | S100A9 | F | M | aromatic | hphobic | -------------------------------------------------------------------------------------------------------------------------------------------------------------MATKTPSMMERNINTIIEVFHQHA-RTEGN-TDTLSQKEFKNMVKSDLANFMK----KEKKKEKLINDIMEDLDTNDDKQLAFEEFVMLVAKLVFATHEKMH |
| S100A9\|Meriones unguiculatus\|XP_021520239.1 | S100A9 | F | M | aromatic | hphobic | -------------------------------------------------------------------------------------------------------------------------------------------------------------MAAKTPSLMERSIDTIIHTFHKHS-RKEGN-RDTLNKKEFKELVTKDLANFMK----KEKRCENLLTDVMEDLDTNQNEELSFEEFIMLIAKLVFATHESLH |
| S100A9\|Arvicanthis niloticus\|XP_034356807.1 | S100A9 | F | F | aromatic | aromatic | -------------------------------------------------------------------------------------------------------------------------------------------------------------MATKTPSQMERSIATIINTFHQHS-RKGGN-DDTLSKKEFKDMVTMDLANFLK----KEKRDDKLINDIFEDLDTNQDNQLSFEESMMLMAKLVFACHEKLH |
| S100A9\|Grammomys surdaster\|XP_028609137.1 | S100A9 | F | F | aromatic | aromatic | -------------------------------------------------------------------------------------------------------------------------------------------------------------MATKTSSLMERSITTVINTFHQHS-RKEGN-DDTLSKKEFKEMVKTDLATFLK----KEKRNERLINDIFEDLDTNQDNQLSFEECMMLMAKLVFACHEKLH |
| S100A9\|Mus musculus\|NP_001268781.1 | S100A9 | F | M | aromatic | hphobic | -------------------------------------------------------------------------------------------------------------------------------------------------------------MANKAPSQMERSITTIIDTFHQYS-RKEGH-PDTLSKKEFRQMVEAQLATFMK----KEKRNEALINDIMEDLDTNQDNQLSFEECMMLMAKLIFACHEKLH |
| S100A9\|Mastomys coucha\|XP_031231932.1 | S100A9 | F | M | aromatic | hphobic | --------------------------------------------------------------------------------------------------------------------------------------------------------------MATRTSQMERNIGTIIDTFHQYS-RREGH-HDTLSKKEFRDLVKMELATFMK----KENRSEAIINDIMEDLDTNQDNQLTFEESMMLMAKLVFACHEKLH |
| S100A9\|Sturnira hondurensis\|XP_036922938.1 | S100A9 | F | L | aromatic | hphobic | -----------------------------------------------------------------------------------------------------------------------------------------------------------------MPSLMECSIENVIKVFHQYS-RRQEH-LDRLNQSEFKELVQKELPTFLK----KEGRDEKAIEEILEDLDTNGDKQLSFEEFSMLVAKLTEAAHEEMH |
| S100A9\|Artibeus jamaicensis\|XP_036982243.1 | S100A9 | F | L | aromatic | hphobic | -----------------------------------------------------------------------------------------------------------------------------------------------------------------MPSLMECSLDHVIKVFHQYS-RQQEH-LDRLNQSEFKEMVKKELPTFLQ----KEGRDEKAIEDILEDLDTNGDKQLSFEEFAMLVAKLTEAAHEEMH |
| S100A9\|Phyllostomus discolor\|XP_028359064.1 | S100A9 | F | L | aromatic | hphobic | -----------------------------------------------------------------------------------------------------------------------------------------------------------------MPSLMECSINNVIKIFHQYS-RQEAH-TDRLNQNEFKELVQKELPNFLQ----KENRNEAAIKEILEDLDTSGDKQLSFEEFAMLVAKLTEAAHEEMH |
| S100A9\|Desmodus rotundus\|XP_024426173.1 | S100A9 | F | L | aromatic | hphobic | -----------------------------------------------------------------------------------------------------------------------------------------------------------------MPSLLECSIDTVIKIFHQYS-TEESH-HDRLSQNEFKELVQKELPNFLK----KEDRDEEAIKEILEDLDTNGDKQLSFEEFSMLVAKLTEAAHEEMH |
| S100A9\|Tupaia chinensis\|ELW47341.1 | S100A9 | F | L | aromatic | hphobic | -------------------------------------------------------------------------------------------------------------------------------------------------------------KIAGEMSQMENSINTIINVFHRYS-VRLDH-FDKLNKKEFGQLVRKELPNFLK----KEKKDETLINDILEDLDTNADGELSFEEYMMMMARLIVASHEEMH |
| S100A9\|Sorex araneus\|XP_004620094.1 | S100A9 | F | M | aromatic | hphobic | -----------------------------------------------------------------------------------------------------------------------------------------------------------------MASRMEASIENIINTFHQYS-TRLKH-PDTLNYAEFKDLVKKELPNFLK----QKNRDEAALKDIMEDLDTNADKQLSFEEFTMLVAKLVVASHEEMH |
| S100A9\|Condylura cristata\|XP_004693683.1 | S100A9 | F | M | aromatic | hphobic | ------------------------------------------------------------------------------------------------------------------------------------------------------------------MSELERSIDVIINTFHQYS-VRRDD-PDTLSQQEFKQLARKELPNFLQ------KKDNVTLQKIMEDLDTNADKQLSFEEFAILVGRLTYASHEEMH |
| S100A9\|Talpa occidentalis\|XP_037356696.1 | S100A9 | F | M | aromatic | hphobic | ----------------------------------------------------------------------------------------------------------------------------------------------------------GCKRIVAEMSELERSIDVIINTFHHYS-VRKGD-HDTLSQQEFKQLAKKELPNFLQ------KKDDAALLKIMEDLDTTVDNQLSFEEFAILVGRLTHASHQEMH |
| S100A9\|Dipodomys ordii\|XP_012870427.1 | S100A9 | F | L | aromatic | hphobic | --------------------------------------------------------------------------------------------------------------------------------------------------------------MADTRPQMEKHLETIVDVFHQYS-NRQGN-RDTLNKKEFKQLVKKELQNFLK----KENRDEKEINEILEDLDTNQDQTLDFNEFAILIARLINASHEEMH |
| S100A9\|Jaculus jaculus\|XP_012806289.1 | S100A9 | F | M | aromatic | hphobic | -----------------------------------------------------------------------------------------------------------------------------------------------------------------MATQLERSLDTIIHTFHEHS-VKEGN-RDTLSQKEFKDLVKKDLQNFLK----KEKRNEQIINDIMEDLDTNQDKMLSFEEFIILIAKLVHASHEEMH |
| S100A9\|Balaenoptera physalus\|KAB0389423.1 | S100A9 | F | M | aromatic | hphobic | ------------------------------------------------------------------------------------------------------------------------------------------------------------------MSQLESSIETIINIFHQYS-VRLQH-PDTLNHKEFKQLVEKELPNFLK------YKDDKAINKIMEDLDTDVDKQMDFEEFLMLVARLTEASHEEMH |
| S100A9\|Physeter catodon\|XP_007124462.1 | S100A9 | F | M | aromatic | hphobic | ------------------------------------------------------------------------------------------------------------------------------------------------------------------LSQLESSIETIINIFHQYS-IRLQH-PDTLNKKEFKQLVIKELPNFLK----KEKKDDQAISDIMEDLDTDVDGELNFQEFSVLVGKLTEASHEEMH |
| S100A9\|Sousa chinensis\|TEA35521.1 | S100A9 | F | M | aromatic | hphobic | ----------------------------------------------------------------------------------------------------------------------------------------------------------ECRKMADQLSQLESSIETIINIFHQYS-IRLQP-PDTLNKKEFKQLVKKELPNFLK----KETKDDKAINEIMEDLDTDVDKELNFHEFSVLVGKLTEASHEEMH |
| S100A9\|Dasypus novemcinctus\|XP_004454075.1 | S100A9 | F | M | aromatic | hphobic | --------------------------------------------------------------------------------------------------------------------------------------------------------------MSEKMTQMECYVETIINVFHQYS-VRLGN-PDTLNQKEFKQLVKKELPNFLK----KEKKNDEILNHIMEDLDTNLDKTLSFEEYAILIARVIHATHEEMH |
| S100A9\|Loxodonta africana\|XP_003414949.1 | S100A9 | F | M | aromatic | hphobic | --------------------------------------------------------------------------------------------------------------------------------------------------------------MADQLSQLERDIETIINVFHQYS-VRLEH-RDALNQKEFKQLVQKELANFLK----REKKNDAVINDIMEDLDTNEDKELSFEEYIVLVAKLTEASHEKMH |
| S100A9\|Ictidomys tridecemlineatus\|XP_005331306.1 | S100A9 | F | L | aromatic | hphobic | --------------------------------------------------------------------------------------------------------------------------------------------------------------MASELSQMESSIETVINVFHQYA-VRLGN-RDTLNQKEFIQMVQKELQNYLK----KEGKNEKLINHILEDLDTNVDKQLSFEEFVMLIAKLIHASHEEMH |
| S100A9\|Marmota monax\|KAF7460632.1 | S100A9 | F | L | aromatic | hphobic | ----------------------------------------------------------------------------------------------------------------------------------------------------------ECKKMARELSQMERSIETVIDTFHQYA-VRSGD-PDTLNQKEFKQLAKKELQNFLK----KENKDEKVINHILEDLDTNQDKQLSFEEFIMLIAKLTHASHEEMH |
| S100A9\|Urocitellus parryii\|XP_026268777.1 | S100A9 | F | L | aromatic | hphobic | ----------------------------------------------------------------------------------------------------------------------------------------------------------ECKKMAQELSQMERSIETVIDVFHQYS-VRLGD-RDTLNQKEFKQLVKTELKNYLK----KESKNEKVINHILEDLDTNLDKQLSFEEFIMLIAKLTHASHEEMH |
| S100A9\|Castor canadensis\|XP_020034902.1 | S100A9 | F | M | aromatic | hphobic | ----------------------------------------------------------------------------------------------------------------------------------------------------------ECKKMANQKSQLERSIDTIINVYHHYS-VRQGN-ADTLNKKEFKEMAKKELPNFLK----KENRDEKIIEDIMEDLDTNQDQQLNFEEFVILVARLTNASHEEMH |
| S100A9\|Choloepus didactylus\|XP_037670241.1 | S100A9 | F | M | aromatic | hphobic | --------------------------------------------------------------------------------------------------------------------------------------------------------------MAEELSPFEKSVETIINVFHQYS-ARLEN-PDTLNQNEFKKLVKKELANFLK----KQKKDDEAINHIMEDLDTNSDKQLNYEEYAILIARLVHASHEEMH |
| S100A9\|Trichechus manatus latirostris\|XP_004388696.1 | S100A9 | F | M | aromatic | hphobic | ------------------------------------------------------------------------------------------------------------------------------------------------------------------MSQMEHSIETIINVFHQYS-TRLGH-PDTLNQKEFKQLVQKELANFLK----NEKKNDAAIKDIMEDLDTNVDKQLSFEEFSILVGQLTEASHEKMH |
| S100A9\|Galeopterus variegatus\|XP_008565859.1 | S100A9 | F | L | aromatic | hphobic | --------------------------------------------------------------------------------------------------------------------------------------------------------------MACEMSLMESSIETVINVFHQYS-TRLGN-PDTLNQKEFKQLVKKELQNFLK----KEERDDKTINHILEDLDTNQDKELSFEEFIIMMARLTQASHVQMH |
| S100A9\|Molossus molossus\|XP_036129444.1 | S100A9 | F | M | aromatic | hphobic | -----------------------------------------------------------------------------------------------------------------------------------------------------------------MSSLMECSIETIINIFHQYS-TRLGH-PDRLNQREFKQMARKELPNFLK----KENRKPGAVEDIMEDLDTNGDKDLSFEEFAILVGRLTEASHEEMH |
| S100A9\|Miniopterus natalensis\|XP_016075607.1 | S100A9 | F | M | aromatic | hphobic | -----------------------------------------------------------------------------------------------------------------------------------------------------------------MSSLMECSIETIINIFHQYS-TRLGH-PDRLNQREFKQMVQKELPHFLK----KEKKNEAAINDIMEDLDTNGDKDLSFEEFIILVAKLTIASHEEMH |
| S100A9\|Eptesicus fuscus\|XP_008145594.1 | S100A9 | F | M | aromatic | hphobic | ----------------------------------------------------------------------------------------------------------------------------------------------------------------MSQSLMECSVDTIINIFHQYS-VRLGH-PDRLNQKEFSQMVKKELPTFLK----KERRNEALIKDMMEDLDTNGDKDLSFEEFITLVGRLTEASHEQMH |
| S100A9\|Myotis lucifugus\|XP_006095366.1 | S100A9 | F | L | aromatic | hphobic | ----------------------------------------------------------------------------------------------------------------------------------------------------------------MSQTLMECSVETIINIFHQYS-TRLGH-PDRLNQREFSQMVKKELPNFLK----KERRNEAAIRDILEDLDTSGDKDLSFEEFITLVGRLTEASHEEMH |
| S100A9\|Pipistrellus kuhlii\|XP_036308990.1 | S100A9 | F | L | aromatic | hphobic | ----------------------------------------------------------------------------------------------------------------------------------------------------------RGQEDEDKMSLLECSVNTIIDIFHQYS-VRLGH-PDRLNQREFSQLVKKELPNFLK----KEKKNENLIKDILEDLDTNGDKDLSFEEFITLVGTLTEASHEEMH |
| S100A9\|Microcebus murinus\|XP_012646736.1 | S100A9 | F | M | aromatic | hphobic | --------------------------------------------------------------------------------------------------------------------------------------------------------------MAREQSQLERAIETVINTFHHYS-VRQQP-PDSLSQKEFKQLVKKDLQNFLK----KEKRNDKAIEHIMEDLDTNVDKQLSFEEFIVLMARLTEASHEEMH |
| S100A9\|Otolemur garnettii\|XP_003802303.1 | S100A9 | F | M | aromatic | hphobic | --------------------------------------------------------------------------------------------------------------------------------------------------------------MATELSQLERSIETIINTFHHYS-VRQEP-PDSLNQKEFKQLVKKELLNFLK----KEKKNDKVINHIMEDLDTNEDKQLSFEEFIVLMARLTEASHEEMH |
| S100A9\|Camelus bactrianus\|XP_010952359.1 | S100A9 | F | M | aromatic | hphobic | --------------------------------------------------------------------------------------------------------------------------------------------------------------MADQMSQLECSIETIINIFHQYS-VRLGH-YDTLNKKEFKDLVKKELPNFLK----KQKKDDKVIEDIMQDMDTNEDETLNFQEFCMLVARLTVASHEEMH |
| S100A9\|Hipposideros armiger\|XP_019482834.1 | S100A9 | F | M | aromatic | hphobic | -----------------------------------------------------------------------------------------------------------------------------------------------------------------MSSLMECSFETIINIFHQYS-VRLGH-PDRLNQREFKQLVKKELPNFLK----KQKRNEAAINKIMVELDTDGDKDLTFEEFAVLIGRLTAASHEEMH |
| S100A9\|Rhinolophus ferrumequinum\|XP_032949639.1 | S100A9 | F | L | aromatic | hphobic | -----------------------------------------------------------------------------------------------------------------------------------------------------------------MSSLMECSLETIINIFHQYS-TRLEH-HDRLNQKEFKQMVRKELPNFLK----KQKRNEAAINHILEELDTNEDKELSFEEFSMLIARLTVASHEEMH |
| S100A9\|Pteropus giganteus\|XP_039705682.1 | S100A9 | F | L | aromatic | hphobic | -----------------------------------------------------------------------------------------------------------------------------------------------------------------MPSLMECSLETIINIFHQYS-TRLEH-PDRLNQREFKQLVKKELPNFLK----KQKKNDKAINHILEDLDTNGDKELSFEEFAILVARLTEASHEEMH |
| S100A9\|Pan paniscus\|XP_003817233.1 | S100A9 | F | M | aromatic | hphobic | --------------------------------------------------------------------------------------------------------------------------------------------------------------MTCKMSQLERNIETIINTFHQYS-VKLGH-PDTLNQGEFKELVQKDLQNFLK----KENKNEKVIEHIMEDLDTNADKQLSFEEFIMLMARLTWASHEKMH |
| S100A9\|Homo sapiens\|NP_002956.1 | S100A9 | F | M | aromatic | hphobic | --------------------------------------------------------------------------------------------------------------------------------------------------------------MTCKMSQLERNIETIINTFHQYS-VKLGH-PDTLNQGEFKELVRKDLQNFLK----KENKNEKVIEHIMEDLDTNADKQLSFEEFIMLMARLTWASHEKMH |
| S100A9\|Piliocolobus tephrosceles\|XP_023069543.1 | S100A9 | F | M | aromatic | hphobic | --------------------------------------------------------------------------------------------------------------------------------------------------------------MSCKMSQLERNIDTIIDTFHRYS-RQQGH-EDALNKKEFKDLVKTELQNFLK----KENKNDKIIDHIMEDLDTNADQQLSFEEFIMLMARLTWASHEKMH |
| S100A9\|Colobus angolensis palliatus\|XP_011783496.1 | S100A9 | F | M | aromatic | hphobic | --------------------------------------------------------------------------------------------------------------------------------------------------------------MSCKMSQLEGNIETIIDTFHEYS-TRLGH-KDTLNQKEFKELVKKELQNFLK----KENKNDKIIDHIMEDLDTNADKQLSFEEFIMLMARLTWASHEKMH |
| S100A9\|Chlorocebus sabaeus\|XP_007975283.2 | S100A9 | F | M | aromatic | hphobic | --------------------------------------------------------------------------------------------------------------------------------------------------------------MSCKMSQLERNIETIINTFHQYS-VRLGH-PDTLNRREFKQLVEKDLQNFLK----KEKKNDKIIDHIMEDLDTNADKQLSFEEFIMLMARLTWASHEKMH |
| S100A9\|Aotus nancymaae\|XP_012293733.1 | S100A9 | F | L | aromatic | hphobic | ------------------------------------------------------------------------------------------------------------------------------------------------------------------MSQLERSIGTIIDTFHHYS-VKLGH-PDTLNQREFKQLVQEELQNFLK----SKERNEQDINHILEDLDTNTDKQLSFEEFIMLMGRLTWASHEHMH |
| S100A9\|Saimiri boliviensis boliviensis\|XP_003942047.2 | S100A9 | F | L | aromatic | hphobic | ------------------------------------------------------------------------------------------------------------------------------------------------------------------MSQLECSIDTIIDTFHHYS-VRLGH-PDTLNQREFKQLVQKELPNFLK----KEKRNEQDINHILEDLDSNADKQLTFEEFIVLMGRLTWASHEHMH |
| S100A9\|Callithrix jacchus\|XP_008982671.1 | S100A9 | F | L | aromatic | hphobic | ------------------------------------------------------------------------------------------------------------------------------------------------------------------MSQLESSIETIINTFHHYS-VRLGH-PDALNQKEFKDLVQKELQNFLK----KEKRNEQDINHILEDLDTNADKQLTFEEFIMLMGRLTWASHEHMH |
| S100A9\|Ochotona princeps\|XP_004588975.1 | S100A9 | L | M | hphobic | hphobic | ------------------------------------------------------------------------------------------------------------------------------------------------------------MSCEWEMSQLEQSIDTIINTFHQYS-VRMGH-PDTLNQRELKQLVQKELPNFLK----MEARNEKAIKEIMEDLDTNQDKQLNFEEFVILVARLVHASHEEMH |
| S100A9\|Oryctolagus cuniculus\|XP_017200972.1 | S100A9 | F | M | aromatic | hphobic | --------------------------------------------------------------------------------------------------------------------------------------------------------------MSCGMSQLERSIDTIINVFHQYS-VRVGP-RDSLSQKEFKQLVQKELHNFLK----KEARDEKAINDIMEDLDTNQDKQLSFEEFVILMARLVHASHEEMH |
| S100A9\|Echinops telfairi\|XP_004717766.1 | S100A9 | F | M | aromatic | hphobic | --------------------------------------------------------------------------------------------------------------------------------------------------------------MACEMSQMEGSIQTIINVFHQYS-TRLGH-PDTLNQQEFKQLVVKDLANFLK----EEKRDKKAIQDIMEDLDTNSDKELSFEEYIILMGRLTHASHEEMH |
| S100A9\|Equus przewalskii\|XP_008537735.1 | S100A9 | F | L | aromatic | hphobic | ---------------------------------------------------------------------------------------------------------------------------------------------------------LECRKMAEL-SQMERDLETIINVFHQYS-VRLGH-PDTLNQKEFKQLVQKELANFLK----NEQKDEGAINHILEDLDTNEDKQLSFEEFIMLVVRLTLASHQKMH |
| S100A9\|Equus caballus\|XP_001493580.1 | S100A9 | F | L | aromatic | hphobic | --------------------------------------------------------------------------------------------------------------------------------------------------------------MAEL-SQMERDLETIINVFHQYS-VRLGH-PDTLNQKEFKQLVQKELANFLQ----NEQKDEGAINHILEDLDTNEDKQLSFEEFIMLVIRLTLASHQKMH |
| S100A9\|Ceratotherium simum simum\|XP_004436105.2 | S100A9 | F | L | aromatic | hphobic | ---------------------------------------------------------------------------------------------------------------------------------------------------------VECRKMAEL-SQMERSIETIINVFHQYS-VRLGH-PDTLNRKEFKQLVQKELANFLQ----NKKKDEKVINHILEDLDTNVDKQLSFEEFIVLVARLTVASHEKMH |
| S100A9\|Diceros bicornis minor\|KAF5924193.1 | S100A9 | F | M | aromatic | hphobic | ---------------------------------------------------------------------------------------------------------------------------------------------------------TECRKMAEL-SELERSIETIINVFHHYS-VRLGH-PDTLNRKEFKQLVQKELANFLK----SKKKDEKAINHIMEDLDTNADKELNFEEFSILIARLTEASHEKMH |
| S100A9\|Equus caballus\|XP_001494428.4 | S100A9 | F | M | aromatic | hphobic | ---------------------------------------------------------------------------------------------------------------------------------------------------------LECRKMAEL-SQMERDIETIINVFHQYS-VRLGH-PDTLNRKEFKQLVQKELANFLK----SKKKDEKAINHIMEDLDTNEDKQLSFEEFIILVARLTHASHEKMH |
| S100A9\|Elephantulus edwardii\|XP_006895953.1 | S100A9 | F | M | aromatic | hphobic | ------------------------------------------------------------------------------------------------------------------------------------------------------------------MSQLERNVETIINIFHQYS-TRLGH-PDTLNQSEFKQLVNKELANFLK----KENRNERVISDIMEDLDTNGDQQLNFEEYIILVARLTNASHEEMH |
| S100A9\|Manis pentadactyla\|XP_036778079.1 | S100A9 | F | M | aromatic | hphobic | --------------------------------------------------------------------------------------------------------------------------------------------------------------MEEQLSQMERHIETIINIFHQYS-VRLEP-RDSLSQKEFKQLVKKELPNFLK----KQKKNDRAINHIMEDLDTNQDKQLSFEEFSMLVGKLTVASHDKMH |
| S100A9\|Ovis aries\|KAG5215315.1 | S100A9 | L | M | hphobic | hphobic | --------------------------------------------------------------------------------------------------------------------------------------------------------------------MESSIETIINIFHQYS-IRLRD-PETLIRKELKQLVQKELPNFLK----KQHKDEEAINEIMEDLDTNQDKQLSFEEFIMLVARLTISA--QMH |
| S100A9\|Sus scrofa\|XP_013852489.2 | S100A9 | F | L | aromatic | hphobic | ---------------------------------------------------------------------------------------------------------------------------------------------------------TECRKMADQMSQMECSIETIINIFHQYS-VRLGN-RDTLNQKEFKQLVKKELPNFLK----KQKRDEKAINHILEDLDTNVDKQLSFEEFSMLVAKLTVASHEEMH |
| S100A9\|Capra hircus\|XP_013832703.2 | S100A9 | L | M | hphobic | hphobic | --------------------------------------------------------------------------------------------------------------------------------------------------------------MADQLSQMECSIETIINIFHQYS-IRLRD-PETLIQRELKQLVQKELPNFLK----KQNKDEEAINEIMEDLDTNQDKQLSFEEFIMLVARLTVASHEEMH |
| S100A9\|Odocoileus virginianus texanus\|XP_020763218.1 | S100A9 | F | M | aromatic | hphobic | --------------------------------------------------------------------------------------------------------------------------------------------------------------MADQMSQMECSIEAIINIFHQYS-VRLGH-PDTLMQKEFKQLVQKELPNFIK----KQKKDEKVINEIMEDLDTNQDKQLSFEEFIMLVAKLTVASHEEMH |
| S100A9\|Bubalus bubalis\|XP_006051754.1 | S100A9 | F | M | aromatic | hphobic | --------------------------------------------------------------------------------------------------------------------------------------------------------------MEDQMSQMECSIETIINIFHQYS-VRLGH-PDTLNQKEFKQLVQKELSNFLK----KQKKNDDAINEIMEDLDTNVDKQLSFEEFIMLVARLTVASHEEMH |
| S100A9\|Bos mutus\|XP_014338369.1 | S100A9 | F | M | aromatic | hphobic | --------------------------------------------------------------------------------------------------------------------------------------------------------------MEDQMSQMESSIETIINIFHQYS-VRLGH-PDTLIQKEFKQLVQKELPNFLK----KQKKNEDAINEIMEDLDTNVDKQLSFEEFIMLVARLTVASHEEMH |
| S100A9\|Bos taurus\|DAA31802.1 | S100A9 | F | M | aromatic | hphobic | --------------------------------------------------------------------------------------------------------------------------------------------------------------MEDKMSQMESSIETIINIFHQYS-VRLGH-YDTLIQKEFKQLVQKELPNFLK----KQKKNEAAINEIMEDLDTNVDKQLSFEEFIMLVARLTVASHEEMH |
| S100A9\|Chrysochloris asiatica\|XP_006861746.1 | S100A9 | F | L | aromatic | hphobic | --------------------------------------------------------------------------------------------------------------------------------------------------------------VAGKMSQMEKSIETIINVFHAYS-VRLGH-PDTLNQKEFKQLVKKELPNFLK----KEKKDDKLINHILEDLDTNGDKQLNFEEFSILVGRLTAASHEEMH |
| S100A9\|Suricata suricatta\|XP_029791997.1 | S100A9 | L | M | hphobic | hphobic | --------------------------------------------------------------------------------------------------------------------------------------------------------QAECRKMAEEQSQLERDIETIINIFHQYS-TRLGD-PDKLNQRELKQLVEKELRNFLK----KPKKNDNAIKKIMEDLDTNGDKELSFEEFTMLVARLTVASHEEMH |
| S100A9\|Orycteropus afer afer\|XP_007946782.1 | S100A9 | F | L | aromatic | hphobic | ------------------------------------------------------------------------------------------------------------------------------------------------------------------MSQLECNIETIINIFHQYS-VRLGH-PDTLNQREFKQLVKKELANFLK----KQKRDDKAINHILEDLDTNGDKQLNFEEFVILVGRLTHASHEEMH |
| S100A9\|Mustela erminea\|XP_032174708.1 | S100A9 | L | M | hphobic | hphobic | --------------------------------------------------------------------------------------------------------------------------------------------------------VQEYRKMADQMSQLERSIETIINIFHQYS-VHMVH-PDMLNQKELKQLVKKELPNFLK----KQ-KNDNAINKIMEDLDTNGDKQLNFEEFSILVGRLTMASHEEMH |
| S100A9\|Gulo gulo\|VCW76681.1 | S100A9 | L | L | hphobic | hphobic | -----------------------------------------------------PFLALTVQLRVATLLTCEAIFARELSIWPGAAYKCWAHTAPSHPSVWL-----------------------------------------------VQEYRKMADQMSQLEGSIETIINIFHQYS-VRLGH-PDMLEQKELKQLVKKELPNFLK----KQKKNDKAINKILEDLDTNGDKQLNFEEFSILVGRLTVASHEEMH |
| S100A9\|Odobenus rosmarus divergens\|XP_004403099.2 | S100A9 | L | M | hphobic | hphobic | -----------------------------------------------------HFLALIVQLRVTTLLTCEAIFPRELSIWPEAAYKCWTHTVLSNPSVWLL----------------------------------------------DLECSKMADQMSQLECSIETIINIFHQYS-VRLQH-PDKLNQKELKQLVKKELPNFLK----RQKKNDYAINKIMEDLDTNGDKELDFEEFSILVGRLTLASHEEMH |
| S100A9\|Mirounga leonina\|XP_034865489.1 | S100A9 | L | M | hphobic | hphobic | --------------------------------------------------------------------------------------------------------------------------------------------------------------MADEMSQLECSIETIINIFHQYS-VRLGH-PDTLNQKELKQLVKKELPNFLK----KQKKNDNAINKIMEDLDTNGDKQLNFEEFSILVGRLTVASHEEMH |
| S100A9\|Crocuta crocuta\|KAF0880560.1 | S100A9 | L | M | hphobic | hphobic | -----------------------------------------------------PFLALTAQLRVTTLLTCEAIFPRELSIRPGAAYKCWTHTVLSNPFVWLL---------------------------------------------CRQSGRKMADELSQLECNIETIINIFHQYS-VRLGH-PDKLNQRELKQMVKKELPNFLK----KQKKNDHAINKIMEDLDTNGDKELNFEEFSMLVARLTVASHEEMH |
| S100A9\|Lynx pardinus\|VFV19105.1 | S100A9 | M | M | hphobic | hphobic | -----------------------------------------------------PFLALTAQLRVTTLLTCEAIFPRELTIRPGAAYKCWTHGVLSNPVVWLL---------------------------------------------GRQECRKMADQLSQMECSIETIINIFHQYS-VRLGH-PDKLNQKEMKQLVKKELPNFLK----KQKKNDNAINKIMEDLDTNGDKELDFEEFSILVARLTVASHEEMH |
| MRP-126\|Anolis carolinensis\|XP_016854246.1 | MRP-126 | T | F | polar | aromatic | ------------------------------------------------------------------------------------------------------------------------------------------------------------------MSQMENAIDIIINVFHQYS-IQKPH-PDKLSKGETKNLIQKELAEFLK-----NQVNPLVVEEMFKQLDQDQDKELSFQEFMQLVVQVTVATHNNLH |
| MRP-126\|Pogona vitticeps\|XP_020666408.1 | MRP-126 | T | F | polar | aromatic | ------------------------------------------------------------------------------------------------------------------------------------------------------------------MTQLEIAFETVINVFHQYS-VLKPH-PDKLSKGETKQLMEKELPEFLK-----KQVNPKGIEELFKDLDTNKDQDITFKEFMGMATQVLIATHTNLH |
| MRP-126\|Varanus komodoensis\|KAF7235497.1 | MRP-126 | L | F | hphobic | aromatic | ------------------------------------------------------------------------------------------------------------------------------------------------------------------MSPLETALDTVINVFHQYS-THKPH-PDKLNNGELKQLIEKELAEFLK-----NQVNPTSIEEMFKHLDQNKDQEISFKEFMVMIGQVTEATHEKLH |
| MRP-126\|Anolis carolinensis\|XP_003229667.1 | MRP-126 | M | F | hphobic | aromatic | ---------------------------------------------------------------------------------------------------------------------------------------------------MS----PTQ-----MESCMETVVNIFHQYS-VRQDH-FDKLSKKEMSQLVHKQFPNWLK-----EQKNPKAIEELFKQLDQDQDNQLSFGEFMVFICKLTIATHEHIH |

| | | | | | | |
|---|---|---|---|---|---|---|
| MRP-126\|Pogona vitticeps\|XP_020666411.1 | MRP-126 | L | F | hphobic | aromatic | ----------MS----QTQ-----LEACIETQINVFHQYS-VRHGH-FDTLSAKELEQLIQKQLPNFLK-----DSKNPKAIEKLFKELDQDKDKQISFGEYAAFLSRVAIASHEHIH |
| S100A9\|Varanus komodoensis\|KAF7235505.1 | S100A9 | L | F | hphobic | aromatic | ----------MA----QTQ-----LEKCFETVINIFHQYS-VRVGH-FDTLTKGELRQLIEKELPTFLK-----EQKNPQAIEKIFTDLDKNKDKQISFGEFMDLVVKVVIAAHEHIH |
| MRP-126\|Dromaius novaehollandiae\|XP_025973891.1 | MRP-126 | L | F | hphobic | aromatic | ----------MPLRKRLHSGPSPSHRFVKMSQTCQTPQGQLSKLEIAINSIIDVFHQYS-RQEGH-MDTLTKVELKLLIQKQLANYLQ-----HVTNQVSIDQIFKDLDTNKDHQLSFGEVMLLIIRVTIATHEHLH |
| S100A9\|Apteryx rowi\|XP_025936655.1 | S100A9 | L | F | hphobic | aromatic | ----------MPLSERLHSSLSPSRRFVKMSQACQTPRGPLSELERSIDTIIDVFHQYS-RREGD-KDTLTNAELKLLIERQLVNYLK-----HVKNQVSIDQIFKDLDTNKDHQLSFGEVMLLIIRVTIATHDHLH |
| MRP-126\|Nothoprocta ornata\|NWY07803.1 | MRP-126 | M | F | hphobic | aromatic | VNRDRELSFEEFCVVLAKLADDAHRHVLVMLE----------PSFSLQTCQTAQGPLSELEKAIDTIIDVFHQYS-RREGD-KDTLSKMEMKLLVEKQLANYLR-----HVKDQSSLDQIFKDLDTDKNQQLSFGEVMLLIVRVTIATHKHLH |
| S100A9\|Eudromia elegans\|NXA43986.1 | S100A9 | M | F | hphobic | aromatic | INRDRELSFEEFCIVLAKLADDAHRISHGSER----------PSFSLQTCQTVQGPLSELEKSIDTIIDVFHQYS-RREGD-KDTLTKAEMKLLVEKQLANYLR-----HVKSQSSLDQIFKDLDVNKDQQLSFGEVMLLLVRVTIATHEHLH |
| S100A9\|Crypturellus soui\|NWI22021.1 | S100A9 | M | F | hphobic | aromatic | ----------CQTVQGPLSELEKAIDTIIDVFHQYS-RREGD-KDTLTKAEMKLLLEKQLANYLR-----NVKTQSSLDQIFKDLDVNKNQQLSFGEVMMLLVRVTIATHEHLH |
| S100A9\|Tinamus guttatus\|XP_010214781.1 | S100A9 | M | F | hphobic | aromatic | ----------MSQTCQTAQAPLSELEKSIDTIIDVFHQYS-RREGD-KDTLTRAEMKLLLEKQLANYLR-----HVKNQSSLDQIFKDLDTNKNQQLSFGEVMMLIVRVTIATHEHLH |
| S100A9\|Struthio camelus australis\|XP_009674396.1 | S100A9 | L | F | hphobic | aromatic | ----------MSQSCQTPQGPISDLERAIDTIIDVFHQYS-RREGD-KDTLTKVELKLMIEKQLANYLK-----HVKNQISIDQIFKDLDTNKDHQLSFGEVMLLIIRVTVATHEHLH |
| MRP-126\|Meleagris gallopavo\|XP_010725281.1 | MRP-126 | L | F | hphobic | aromatic | ----------NMSKGCQTQGPLSELEKAIDVIIDVFHQFS-RREGD-KDTLTKKELKLLIEKQLANYLK-----HVKNQASIDQIFKDLDGNKDQQLSFGEVMLLIMRVTAATHEHLH |
| MRP-126\|Gallus gallus\|NP_001292080.1 | MRP-126 | L | F | hphobic | aromatic | ----------MSKGCQTQGPLSELEKAIDVIIDVFHQYS-RREGD-KDTLTRKELKLLIEKQLANYLK-----HVKNQVSIDQIFKDLDNNKDQQLSFGEVMLLIIRVTVATHEHLH |
| S100A9\|Anseranas semipalmata\|NXI72433.1 | S100A9 | L | F | hphobic | aromatic | LNKDRELSFEEFTIVLAKVADDAHRIIHKDDR----------LYSFLFQGCQTQGPLSELEKSIDVIIDVFHQYS-RREGD-KDTLTRRELKLLIEKQLANYLK-----HVKNQVSIDQIFKDLDMNKDEQLSFGEVMLLIIRVTVATHEHLH |
| S100A9\|Cygnus atratus\|XP_035423362.1 | S100A9 | L | F | hphobic | aromatic | ----------FLFQGCQTQGPLSELEKSIDVIIDVFHQYS-RREGD-KDTLTRRELKLLIEKQLVNYLK-----HVKNQVSIDQIFKDLDTNKDEQLSFGEVMLLIIRVTVATHEHLH |
| S100A9\|Cygnus olor\|XP_040395106.1 | S100A9 | L | F | hphobic | aromatic | ----------NMSKGCQTQGPLSELEKSIDVIIDVFHQYS-RREGD-KDTLTRRELKLLIEKQLVNYLK-----HVKNQVSIDQIFKDLDTNKDEQLSFGEVMLLIIRVTVATHEHLH |
| MRP-126\|Phalacrocorax carbo\|XP_009505423.1 | MRP-126 | L | M | hphobic | hphobic | ----------PRGASPRHTPQGQEQLSELEKSMDVIIDIFHKYS-RQEGH-SDTLTKNELKLLIQSQLANYLR-----HVKNKATIDQIMNDLDINKDQQISFSETMLLITRVTIATHEHLH |
| S100A9\|Tauraco erythrolophus\|XP_009984260.1 | S100A9 | L | M | hphobic | hphobic | ----------MESGIECTSSKFAEDINRVIDVFHQYS-VQEGH-RDTLSKKELKLLIEKQLANYLK-----HVKNRGTIDEIMKDLDVNKDQQVSFSEVMMLITRVTIATHEHLH |
| MRP-126\|Orthonyx spaldingii\|NXC06591.1 | MRP-126 | L | M | hphobic | hphobic | LNKDKQVSFEEFTIVLTKLADDAHRISHGSDR----------CGPDKDTSQTRLSELEKSIDVIIDVFHHYS-RREGD-KDTLTKRELKLLITEQLANYLQ-----HVKNKQTIDEIMKDLDTNQDAQLSFEETMVLVTRVTCATHEDLH |
| MRP-126\|Pelecanus crispus\|XP_009480622.1 | MRP-126 | L | M | hphobic | hphobic | ----------TTTALLLFSDLEKAMNEIIDVFHHYS-RREGD-KDTLTKRELKLLIEEQLRNYLQ-----HVKNKATIDEIMKDLDINKDEQLSFCEVMLLITRVTIATHEHLH |
| MRP-126\|Rhinopomastus cyanomelas\|NXO04703.1 | MRP-126 | L | M | hphobic | hphobic | LNKDKELSFEEFTIVLAKLTNDAHHIIHKEDR----------WTGLWAPWCGQLSELEKSMDVIIDVFHHYS-RREGD-KDTLTKKELKLMIESQLANYLN-----HVKDKATIDEIMKDLDVNKDAQISFNEMMLLIVRVTIATHDHLH |
| MRP-126\|Amazona aestiva\|KQL60522.1 | MRP-126 | L | M | hphobic | hphobic | ----------SSIIHPTFGTXQTQEPLSPLEKSMDVIIDVFHQYS-RREGD-RDTLTKTELKLLIEKQLANYLK-----HVKCSATIDEIMKDLDMNRDAQLSFSEVMLLITRVTIATHEHLH |
| MRP-126\|Nyctibius grandis\|NXQ82064.1 | MRP-126 | L | M | hphobic | hphobic | INKDRDLTFEEFIIVLSKVTDDAHRLIHGHDR----------SVGQWDGFGCSRSARTQEPLSELERAMDVIIDVFHQYS-RREGD-KDTLTKKELKLMIEKQLVNYLK-----HVKNKATVDEIMKDLDINKDEELNFCETMLLITRVTIATHEHLH |
| S100A9\|Bucco capensis\|NXH18374.1 | S100A9 | L | M | hphobic | hphobic | INQDKQLSFEEFCIVLAKLTDDAHRAMHGHER----------CGESFLLQTPQSQEHLSELEKAIDVIIDVFHQYS-RREGD-RDTLSKSELKLLIEKQLANYLK-----HVKNKATLDEIMNDLDVKKDAQLSFCEAMLLIIRVTIATHEHLH |
| MRP-126\|Pipra filicauda\|XP_027583069.2 | MRP-126 | L | F | hphobic | aromatic | ----------MA----------DFHTHCGSCCYLVQPFLFQSSQSQEQLSELEKAMDVIIDVFHHYS-RREGD-RDTLTKKELKLLIEKHLANYLK-----HVKNKATIDEIFKDLDINKDSQVSFCEMMLLVVRVTSATHEHLH |
| MRP-126\|Megadyptes antipodes antipodes\|KAF1466865.1 | MRP-126 | L | M | hphobic | hphobic | LNQDRELTFEEFTIVLAKLTDDAHRISHGDDR----------LQLLLFQTPQAREPRSELEKAMDIIIDVFHQYS-RREGD-RDTLTKKELKLLIEQQLVNYLK-----LVKNRATIDEIMKDLDINKDAQISFSEVMLLITRVTIAAHEYLH |
| MRP-126\|Pygoscelis adeliae\|XP_009320488.1 | MRP-126 | L | M | hphobic | hphobic | ----------MSKTPQTREQRSELEKAMDIIIDVFHQYS-RREGN-RDTLTKKELKLLIEQQLVNYLK-----LVKDKATIDEIMKDLDINKDAQISFSEVMLLIIRVTIAAHEHLH |
| MRP-126\|Pachyramphus minor\|NWS21030.1 | MRP-126 | L | F | hphobic | aromatic | LNKDKELSFEEFSIVLAKVADDAHRITHRSDR----------YFHIHCGSCCYLMQPFLFQISQSQEPLSELEKAMDVIIDVFHQYS-RREGD-RDTLTKKELKLLIEKQLVNYLK-----HVKNKATIDEIFKDLDVNKDAQINFCETMLLIVRVTIATHEHLH |
| MRP-126\|Piprites chloris\|NXK43174.1 | MRP-126 | L | F | hphobic | aromatic | LNKDKELTFEEFSIVLAKVADDAHRIIHRSDR----------CGPDHSQSQEPLSELEKAMDVIIDVFHQYS-RREGD-RDTLTKKELKLLIDKQLVNYLK-----HVKNKATIDQIFKDLDVNKDAQVGFCEMMQLVVRVTIATHEHLH |
| MRP-126\|Oxyruncus cristatus\|NXM33156.1 | MRP-126 | L | W | hphobic | aromatic | LNKDKEVSFEEFCIVLAKVADDAHRISHRSDR----------CGPDHSQSQAPLSELEKAMDVIIDVFHQYS-RREGD-RDTLTKKELKLFIEKQFVNYLK-----HVKSQATIDQIWKDLDANKDAQISFGEMMLLVVRVTIATHEHLH |
| MRP-126\|Falco rusticolus\|XP_037227195.1 | MRP-126 | L | M | hphobic | hphobic | ----------MSKTSQTQEPLSELEKAMNVIIDVFHQYS-RREGD-RDTLTKTELKLMIEKQLANYLR-----NVKDKATIDEIMKDLDINKDAQLSFCEVMLLIIRVTIATHEHLH |
| MRP-126\|Oriolus oriolus\|NXO15483.1 | MRP-126 | L | M | hphobic | hphobic | LNKDSEVSFEEFTIVLSKVADDAHRISHDSDR----------CGPDGSQTHEHLSELEKAMDVIIDVFHQYS-RREGD-RDTLTKQELKLMIEKQLANYLK-----HVKNKATIDQIMKDLDLNKDAQISFGEMMLLVTRVTCATHEHLH |
| MRP-126\|Catharus ustulatus\|XP_032938530.1 | MRP-126 | L | M | hphobic | hphobic | ----------MFYSHFLPQSQQSQAPLSELERAMDSIIDVFHQYS-RREGD-RDTLSRGELKLLIEKQLSNYLK-----HVKNKATIDQIMKDLDVNKDAQLSFGEVMLLVTRVTCATHEHLH |
| MRP-126\|Irena cyanogastra\|NXI20392.1 | MRP-126 | L | M | hphobic | hphobic | ----------QSQEPLSQLEKAMDTIIDVFHLYS-RREGD-RDTLTKQELKLLIEKQLANYLK-----HVKSKATIDQIMKDLDLNKDAQLSFGEVMLLIVRVTCATHEHLH |
| MRP-126\|Sitta europaea\|NXO87048.1 | MRP-126 | L | M | hphobic | hphobic | LNQDKELSFEEFAIVLSKLADDAHRISHGSDR----------CAASVCPQVQQSQEPLSELEKAMDVIIDVFHQYS-RREGD-RDTLTKNELKLMIEKQLANYLK-----HVKNKATIEQIMKDLDVNKDAQISFGEMMLLVTRVTCATHEHLH |
| MRP-126\|Galbula dea\|NXI41603.1 | MRP-126 | L | M | hphobic | hphobic | INKDKDLTFEEFSIFLAKVTDDAHRIIHNEEL----------SSFLFQTSQTQEALSELEKAMDVIIDVFHQYS-RREGD-RDTLTKSELKLLIEKQLANYLK-----HVKNKATIDEIMKDLDLNKDAQLSFGETMLLIIRVTIATHEHLH |
| MRP-126\|Zosterops hypoxanthus\|NXR33221.1 | MRP-126 | L | M | hphobic | hphobic | ----------PQSQQSQEPLSDLEKAMNVIIDVFHQYS-RREGD-KDTLTKKELKLLIEKQLANYLK-----HVKNKPSLDQIMKDLDVNKDAQVSFGETMLLVTRVTCATHEHLH |
| MRP-126\|Cyanistes caeruleus\|XP_023797607.1 | MRP-126 | L | M | hphobic | hphobic | ----------SAGSWLLGEGQQSQEPLSELEKAMDVIIDVFHQYS-RREGD-RDTLSKKELKLMIEKQLANYLK-----HVKNKATIDQIMKDLDVNKDAQISFGEMMLLVTRVTCATHEHLH |
| MRP-126\|Menura novaehollandiae\|NXE99101.1 | MRP-126 | L | M | hphobic | hphobic | ----------QTQEPLSELEKAMDVIIDVFHQYS-RREGD-RDTLTKRELKLLIEKQLGNYLK-----HVKNKTTIDQIMKDLDLNKDAQLNFGETMLLVIRVTCATHEHLH |
| MRP-126\|Leptocoma aspasia\|NXL78334.1 | MRP-126 | L | M | hphobic | hphobic | ----------QSQEHLSELEKAMDAIIDVFHHHS-RREGD-KDTLTKKELKLLIDTQLANYLK-----HVKNKATIDQIMKDLDLNKDAQISFGEMMLLVTRVTCATHEHLH |
| MRP-126\|Loxia leucoptera\|NXG97562.1 | MRP-126 | L | M | hphobic | hphobic | LNKDKEVSFEEFTMVLSKLADDAHRISHGSER----------CGPDGQQSQQPLSELEKAMDAIIDVFHQYS-RREGD-RDTLTKKELKLLLEKQLANYLK-----HVKSQATIDQIMKDLDVNKDAQISFGEMMLLVTRVTCATHEHLH |
| MRP-126\|Parus major\|XP_015472318.1 | MRP-126 | L | M | hphobic | hphobic | ----------IWFLPQGQQSQEPLSELEKAMDVIIDVFHQYS-RREGD-RDTLSKKELKLLIEKQLANYLK-----HVKNKATIDQIMKDLDVNKDAQISFGEMMLLVTRVTCATHEHLH |
| MRP-126\|Chloebia gouldiae\|RLV84402.1 | MRP-126 | L | M | hphobic | hphobic | ----------GWVFLQAQQSQEPLSELEKSMDVIIDVFHQYS-RREGD-RDTLTKKELKLLIEKQLANYLK-----HVKSKATIDQIMKDLDVNKDAQISFGEMMLLVTRVTCATHEHLH |
| MRP-126\|Lonchura striata domestica\|OWK52003.1 | MRP-126 | L | M | hphobic | hphobic | ----------KMSKAQQSQEPLSELEKAMDVIIDVFHQYS-RREGD-RDTLTKKELKLLIDKQLANYLK-----HVKSKATIDQIMKDLDVNKDAQLSFGEVMLLVTRVTCATHEHLH |
| MRP-126\|Eurystomus gularis\|NXW62297.1 | MRP-126 | L | M | hphobic | hphobic | INKDKDLNFEEFTIVLSKLTDDAHRITHSDDR----------CTPDKDTSQTQEPLSELEKAMDVIIDVFHQYS-RQEGD-RDTLTKGELKLMIEKQLVNYLK-----HVKNKATIDEIMKDLDFNKDEQLSFNEVMLLIIRVTIATHDHLH |
| MRP-126\|Oreotrochilus melanogaster\|NXU79994.1 | MRP-126 | L | M | hphobic | hphobic | LNKDKELTFEEFVIVLAKVTDDAHRISHGQDR----------WWLLILSFLFQTSQTQEPLSDLEKAMNVIIDVFHQYS-RREGD-RDTLTKKELKLLIEKQLVNYLK-----NVKNKATIDEIMKDLDINKDAQISFCEVMLLITRVTIATHEHLH |
| MRP-126\|Calyptomena viridis\|NWI56398.1 | MRP-126 | L | M | hphobic | hphobic | INGDKELTFEEFVIILCKVTDDAHRISHNSDR----------CRAGGSWCYLVQSFLFQTSQSQGPLSELEKSMNVLIDVFHQYS-RREGD-RDTLTKKELKLLIEKQLVNYLQ-----HVKNKATIDEIMKDLDINKDAQLSFCEVMLLITRVTIATHEHLH |
| MRP-126\|Chaetura pelagica\|XP_009999847.1 | MRP-126 | L | M | hphobic | hphobic | INHDKEVSFEEFTMVLAKAALATQAVEQQDQE----------MLVGAQDHVVMWVHQVVLWQTSQTQEALSELERSMDVIIDVFHQYS-RREGH-RDTLTKRELKLLIEKQLANYLK-----HVKSQATIDEIMKDLDINRDAQLSFSEVMLLITRVTIATHEHLH |
| MRP-126\|Nyctiprogne leucopyga\|NXW51825.1 | MRP-126 | L | M | hphobic | hphobic | LNKDKQLTFEEFSIVLSKLTDDAHRISHGDDR----------SCQSQEQLSELEKAMDLIIDVFHHYS-RREGD-RDTLTKKELKLLIEKQLANYLK-----HVNNKVTIDEIMKDLDVNRDAQISFPEMMMLITRVTTATHEHLH |
| MRP-126\|Sakesphorus luctuosus\|NXG09850.1 | MRP-126 | L | N | hphobic | polar | ----------WWYFWGPSFLFQTSQSQAPLSELEKAIDVIIDVFHQYS-RREGD-RDTLTKAELKLLIEKQMVNYLK-----HVKNQTTIDEINKDLDINKDAQISFCEMMLLITRVTIATHDHLH |
| MRP-126\|Fregetta grallaria\|NXW09654.1 | MRP-126 | L | M | hphobic | hphobic | LNKDKELSFEEFTIVLAKVTDDAHRINHGDNR----------GPHGASTQEPLSELEKAMDVIIDVFHQYS-RQEGH-RDTLTKNELKLLIEKQLANYLK-----HVKNKATIDEIMKDLDVNKDEQISFCETMLLIVRVTVATHEHLH |
| MRP-126\|Rhynochetos jubatus\|NWW87098.1 | MRP-126 | L | M | hphobic | hphobic | INKDKELTFEEFTVVLAKLTDDAHRIIHNDPR----------CHPTSQIQEPLSELEKAMDVIIDVFHQYS-RREGD-RDTLSKNELKLLIEKQLVNYLK-----HVKNKATIDEIMRDLDINKDAQISFSETMLLVTRVTIATHEHLH |
| MRP-126\|Aquila chrysaetos chrysaetos\|XP_029879656.2 | MRP-126 | L | M | hphobic | hphobic | ----------MSKTSQIQEPLSELEKAMDVIIDVFHQYS-RREGD-RDTLTKNELKLLIEKQLVNYLK-----HVKNKATIDEIMKDLDINKDAQISFCEVMLLIVRVTIATHEHLH |
| S100A9\|Melopsittacus undulatus\|XP_030910547.2 | S100A9 | L | M | hphobic | hphobic | ----------MSKTSQSQEPLSQLEKAMDVIIDVFHQYS-RREGD-RDTLTKAELKLLIEKQLANYLK-----HVTSTATIEEIMKDLDRNKDAQLSFSEVMLLIVRVTIATHEHLH |
| S100A9\|Buceros rhinoceros silvestris\|XP_010142943.1 | S100A9 | L | M | hphobic | hphobic | LDNDREVSFEEFTIVLAKLTSDTYYILHEEDR----------SSCHTPQTQEHLSELEKAIDVIIDVFHQYS-RREGD-RDTLTKSELKLLIEKQLANYLK-----HVKNQATIDEIMKDLDLNKDQQLSFNETMLLIIRVTVATHDHLH |
| MRP-126\|Ceyx cyanopectus\|NXY92371.1 | MRP-126 | L | M | hphobic | hphobic | ----------QTQGPLSELEKAMDVIIDVFHQYS-RREGD-RDTLTKNELKLLIEKQLANYLK-----HVNNQATVDQIMTDLDMNKDQQLSFNEAMLIVIRVTIATHDHLH |
| MRP-126\|Zapornia atra\|NXT78576.1 | MRP-126 | L | M | hphobic | hphobic | LNRDRELSFDEFTIVLSKLADDAHRISHGQDR----------SQEPLSELEKSMDVIIDVFHQYS-RREGD-RDTLTKKELKLLIEQQLVNYLR-----HVKNKATIDEIMKDLDINKDAQLSFCEVMLLITRVTIATHEHLH |
| MRP-126\|Corythaeola cristata\|NXC16638.1 | MRP-126 | L | M | hphobic | hphobic | ----------QTQEPLSELEKAMNNIIDVFHNYS-VQEGH-RDTLTKKELKLLIEKQLANYLK-----HVKNKGTIDEIMTDLDLNKDAQLSFGEVMLLITRVTIATHEHLH |
| MRP-126\|Caloenas nicobarica\|NWX11064.1 | MRP-126 | L | M | hphobic | hphobic | LNKDRQLTFEEFTIVLAKLTDDAHRLSHGHNPGQWDTFGCAGPR----------GARIQQIQEPLSELEKSMDVIIDVFHQYS-RREGD-RDTLTKKELKLLIEKQLANYLK-----HVKSKATIDEIMKDLDINRDAQISFCEVMLLITRVTIATHEHLH |
| MRP-126\|Cochlearius cochlearius\|NXE84340.1 | MRP-126 | L | M | hphobic | hphobic | INKDRKLTFEEFTIVLGKLTNDAHCISHNHNR----------CTPTSQIQGPLSELEKAMDVIIDVFHQYS-RQEGD-RDTLTKKELKLLIEKQLANYLQ-----HVKNKATIDEIMKDLDINRDAQISFSEVMLLIVRVTIATHEHLH |
| MRP-126\|Egretta garzetta\|XP_035746619.1 | MRP-126 | L | M | hphobic | hphobic | ----------MSKTSQIQGPLSELEKAMDVIIDVFHQYS-RQEGD-RDTLTKKELHLLIEKQLANYLQ-----HVENKATIDEIMKDLDINKNMQISFSEAMLLIIRVTVATHKHLH |
| MRP-126\|Acanthisitta chloris\|XP_009079040.1 | MRP-126 | L | M | hphobic | hphobic | ----------DLATAQSQGQLSELEKSMDVIIDVFHQYS-RREGD-RDTLTKKELKLLIDKQLANYLK-----HVRNKTTIDEIMRDLDTNRDHQISFSEVMLLITRVTIATHEHLH |
| MRP-126\|Serilophus lunatus\|NXM75952.1 | MRP-126 | L | M | hphobic | hphobic | LNKDREMSFEEFTIVLSKLADDMHRVSHHSDR----------YFHTYRGSWCHLVQSFLFQTSQSQGSLSELEKAMDVIIDVFHHYS-RREGD-RDTLTKKELKLLIEKQLANYLQ-----HVKNKATIDEIMKDLDINKDAQISFSEVMLLITRVTIATHEHLH |
| MRP-126\|Furnarius figulus\|NWR95575.1 | MRP-126 | L | M | hphobic | hphobic | LNKDKELSFEEFTVVLSKLADDAHRISHHSDR----------YLHISRGSWWHLGQSFLFQTSQSQEQLSELEKAMDIIIDVFHQYS-RREGD-RDTLTKKELKLLIEKQLVNYLK-----HVKNKATLDEIMKDLDINKDAQISFCEVMLLITRVTIATHEHLH |
| MRP-126\|Chloroceryle aenea\|NXI49743.1 | MRP-126 | L | M | hphobic | hphobic | LNKDKDLSFEEFTIVLAKVADDAHRIIHGEDR----------WCWLGRWSQGPLSELEKAMDAIIDVFHQYS-RREGD-RDTLTKNELKLLIEKQLANYLK-----HVKKQATIDQIMKDLDTNKDAQLSFNEVMLLIISVTIATHDHLH |
| MRP-126\|Columba livia\|XP_005512844.2 | MRP-126 | L | M | hphobic | hphobic | ----------MSKSSQTQEPLSELEKSMDVIIDVFHQYS-RREGD-RDTLTKKELKLLIEKQLANYLK-----HVKSKATIDEIMKDLDINRDAQISFCEVMLLITRVTIATHEHLH |
| MRP-126\|Ciconia maguari\|NXJ34161.1 | MRP-126 | L | M | hphobic | hphobic | LNKDREVAFEEFTMVLAKLSDDAHRLMHHQER----------SCQIQEPLSELEKAMDVVIDVFHQYS-RREGD-RDTLTKNELKLLIEKQLVNYLK-----HVKNKATIDEIMKDLDINKDAQISFCEVMLLITRVTIATHEHLH |
| MRP-126\|Psophia crepitans\|NXI98983.1 | MRP-126 | L | M | hphobic | hphobic | INKDKVLTFEEFTIVLAKLADDAHRISHRDDR----------TFGTSQIQEPLSELEKSMDVIIDVFHQYS-RREGD-RDTLTKRELKLLIEKQLVNYLR-----HVKNKTTIDEIMKDLDINKDAQISFCEVMLLITRVTIATHEHLH |
| MRP-126\|Nipponia nippon\|XP_009469672.1 | MRP-126 | L | M | hphobic | hphobic | ----------CSSQIKDPLSELEKAMDVIIDVFHQYS-RREGD-RDTLTKRELKLLIEKQLVNYLR-----HVKNKATIDEIMKDLDINKDAQISFCEVMLLITRVTIATHEHLH |
| S100A9\|Colius striatus\|XP_010194785.1 | S100A9 | L | M | hphobic | hphobic | ----------MSKTPQSQEQLSELEKAMDVIIDVFHQYS-RREGD-RDTLTKSELKLLVEKQLANYLK----KHVKSKATIDEIMKDLDFNKDAQLSFSEVILLITRVTIATHEHLH |
| MRP-126\|Eurypyga helias\|XP_010146505.1 | MRP-126 | L | M | hphobic | hphobic | ----------DSGIKISQSQEPLSELEKSMDAIIDVFHQYS-RREGD-RDTLSKMELKLLIEKQLANYLR-----HVKNKATIDEIMKDLDINKDAQISFCEMMLLITRVTIATHEHLH |
| MRP-126\|Gavia stellata\|XP_009814804.1 | MRP-126 | L | M | hphobic | hphobic | ----------LSNAQTTQTQEQLSELEKAMDVIIDVFHQYS-RREGD-IDTLTKKELKLLIDKQLVNYLK-----HVRNKTTIDEIMKDLDVNKDAQISFSEMMLLVTRVTIATHEHLH |
| MRP-126\|Limosa lapponica baueri\|PKU30472.1 | MRP-126 | L | M | hphobic | hphobic | ----------QTPQSQEQLSELEKAMNVIIDVFHQYS-RREGD-RDTLTKSELKLLIEKQLLNYLR-----HVRNKATIDEIMKDLDINKDAQISFCEVMLLITRVTIATHEHLH |
| MRP-126\|Leptosomus discolor\|XP_009960271.1 | MRP-126 | L | M | hphobic | hphobic | ----------MSKTSQTQGPLSELERAMDVIIDVFHQYS-RREGD-RDTLTKAELKLLIEKQLANYLR-----HVNNKTTIDQIMKDLDINKDAQISFCETMLLVTRVTIATHEHLH |
| MRP-126\|Podargus strigoides\|NXX20266.1 | MRP-126 | L | M | hphobic | hphobic | LNKDKEVSFEEFTIVLAKVTDDAHRIIHGDER----------CLWLRCGEWVRATSQTQEPLSELEKSMDVIIDVFHQYS-RREGD-RDTLTKKELKLLIEQQLVNYLK-----HVKNKATIDEIMKDLDINKDAQLSFCEVMLLITRVTIATHEHLH |
| MRP-126\|Oceanodroma tethys\|NXH66189.1 | MRP-126 | L | M | hphobic | hphobic | LNEDKELTFEEFTIVLAKVTDDAHRISHGDDR------CCPSSRGSC----------SCQTQEQLSELEKAMDVIIDVFHQYS-RREGH-RDTLTKNELKLLIEKQLVNYLR-----HVKNKATIDEIMKDLDANKDDQLSFCETMLLITRVTIATHEHLH |
| MRP-126\|Pelecanoides urinatrix\|NXT43348.1 | MRP-126 | L | M | hphobic | hphobic | LNKDKELSFEEFTIVLAKVTDDAHRINHGDDR----------LLGHWRRAGMMWSQEQLSELEKAMDVIIDVFHQYS-RQEGH-KDTLTKHELKLLIEKQLANYLR-----HVKNRATIDEIMKDLDVNKDQQINFCEMMLLITRVTIATHEHLH |
| MRP-126\|Glaucidium brasilianum\|NXL39676.1 | MRP-126 | L | M | hphobic | hphobic | LNKDKELTFEEFTIVLAKVTDDAHRIMHKDDR----------LQLFLFQTTQTQEPLSELEKAMDVIIDVFHQYS-RREGD-RDTLSKKELKLLVEKQLANYLK-----HVKSKVTIDEIMKDLDVNKDEQLSFCEAMLLIIRVTIATHEHLH |
| MRP-126\|Anhinga anhinga\|NXC72277.1 | MRP-126 | L | M | hphobic | hphobic | INKDKEVNFKEFTIVLAKVADDAHRLFHHQDRGQWDAFGCTGPRGAC----------LSELEKAMDVIIDVFHQYS-RREGD-RDTLTKNELRLLIEKQLVNYLR-----HVKNKATIDQIMSDLDINKDQQISFCEVMLLITRVTIATHEHLH |
| MRP-126\|Alca torda\|NWX73657.1 | MRP-126 | L | M | hphobic | hphobic | ----------QAQQQLSELEKAMDVIIDVFHQYS-RREGD-RDTLTRNELKLLIEKQLVNYLK-----HVRNKATIDQIMKDLDTNGDAQICFSEVMLLITRVTIATHEHLH |
| MRP-126\|Phaetusa simplex\|NXW37636.1 | MRP-126 | L | M | hphobic | hphobic | LNKDKELTFEEFTIVLAKLTNDAHHLIHKEDR----------LPSFLLQNPQAQEQLSELEKAMDVIIDVFHQYS-RREGD-RDTLTKNELKLLIEKQLVNYLR-----HVRNKATIDQIMKDLDVNRDAQISFSEVMLLITRVTIATHEHLH |
| MRP-126\|Sula dactylatra\|NWI33292.1 | MRP-126 | L | M | hphobic | hphobic | LNKDKDVSFKEFTTVLAKVADDAHRISHNDER----------LSSFLFQTPQVQEQLSELEKAMDVIIDVFHQYS-RREGD-RDTLTKNELRLLIEKQLVNYLR-----HVKNKATIDQIMTDLDINKDQQISFCEVMLLITRVTIATHEHLH |
| MRP-126\|Centropus unirufus\|NWR80267.1 | MRP-126 | L | M | hphobic | hphobic | ----------QTQEPLSELEKAMNAIIDVFHNFS-RREGD-RDTLTKQELKLLIEKQLANYLR-----HVKNQATIDQIMTDLDVNKDAQISFCEVMMLITRVTIATHEHLH |
| MRP-126\|Fregata magnificens\|NWH46225.1 | MRP-126 | L | M | hphobic | hphobic | LNKDKELTFEEFTIVLAKLTDDAHRISHQDDRGQWDAFGCTGPC----------DTCVYWLSELEKAMDVIIDVFHQYS-RREED-RDTLTKKELKLLIEKQLVNYLR-----HVKSKATIDQIMNDLDVNKDAQISFCEVMLLITRVTIATHEHLH |
| MRP-126\|Mesitornis unicolor\|XP_010177863.1 | MRP-126 | L | M | hphobic | hphobic | ----------MSKTSQTQEPLSELEKAMDAIIDVFHQYS-RREGH-SDTLTKNELKLLIEKQLANYLR-----HVKSKATIDEIMKDLDVNKDAQISFCETMLLITRVTIATHEHLH |
| MRP-126\|Burhinus bistriatus\|NWQ90674.1 | MRP-126 | L | M | hphobic | hphobic | LNKDRQLTFEEFTIILAKLTDDAHRIIHKDDR----------TSQTQEPLSELEKAMDVIIDVFHQYS-RREGD-RDTLTKKELKLLIEAQLVNYLR-----HVRNKATIDEIMKDLDINKDAQISFCEVMLLITRVTIATHEHLH |
| MRP-126\|Arenaria interpres\|NXK24119.1 | MRP-126 | L | M | hphobic | hphobic | LNKDRELTFEEFTIILAKITDDAHRIIHNDDR-------CTP----------DMSPFVFQTPQSQEQLSELEKAMDVIIDVFHQYS-RREGD-RDTLTKSELKLLIEKQLVNYLR-----HVRNKATIDEIMKDLDINKDAQISFCEVMLLITRVTIATHEHLH |
| MRP-126\|Fulmarus glacialis\|XP_009577265.1 | MRP-126 | L | M | hphobic | hphobic | ----------TPQTQEQLSELEKAMDVIIDVFHQYS-RQEGH-KDTLTKHELKLLIEKQLANYLR-----HVKNKATIDEIMKDLDVNKDEQISFCEMMLLITRVTIATHEHLH |
| MRP-126\|Crotophaga sulcirostris\|NWS76993.1 | MRP-126 | L | M | hphobic | hphobic | ----------QTQEPLSELEKAMDVIIDVFHQYS-RREGH-HDTLTKQELKLLIEKQLANYLR-----HVVNKVTIDEIMKDLDVNKDAQISFCEVMMLITRVTIATHEHLH |
| MRP-126\|Himantopus himantopus\|NXN74949.1 | MRP-126 | L | M | hphobic | hphobic | LNKDRDLTFEEFTIVLAKLTDDAHRISHNDNRGQWDAFGCTGPCGAC----------GRQTQEPLSELEKAMDVIIDVFHQYS-RREGD-RDTLTKKELKLLIEKQLANYLR-----HVKNKATIDEIMKDLDINKDAQISFCEVMLLITRVTIATHEHLH |
| MRP-126\|Cuculus canorus\|XP_009556431.1 | MRP-126 | L | M | hphobic | hphobic | ----------MSKTSQTQEPLSELEKAMDVIIDVFHQYS-RREGD-RDTLTKQELKLMIEKQLANYLR-----HVKNKATIDEIMKDLDVNKDAQISFSEMMMLVIRVTIATHEHLH |
| MRP-126\|Sagittarius serpentarius\|NXR00391.1 | MRP-126 | L | M | hphobic | hphobic | LNKDSELTFEEFTIVLAKLTDDAHRISHRDDR----------SQFFLFQTSQIQEQLSELEKAMDVIIDVFHQYS-RREGD-RDTLTKRELKLLIENQLANYLK-----HVKTQATIDEIMKDLDINKDAQISFCEMMLLVTRVTIATHEHLH |
| MRP-126\|Lophotis ruficrista\|NXE15413.1 | MRP-126 | L | M | hphobic | hphobic | LNKDKDLTFEEFTIVLAKLTDDAHHISHGEDR----------SQSFLFQTTQTQEPLSELEKAMDVIIDVFHQYS-RREGD-RDTLTKRELKLMIEKQLANYLK-----HVKNKATIDEIMKDLDINKDAQISFCETMLLVTRVTIATHEHLH |
| MRP-126\|Alligator mississippiensis\|XP_006278580.2 | MRP-126 | M | F | hphobic | aromatic | ----------MSKKCPDAQTELEKAIETIINIFHQYS-VRVGH-FDTLTKMEMKLLIEKQLPNYMK-----NQTKPGEIDALFKDLDKNKDKQLSFGEFMVLITRVTIATHEHLH |
| MRP-126\|Crocodylus porosus\|XP_019412313.1 | MRP-126 | L | F | hphobic | aromatic | ----------SFAKMTKKCPEAQTELEQSIEKIIDVFHVYS-VRVGH-FDTLTKMELKLLIQKQLPNYLK-----NQTSPGQIDALFKDLDKNKDQQLSFGEFMVLITRVTIATHEHLH |
| MRP-126\|Chelydra serpentina\|KAG6923790.1 | MRP-126 | L | F | hphobic | aromatic | ----------MAQQQQTTIKGDLSELEKALETVISVFHKFS-KRVGH-FDTLTKGELKQLIDKQLVNYLK-----HVKNQVSIDVLFKDLDKNRDQQLSFAEFMVLITRVTIACHDHMH |
| MRP-126\|Platysternon megacephalum\|TFJ97784.1 | MRP-126 | L | F | hphobic | aromatic | ----------MLKSTQQVIKGDFSELEKAIETIINIFHQYS-VRVGH-FDTLTKGELKQLIDKQLVNFLK-----KQKDQASIDSLFKELDKNRDQQLSFGEFMVLITRVTIATHEHIH |
| MRP-126\|Dermochelys coriacea\|XP_038239226.1 | MRP-126 | L | F | hphobic | aromatic | ----------MSQVSQTLIKGDFSELEKAIETIINIFHQYS-VRHGH-FDTLTMGELRQLIDRHLVNFLK-----KQKDQTTIDSLFRELDKNKDQQLSFGEFMVLITRVTIATHEHIH |
| S100A12\|Alligator mississippiensis\|XP_014451269.2 | S100A12 | F | F | aromatic | aromatic | ----------KTQLEKELQCIVDIYHQYS-FRNPI-DDYLQKGEFKKLLKEQAADFLRNTKPPKMTLEQYLELLFEKADKNHDGHIKFTEFLTTLARIAIDAHDSSH |
| S100A12\|Pogona vitticeps\|XP_020666471.1 | S100A12 | F | F | aromatic | aromatic | ----------MKTLMEQGLECVVNIFHQYC-FGKHP-DDYLQPEEFKKMLKEQAQPFLRDTTPDGVSQDAYIDQLFKKADRDHNGKLKFTEVIVVVAGALIDAHNRSH |
| S100A12\|Lacerta agilis\|XP_033030861.1 | S100A12 | F | F | aromatic | aromatic | ----------MKTLLEKALECTVNIFHQYA-IRNPK-DDYLQFKEFEKLLKEQAQPFLKDTLPPGMTQDAYIKKMFQKADKNRDGKLKFTEFLPVLEAALVDAHNRSH |
| S100A12\|Pterocles gutturalis\|XP_010074898.1 | S100A12 | F | F | aromatic | aromatic | ----------MKTDLELALECVVNIYHQYALKCPPI-DDYLNKYEFTMLLKETASPFLHSTMPPSVSIDDYINKLFDKADGNHNGRLKFTEFLKVLTCIAIDAHKRFH |
| S100A12\|Columba livia\|XP_005512854.1 | S100A12 | F | F | aromatic | aromatic | ----------MKTDLELALECIVNIYHQYAVKCPPI-DDYLNKTEFSQLLKENAEPFLHRNMAPGLSVDKYISQLFSRADKNRDGRLRFTEFLTTLSGVVIDAHERFH |
| S100A12\|Dromaius novaehollandiae\|XP_025973895.1 | S100A12 | F | F | aromatic | aromatic | ----------MKTDLELALDCAVNIYHQYA-VKKPL-DDYLDRGEFKALLTDNAKPFLAHTTPPGLDRDVYINQLFNRADRNHNGRLKFTEFLTTLILIFIEAHKRSH |
| S100A12\|Apteryx rowi\|XP_025936684.1 | S100A12 | F | F | aromatic | aromatic | ----------MKTDLELALDCAINIYHQYS-VKKPL-DDYLSRGEFRALLTENAKPFLTNTKPPGLGIDAYIDQLFVKADHNRDGRLKFTEFLTTLNLVAIEAHKRSH |
| S100A12\|Acanthisitta chloris\|XP_009079041.1 | S100A12 | F | F | aromatic | aromatic | ----------MKTKMELAMESIINTYHQYS-ARDPL-DDYLSKPEFSQLMKETAEHFLHNTIPPNMTIDDYINKLFTKADANHDGRLKFTEFLTTLSLVAIDAHNKSH |
| S100A12\|Pelecanus crispus\|XP_009491269.1 | S100A12 | F | F | aromatic | aromatic | ----------MKTDLELALECVVNIYHQYA-MKWPI-DDYLNRTEFSMLLKETAKPFLHNTKPPNMSTDDYISSLFVRADRNRNGRLKFTEFLTTLSLVAIDAHNRFH |
| S100A12\|Mesitornis unicolor\|XP_010177864.1 | S100A12 | F | F | aromatic | aromatic | ----------MKTDLELALECIVNLYHKYA-LKYPL-DDYLSKPEFSQMLKETAKPFLQSTLPPKTSIDDYISQLFSKADHNHDGRLKFTEFLSTLSLVFIDGHNRSH |
| S100A12\|Dryobates pubescens\|XP_009905013.1 | S100A12 | F | F | aromatic | aromatic | ----------MKTDLELALECAINIYHQYA-IRKPI-DDYLSKAEFSQLLKDTAQPFLHDTTPPNMTEDAYIQKLFSQADSNHDGRLKFTEFLTTLNLALVDAHNRSH |
| S100A12\|Chiroxiphia lanceolata\|XP_032568729.1 | S100A12 | F | F | aromatic | aromatic | ----------MKTDLELALECAVNIYHQYAVLNRPI-DDYLSKTEFSKLLKETAKPFLNDTKPPNTSTDDYINQLFNKADGNHDGRLKFTEFMTTLNLVAINAHDRSH |
| S100A12\|Zosterops borbonicus\|TRZ09036.1 | S100A12 | F | F | aromatic | aromatic | ----------MKTDLELALECAVNIYHRYA-VRYPI-DDYLSRKEFSQLLQDTAQPFLRDTAPPNTSTDAYIKQLFAKADANHDGRLKFTEFLTTLSLVAIDAHNKSH |
| S100A12\|Corvus brachyrhynchos\|XP_008636383.1 | S100A12 | F | F | aromatic | aromatic | ----------MKTDLELALECAINIYHQYA-MRRPI-DDYLSKNEFSKLLKETAQPFLQNTLPPNTTTNNYINQLFTKADANHDGRLKFTEFMTTLSLVAIDAHNRSH |
| S100A12\|Passer montanus\|XP_039564619.1 | S100A12 | F | F | aromatic | aromatic | ----------MKTDLELALECAVNVYHQYA-MRYPM-DDYLSKEEFSSLLKDTAEPFLRNTVPPKTTIDSYIQQLFTKADANHDGRLKFTEFLTTLSLVAIDAHNRSH |

```
S100A12|Taeniopygia guttata|XP_030147529.3 S100A12 F F aromatic aromatic --------------------------------------------------------------------------------------------------------------------------------------------------MKTDLELALECAVNIYHRYA-VQRPV-DDYLSKNEFSKLLKETAEPPLRSTAPPNTSTDSYIQQLFTKADANHDGRLKFTEFLTTLSLVAIDAHNRSH
S100A12|Aythya fuligula|XP_032060413.1 S100A12 F F aromatic aromatic -----------------------------------------------------------------------------------------------------------------------------------MSASEQAIPRPTTRKTKTDLELALECAINIYHQYA-IKNPR-DDYLSKSEFTQLLKETAKTFLTRTVPPNMSIDDYINQLFTKADKNRDGRLKFTEFLTTLNLAAIDAHNRSH
S100A12|Anser cygnoides domesticus|XP_013056756.1 S100A12 F F aromatic aromatic -----------------------------------------------------------------------------------------------------------------------------------MSLSEQANPRATPRKMKTDLELALECAINIYHQYA-IKNPR-DDYLSKSEFTQLMKETAKTFLTNTVPPKTSIDAYINQLFARADKNRDGRLKFTEFLTTLNLAAIDAHKRSH
S100A12|Oxyura jamaicensis|XP_035202653.1 S100A12 F F aromatic aromatic -----------------------------------------------------------------------------------------------------------------------------------MSLSEKPIPKATARKMKTDLELALECAINIYHQYA-IKNPL-DDYLSKSEFTQLLKETAKTFLTNTVPPKMSIDAYINQLFTKADKNRDGRLKFTEFLTTLNLAVIEAHKRSH
S100A12|Antrostomus carolinensis|XP_028941632.1 S100A12 F F aromatic aromatic --------------------------------------------------------------------------------------------------------------------------------------------------MKTDLELALECIVNVYHQYA-VKNPI-DDYLSRSEFSELLQETAKPFLHNTLPPNTSVDSYINKLFNKADANHNGRLKFTEFLTTLVVVIDAHNRSH
S100A12|Phalacrocorax carbo|XP_009505422.1 S100A12 F F aromatic aromatic --------------------------------------------------------------------------------------------------------------------------------------------------MKTDLERALEDVVNVYHQYA-IKKPI-DDYLSKNEFSELLKENAKSFLHNTLPPNMSIDDYISKLFVTADRNHNGRLKFTEFLTTLNLAVIDAHNRSH
S100A12|Fulmarus glacialis|XP_009577264.1 S100A12 F F aromatic aromatic --------------------------------------------------------------------------------------------------------------------------------------------------MKTDLELALECAINIYHQYA-IKYPI-DDYLSKPEFSELLKENAKPFLHSTIPPNTVDDYISKLFTRADGNHDGRLKFTEFLTTLNLVVIDAHNRSH
S100A12|Haliaeetus albicilla|XP_009925850.1 S100A12 F F aromatic aromatic --------------------------------------------------------------------------------------------------------------------------------------------------MKTDLELALECAINIYHQYA-MKCPI-DDYLSKAEFSKLLKETAKPFLYNTVPPNTSIDDYISKLFARADHNHDGRLKFTEFLTTLNLVVIDAHNRSH
S100A12|Pygoscelis adeliae|XP_009320495.1 S100A12 F F aromatic aromatic --------------------------------------------------------------------------------------------------------------------------------------------------MKTDLELALECAINIYHQYA-MKCPI-DDYLSKTEFSKLLKETAKPFLYNTMPPNMTIDDYISKLFARADCNHDGRLKFTEFLTTLNLVVIDAHNRSH
S100A12|Opisthocomus hoazin|XP_009929499.1 S100A12 F F aromatic aromatic --------------------------------------------------------------------------------------------------------------------------------------------------MKTDLELALECIVNVYHQYA-MKKPI-DDYLSKTEFSDLLKETAQPFLHNTLPPKTTVDDYIRKLFTKADHNRDGRLKFTEFLTTLSLIVIEAHKRSH
S100A12|Chaetura pelagica|XP_009999834.1 S100A12 F F aromatic aromatic --------------------------------------------------------------------------------------------------------------------------------------------------MKTDLELALECVINVYHQYA-IRNPM-DDYLSKPEFSELLKENAKPFLHDTLPPNTTTDDYIKQLFTKADANRDGRLKFTEFLTTLNGVFIDAHKRSH
S100A12|Calypte anna|XP_008500809.1 S100A12 F F aromatic aromatic --------------------------------------------------------------------------------------------------------------------------------------------------MKTDLELALECVINIYHQYA-IKNPL-DDYLSKNEFSELLKKSAQPFLHNTLPANTSVDDYISKLFTKADGNHDGRLKFTEFLTTLNLVVIEAHKRSH
S100A12|Calidris pugnax|XP_014816950.1 S100A12 F F aromatic aromatic --------------------------------------------------------------------------------------------------------------------------------------------------MKTDLELALECIINVYHQYA-VKKPI-DDYLSKDEFSELLKENAKPFLHNTSPPNTPVDDYIANLFKKADRNRDGRLKFTEFLTTLNLVVIDAHNRSH
S100A12|Apaloderma vittatum|XP_009874754.1 S100A12 F Y aromatic aromatic --------------------------------------------------------------------------------------------------------------------------------------------------MKTDLELALECVINIYHQYA-VKRPI-DDYLSKEEFSELLKKDAQPFLHNTMPPNTTVDDYIGKLYAKADSNHDGRLKFTEFLTTLNLVFIDAHNRSH
S100A12|Cariama cristata|XP_009701719.1 S100A12 F F aromatic aromatic --------------------------------------------------------------------------------------------------------------------------------------------------MKTDLELALECTINIYHQYS-IKYPR-DDFLSKSEFTSLLEETAQPFLRNTVPPNMSINDYISKLFIKADHNHDGRLKFTEFLTTLNLVAIDAHNRSH
S100A12|Chlamydotis macqueenii|XP_010124165.1 S100A12 F F aromatic aromatic --------------------------------------------------------------------------------------------------------------------------------------------------MKTDLEHALECVVNIYHQYAVRNPPI-DDYLNKEFFSKLLKETAKPFLHNTMPPNTSVDDYISKLFAKADRNHDGRLKFTEFLTTLNLVVIDAHNRSH
S100A12|Athene cunicularia|XP_026719717.1 S100A12 F Y aromatic aromatic --------------------------------------------------------------------------------------------------------------------------------------------------MKTDLELALECAVNIYHQYA-IKNPI-DDYLSKDEFSQLLKENAKPFISNTVPPNTSIDDFINTLYTRADGNHDGRLKFTEFLTTLNLLVIDAHNRSH
S100A12|Charadrius vociferus|XP_009888759.1 S100A12 F F aromatic aromatic --------------------------------------------------------------------------------------------------------------------------------------------------MKTDLELALECAVNVYHHYA-IKNPL-DDYLSKTEFSQLLKETAKPFLQNTVPPNTTIDDYISKLFNKADRNRDGRLKFTEFLTTLNLVAIDAHNRSH
S100A12|Strigops habroptila|XP_030330011.1 S100A12 F F aromatic aromatic --------------------------------------------------------------------------------------------------------------------------------------------------MKTDLELALECAVNIYHQYA-IKNPI-DDYLDKSEFSQLLKETAKPFLHNTMPPNMSADEYISKLFAKADSNHDGRLKFTEFLTTLNLVVIDAHNRSH
S100A12|Tyto alba alba|XP_009973221.2 S100A12 F F aromatic aromatic --------------------------------------------------------------------------------------------------------------------------------------------------MKTDLELALECVVNIYHQYA-IKYPI-DDYLNKTEFSDLLKETAKPFLHNTMPPNVSIDDYINKLFTRADANHDGRLKFTEFLTTLNLVAIDAHNRSH
S100A12|Paroedura picta|GCF61138.1 S100A12 F F aromatic aromatic --------------------------------------------------------------------------------------------------------------------------------------------------MKTDLELALECVINIYHQYA-IKNPL-DDYLSKTEFSELLKETAKPFLHNTMPPNLDQEAYIKQLFQEADKNKDGNVKFTEFLCTFARFLDNAHHRSH
S100A12|Paroedura picta|GCF61139.1 S100A12 F F aromatic aromatic --------------------------------------------------------------------------------------------------------------------------------------------------MKTKLEQALDCVVHIYHQYC-ILDPV-DDYLHLREFQKLVKEQAQPFLQATTPPGLDPDTYICQLFRNADRDQSGYLKFTEFLWTFASILDYEHEKSH
S100A12|Xenopus laevis|ARQ79451.1 S100A12 M M hphobic hphobic -----------------------------------------------------------------------------------------------------------------------------------------------------------VIHEIIEIFLSYS-GGECG-PHKLNKHEMDKLIQNEFSDVIK----NP-KDPKTVEALMKVLDQNKDEEVDFDEFSELLCKVLKAT-RAMQ
S100A12|Hippocampus comes|XP_019734136.1 S100A12 F F aromatic aromatic ---------------------------------------------------------------------------------------------------------------------------------------------------MFKLEECIQKF-DLFEEYA---SED-NDCLSRDEFKALVIDEIPDY-K----GK-LEKCDIDEVFDELDKNDDDQLSFQEYIMCMVKMTKCQRKKAG
S100A12|Epinephelus lanceolatus|XP_033492604.1 S100A12 V F hphobic aromatic ---------------------------------------------------------------------------------------------------------------------------------------------------MSNLQMALCNLIATFDQYA-KKDGD-NKSLTKAEVKDLLKSELGELL-----GKANDQAAMDSVFTAMDINQDETVNFEEFIIMFAGLATVLHNTVTS
S100A12|Gekko japonicus|XP_015265129.1 S100A12 L F hphobic aromatic -------------------------------------------------------------------------------------------------------------------------------------------------MPLTEMEQCIQTILKHFHESA-GKEGN-YDTLNKAELKGFITKNFPNYMK-----KVTDPSILDNVFEQMDTNKDQEISFGEIMQFVTIVAILSHDRLH
S100A12|Paroedura picta|GCF56322.1 S100A12 F F aromatic aromatic -------------------------------------------------------------------------------------------------------------------------------------------------MSKTLEKCCDCFITMFHKYS-VRTDH-FDRLSLKEFKTMLEEQLPHYI-----EKPKDPAALKRLFDELDKNKDKEIDFTEWMGWTGKVLIAAHNHIHI
S100A12|Rousettus aegyptiacus|KAF6397907.1 S100A12 L L hphobic hphobic --------------------------------------------------------------------------------------------------------------------------------------------------MSKLENSLREVIAVFRQYA-EKKCK-KDTMSKGELKRLQLILIEMADSIK-----NTKEKHIIQKILKELDDDRDDQIDFFHSVVKMTQVLKTT----I
S100A12|Crotalus tigris|XP_039219436.1 S100A12 F F aromatic aromatic -------------------------------------------------------------------------------------------------------------------------------------------------MCKTQLEKCCEYIINIFHQYS-IRTPP-PDMLSKGEFAKMVKEQLPNVIR-----NTKNPEAMNRFFNELDQNKDGQVSFDEFVNLVSRLLTST--TSTI
S100A12|Python bivittatus|XP_007443023.1 S100A12 L F hphobic aromatic -------------------------------------------------------------------------------------------------------------------------------------------------MSLTQLEKDLESIIDIFHKYS-IKQEH-HDKLSKKELSRLLKEQLPNFLK-----SQKNPDAIDCIFKELDQDQDAQVSFQEFLILISRVLGNTHDNI--
S100A12|Notechis scutatus|XP_026549429.1 S100A12 L F hphobic aromatic -------------------------------------------------------------------------------------------------------------------------------------------------MPQTQMEKCCECIIDVFHKYS-MQKKP-YDMLSKAELAQLLREQLPNFIQ-----GSRDPAATDRLFKELDQNKDEKVSFDEFMRLVGTVFTVSYRDL-
S100A12|Echinops telfairi|XP_004717759.1 S100A12 M F hphobic aromatic --------------------------------------------------------------------------------------------------------------------------------------------------MTKMEEHLEGLLNIFHQYS-VRLGH-VDTLTKGEMRQLIPKELPNTLK-----NTKNPSANDKIFQELDTDRDGQIDFREFIAVVARLMVNAHDSIH
S100A12|Marmota monax|KAF7460633.1 S100A12 L R hphobic basic --------------------------------------------------------------------------------------------------------------------------------------------------MPTLEEHLEGIINIFHQHS-AHVGD-FDTLSKKELKTLITRDLANTIK-----TTCPPPALREWR---VPGEDGG-TRTGVGIPKIKLPLTEYSRIWN
S100A12|Monodelphis domestica|XP_007481979.1 S100A12 F L aromatic hphobic --------------------------------------------------------------------------------------------------------------------------------------------------MTLDENAMEKLINIFHQYS-VRVGD-FDKLSLSEFKELVKEQFPTWVK-----KPTDPESMKIVLQSFDENKDREMDFEEFMKMLVAIIITENLPIH
S100A12|Sarcophilus harrisii|XP_012403602.2 S100A12 F F aromatic aromatic -------------------------------------------------------------------------------------------------------------------------------------------------KMTDLENSCETIVNIFHQYS-VRTDD-FDTLSFNEFNRLVKEQFPNWLG-----RVNDPEFITNTFRSVDKNKDKQVDFGEFVNLLIPLIKATHIIFMI
S100A12|Erinaceus europaeus|XP_016044694.1 S100A12 L F hphobic aromatic --------------------------------------------------------------------------------------------------------------------------------------------------MTKLEDHLDGIINIFHQYS-VRLGH-SDTLSKGELMQLITKELANFLQ-----HVIDQASVDKIFKDLDADQDTQVNFSEFANLMITLLATHEEFHI
S100A12|Pan paniscus|XP_003817231.1 S100A12 L F hphobic aromatic --------------------------------------------------------------------------------------------------------------------------------------------------MTKLEEYLEGIVNIFHQYS-VRKGH-FDTLSKGELKQLLTKELANTIK-----NIKDKAVIDKIFQGLDANQDEQVDFQEFISLVAIALKAAHYHTHI
S100A12|Homo sapiens|NP_005612.1 S100A12 L F hphobic aromatic --------------------------------------------------------------------------------------------------------------------------------------------------MTKLEEHLEGIVNIFHQYS-VRKGH-FDTLSKGELKQLLTKELANTIK-----NIKDKAVIDEIFQGLDANQDEQVDFQEFISLVAIALKAAHYHTHI
S100A12|Trachypithecus francoisi|XP_033052964.1 S100A12 L F hphobic aromatic --------------------------------------------------------------------------------------------------------------------------------------------------MTKLEEHLEGIVNIFHNKS-VWTGH-FDTLSKGELKQLLPKELANTIK-----NTRDKAVINKIFQGLDANQDEQVDFQEFISLSAIALKAAHDNTHI
S100A12|Aotus nancymaae|XP_012293717.1 S100A12 L F hphobic aromatic --------------------------------------------------------------------------------------------------------------------------------------------------MTKLEEHLEGIVNIFHQYS-VRMGN-FDTLSKGELKQLMAKELVNLIK-----NAKDKDTVDKTFKVLDVNQDDEVDFQEFVSPLAVVLKAFHENIHI
S100A12|Saimiri boliviensis boliviensis|XP_003941944.1 S100A12 L F hphobic aromatic --------------------------------------------------------------------------------------------------------------------------------------------------MTKLEEHLEGIVNIFHQYS-VRTGS-PDTLSKGELKKMMAKELVNLIE-----NAKDKATVDETFQGLDANQDDQVSFQEFISLVARVLKASHENTHI
S100A12|Callithrix jacchus|XP_035135789.1 S100A12 L F hphobic aromatic --------------------------------------------------------------------------------------------------------------------------------------------------MTKLEEHLEGIVNIFHQYS-VRTGN-FDTLSKGELKQLMAKELVNLIK-----NAKDKATVDKTFQDLDANQDNQVNFQEFISLLAVVLKASHENIHI
S100A12|Choloepus didactylus|XP_037670217.1 S100A12 L F hphobic aromatic --------------------------------------------------------------------------------------------------------------------------------------------------MTKLEEHLEGIVNIFHQYS-VRVGH-FDTLTKGELKKMIQKELPNIIK-----NTKDQATIDKIFQDLDTDQDGQITFNEILLPLLVSMLMTVHKITH
S100A12|Manis pentadactyla|XP_036778080.1 S100A12 L F hphobic aromatic --------------------------------------------------------------------------------------------------------------------------------------------------MTKLEDYMEGIINIFHQYS-IRTGN-FDTLSKGELKQLMTKELPNAIE-----NTKDKAAIDDIFQDLDKDKDGQVSFDEFIVLVARVLKTAHKEIH
S100A12|Tupaia chinensis|XP_006159899.1 S100A12 L F hphobic aromatic --------------------------------------------------------------------------------------------------------------------------------------------------MSKMEDHMEGIINIFHQYS-VRVGD-FDTLSKGELKRLITKELPNTIK-----NTKDKAAIDKIFQDLDKDKDGLSFDEFMVLVISALKTVHKEIH
S100A12|Otolemur garnettii|XP_003783785.1 S100A12 L F hphobic aromatic --------------------------------------------------------------------------------------------------------------------------------------------------MSKLEEHLEGIINIFHQYS-VLTGN-YDTLSKGELKKLLTKEFSNTIQ-----NAKDKGGIDKIFEGLDANRDQQVDFVEFMTLVANLLRLTHEEIHI
S100A12|Sorex araneus|XP_004620093.1 S100A12 L F hphobic aromatic --------------------------------------------------------------------------------------------------------------------------------------------------MTKLEDHLEGIVNIFHQYS-VREGH-ADYLTKGDLKKLINKEMANILE-----QSKDPDSIDKIFQDLDADQDGQVNFEEFANLVITVLQTAHENIHI
S100A12|Microcebus murinus|XP_012646629.1 S100A12 L F hphobic aromatic --------------------------------------------------------------------------------------------------------------------------------------------------MSKLEEHLEGIVNIFHQYS-VRTGD-FDTLSKGELKRLLTKELANTIK-----NTRDKETVDKIFAGLDANRDDQVDFVEFTTLVATVLTAHENIHI
S100A12|Propithecus coquereli|XP_012514080.1 S100A12 L F hphobic aromatic --------------------------------------------------------------------------------------------------------------------------------------------------MSKLEQHMEGIINVFHQYS-IRTGD-YDTLSKGELSQLLSKELANTIK-----NTRDKETVDNIFAGLDTNRDQQVDFVEFMTLVASVLKTAHENIHI
S100A12|Elephantulus edwardii|XP_006895994.1 S100A12 L F hphobic aromatic --------------------------------------------------------------------------------------------------------------------------------------------------MSKLEEHMEGIINIFHQYS-VRVGD-FDTLSKGELKRLITRELPNTLK-----NTRDQSTIDKIFQELDRDGDAQINFREFVVLISRVLATAHDNIH
S100A12|Carlito syrichta|XP_008062091.1 S100A12 M F hphobic aromatic --------------------------------------------------------------------------------------------------------------------------------------------------MSKLEEHMEGIVNIFHQYS-VRGGH-YDRLEKGEMKLLLTKELSNTLK-----NTRDQATVDRIFQGLDTDRNQQVDFPEFVALIASMLETTHGNIHI
S100A12|Talpa occidentalis|XP_037355358.1 S100A12 L F hphobic aromatic --------------------------------------------------------------------------------------------------------------------------------------------------MTKLEDYLEGIINIFHQYS-VRVGN-FDTLSKGELKRLITKELTNTLK-----STKDQPFIDKIFKELDADGDGQVDFKEFVKLVTEVLVSAHDELHI
S100A12|Phyllostomus discolor|XP_028358930.1 S100A12 L F hphobic aromatic --------------------------------------------------------------------------------------------------------------------------------------------------TTKLEDHLEGIINVFHQHA-VQVGD-FDAFSKGELKQLITKELDNTIK-----NTKHQPTINRIFKELD-NLDGQVMFSEFISLVASVPETGHENIHI
S100A12|Desmodus rotundus|XP_024425820.1 S100A12 L F hphobic aromatic --------------------------------------------------------------------------------------------------------------------------------------------------VIKLEDYLEGIINIFHHYS-VRKGG-FDALTKGELKQLITKELANTIK-----NTKHQPTINKIFRELDANLDGQVIFSEFISLVASVQETGHENIHI
S100A12|Sturnira hondurensis|XP_036922954.1 S100A12 L F hphobic aromatic --------------------------------------------------------------------------------------------------------------------------------------------------MTKLEDYLEGVINIFHQYA-VRVGD-FNALSKGELKQLITQELANTIK-----STKHQPTSDKIFQELGANLDGQDMFSEFISLVASVQEAGRENIHI
S100A12|Artibeus jamaicensis|XP_036982516.1 S100A12 L F hphobic aromatic --------------------------------------------------------------------------------------------------------------------------------------------------MTKLEDYLEEIINIFHQYC-VRVGD-FNALSKGELKQLITKELANTIK-----NTKHQPTINKIFQELDANLDGQVMFSEFISLVASVQETGHENIHI
S100A12|Castor canadensis|XP_020034905.1 S100A12 L F hphobic aromatic --------------------------------------------------------------------------------------------------------------------------------------------------MSTFENYLEGIINIFHQYS-VRVGH-FDTLSKKELKKLVTKELANTIK-----NTKDQATIDKIFQDLDENGDGQVTFKEFLFLVASVLEISHDNIHI
S100A12|Ictidomys tridecemlineatus|XP_005331307.1 S100A12 M F hphobic aromatic --------------------------------------------------------------------------------------------------------------------------------------------------MTKLEEHLEGIINIFHQCS-ALVGN-YDTLSKSEMKKLITRELANTIK-----NTQDPATIDRIFQDLDDNQDGQVDFNEFVSLVASVLETAHKNLHI
S100A12|Marmota marmota marmota|XP_015358169.1 S100A12 L F hphobic aromatic --------------------------------------------------------------------------------------------------------------------------------------------------MPTLEEHLEGIINIFHQHS-AHVGD-FDTLSKKELKTLITRDLANTIK-----NTQDQATIDRIFQDLDDNRDGQVDFNEFLLLVASVLETTHKNLHI
S100A12|Dipodomys ordii|XP_012870575.1 S100A12 L F hphobic aromatic --------------------------------------------------------------------------------------------------------------------------------------------------MTKLEEHLEGIINIFHQYS-IRTDH-PDTLTKSELKKLIIRELANTIK-----NTKDQATIDRIFKELDDNGDGEVSFREFLFLASVLETAHENIHI
S100A12|Oryctolagus cuniculus|XP_002715562.2 S100A12 L F hphobic aromatic --------------------------------------------------------------------------------------------------------------------------------------------------MTKLEDHLEGIINIFHQYS-VRTGH-YDTLSKCELKKLITTELVNTIK-----NTKDQATVDRIFRDLDEDGDHQVDFKEFLSLASVLVTAHENIHI
S100A12|Ochotona curzoniae|XP_040854851.1 S100A12 L F hphobic aromatic --------------------------------------------------------------------------------------------------------------------------------------------------MTKLEEHLEGIINIFHQYS-VRTGH-YDTLTKCELKKLITTELANTIK-----NTKDRPAVDQIFRDLDEDRDGQVDFNEFLSLLASVLLTTHDNIHI
S100A12|Dasypus novemcinctus|XP_004454076.1 S100A12 L F hphobic aromatic --------------------------------------------------------------------------------------------------------------------------------------------------MTKLEDHMEGVINVFHQYS-VRWGH-HDTLSKGELKQLILKELTNTLQ-----NTKDQATIDKIFNDLDANDDGQITFNEFVPLIATLVMTMHDNIHI
S100A12|Loxodonta africana|XP_003414950.1 S100A12 L F hphobic aromatic -------------------------------------------------------------------------------------------------------------------------------------------------MSTTLEEYMEGVINVFHRYS-VRVGD-FDTLTKGELKQLITRELPKTLQ-----NSKDKASIEKLFQELDTDRDGQISFGEFMKLLVQLVLTAHNHIH
S100A12|Nyctereutes procyonoides|CAD7674840.1 S100A12 M F hphobic aromatic --------------------------------------------------------------------------------------------------------------------------------------------------MTKLEDHLEGIINIFHQYS-ARVGD-PDTLSKGELKQLIVKELPNTLK-----NTKDQATVDKLFQDLDADKDGQVNFNEFISLVSRVLVTSHENIH
S100A12|Condylura cristata|XP_004693684.1 S100A12 L F hphobic aromatic --------------------------------------------------------------------------------------------------------------------------------------------------MTKLEDHLEGIINVYHHYS-VRVGD-YDTLSRGELKRLITKELANTIK-----NNKDQATIDKIFQDLDTNKDGEVTFNEFIKLVSTVLLTAHQDIHI
S100A12|Trichechus manatus latirostris|XP_004388693.1 S100A12 L F hphobic aromatic -------------------------------------------------------------------------------------------------------------------------------------------------MSTTLEDYMEGVINVFHRYS-VRVGH-FDTLSRGELKQLITRELPNTLK-----NTKDKAAIDKLFQDLDADRDGQVNFSEFIILLARVLETAHENIHI
S100A12|Hipposideros armiger|XP_019482849.1 S100A12 L F hphobic aromatic -------------------------------------------------------------------------------------------------------------------------------------------------MMTKLEEHLEGIINIFHQYS-VRVGD-FDTLSKGELKKLLITELANTIK-----NTKDQATIDRIFQELDADRDGQVDFTEFDTALVARVLETAHKNIHI
S100A12|Myotis davidii|ELK36305.1 S100A12 L F hphobic aromatic --------------------------------------------------------------------------------------------------------------------------------------------------MTKLEDHLEGVVNVFHQYS-VRVGD-FDTLNKRELKQLLTKELANTIK-----NTKDQATIDKIFQDLDADRDGQVNFNEFVALVSKVLMTTHENIHI
S100A12|Molossus molossus|XP_036128623.1 S100A12 L F hphobic aromatic --------------------------------------------------------------------------------------------------------------------------------------------------MTKLEDHLEGIINVFHQYS-ARVGD-FDTLTKRELKQLITKELANTIK-----NTKDQATIDKIFQELDVNGDGQVNFSEFIILVARVLATAHDNIHI
S100A12|Pipistrellus kuhlii|XP_036308986.1 S100A12 L F hphobic aromatic --------------------------------------------------------------------------------------------------------------------------------------------------MTKLEDHLEGIINVFHQYS-VRVGD-FDTLTKRELKQLITKELVNTIK-----NTKDQATIDKIFQDLDADRDGQVNFNEFIVLVARVLVTAHDNIHI
S100A12|Miniopterus natalensis|XP_016075618.1 S100A12 L F hphobic aromatic --------------------------------------------------------------------------------------------------------------------------------------------------MTKLEDHLEGIINVFHQYS-ARLGD-FDTLTKRELKQLISKELVNTIK-----NTKDQATIDKIFQDLDANGDGQVNFNEFIVLVSRVLLTAHENIHI
S100A12|Puma concolor|XP_025771049.1 S100A12 L F hphobic aromatic --------------------------------------------------------------------------------------------------------------------------------------------------MTKLEDHLEGIINVFHRYS-SRVGD-FDTLSKRELKQLITRELTNTLQ-----NTKDQPTIDKIFQDLDADKDGQVTFDEFVTLVSRVLRTAHVNIHI
S100A12|Rhinolophus ferrumequinum|XP_032948956.1 S100A12 L F hphobic aromatic --------------------------------------------------------------------------------------------------------------------------------------------------MTKLEDHLEGIINIFHQYS-VRVGD-FDTLVKRELKQLLSKELANTIK-----NTKDQATIDKIFEGLDTDRDGQVNFSEFIALVASVLETAHADIHI
S100A12|Globicephala melas|XP_030697789.1 S100A12 L F hphobic aromatic ------------------------------------------------------------------------------------------------------MREVSSLEVKGDTCLIKTSLAQGSSPVLLPPHFCALRGECEVREMTKLEDYLEGIINIFHQYS-IRLEN-YDTLNKSELKKLITKELPNALK-----NTKDQPSIDKVFQELDADKDGQVNFSEFVVLVSKVLETAHADIH
S100A12|Physeter catodon|XP_007124461.2 S100A12 L F hphobic aromatic ------------------------------------------------------------------------------------------------------MREVSSLAVKGDTCLIKTSLVQGSSPVLLPPHFCALRGECEVREMTKLEDYLEGIINIFHHYS-IRVGD-YDTLSKCELSQLITKELPNALK-----NTKDQPSIDKIFQELDADKDAQITFEEFVVLVSRVLQTAHEAIH
S100A12|Balaenoptera physalus|KAB0389424.1 S100A12 L F hphobic aromatic --------------------------------------------------------------------------------------------------------------VKGDTCLINTSLAQGSSPVLLPPHFCALRGECEVREMTKLEDHLEGIINIFHQYS-VRLGH-YDTLSKCELMQLITKELPNALK-----NTKDQPSVDKVFQELDADKDGQVNFEEFIVLVSRVLETAHEAIH
S100A12|Balaenoptera musculus|XP_036700079.1 S100A12 L F hphobic aromatic -------------------------------------------------------------------------------------------------------------------------------------------IEGECEVREMTKLEDHLEGIINIFHQYS-IRVGH-YDTLSKCELMQLITKELPNALK-----NTKDQPSVDKVFQELDADKDGQVNFEEFIVLVSRVLETAHEAIH
S100A12|Sus scrofa|NP_001153744.1 S100A12 L F hphobic aromatic --------------------------------------------------------------------------------------------------------------------------------------------------MTKLEDHLEGIINIFHQYS-VRLGH-YDTLIKRELKQLITKELPNTLK-----NTKDQATIDKIFQNLDANQDGQVNFSEFVVLVTDVLITAHDNIHI
S100A12|Bos mutus|XP_005909632.1 S100A12 L F hphobic aromatic --------------------------------------------------------------------------------------------------------------------------------------------------MTKLEDHLEGIINIFHQYS-VRVGH-FDTLNKRELKQLITKELPQTLQ-----NTKDQPTIDKIFQDLDADKDGAVSFEEFVVLVSRVLKTAHIDIHI
S100A12|Oryx dammah|XP_040086511.1 S100A12 L F hphobic aromatic --------------------------------------------------------------------------------------------------------------------------------------------------MTKLEDHLEGIINIFHQYS-VRLGD-YDTLIKRELKQLITKELPNCLN-----NTKNQATIDKIFQDLDANKDKEVSFEEFVVLVSRVLKTAHENIHI
S100A12|Cervus hanglu yarkandensis|KAF4021312.1 S100A12 L F hphobic aromatic --------------------------------------------------------------------------------------------------------------------------------------------------MTKLEDHLEGIINIFHQYS-VRVGH-YDTLSKRELKQLITKELPNTLK-----NTKDQATIDKIFQELDVDRDGQVNFEEFIVLVSRVLETAHEHIHI
S100A12|Orycteropus afer afer|XP_007946781.1 S100A12 L F hphobic aromatic --------------------------------------------------------------------------------------------------------------------------------------------------MTKLEDHMEGVINVFHQYS-VRVGH-FDTLSKKELKQLLTKELANTLK-----NTKDQATIDRIFQELDVNRDGQVSFNEFVSLVARVMVTAHENIHI
S100A12|Chrysochloris asiatica|XP_006861514.1 S100A12 M F hphobic aromatic --------------------------------------------------------------------------------------------------------------------------------------------------MTKLEDCLEGIINVFHQYS-VRVGH-FDTLSKNEMKQLITKELPNTLK-----NTKDQGAIDKIFQELSDKDGQVNFVEFIVLVAKVLETAHENIHI
S100A12|Gulo gulo|VCW76682.1 S100A12 M F hphobic aromatic --------------------------------------------------------------------------------------------------------------------------------------------------MTKLEDHLEGIINVFHQYS-VRVGH-PDKLSKGEMKKLITKELPNALK-----NTKDQASIDQTFQELDADKDGEVSFPEFISLLARVLITAHEDIHI
S100A12|Halichoerus grypus|XP_035922749.1 S100A12 M F hphobic aromatic --------------------------------------------------------------------------------------------------------------------------------------------------MTKLEDHMEGIINVFHQYS-VRVGH-YDKLSRCEMRKLITRELPNTLK-----NTKDQAAIDKIFQDLDDDKDGEVDFGEFVSLIVRVLVTAHEDIH
S100A12|Ailuropoda melanoleuca|XP_002922563.1 S100A12 M F hphobic aromatic --------------------------------------------------------------------------------------------------------------------------------------------------MTKLEDHMEGIINVFHQYS-VRVGH-YDKLSRGEMKQLIIRELPNTLK-----NTKDQATIDKIFQDLDADKDGQIDFSEFVTLVVRVLLTAHEDIHI
S100A12|Mirounga leonina|XP_034865352.1 S100A12 M F hphobic aromatic --------------------------------------------------------------------------------------------------------------------------------------------------MTKLEDHMEGIINVFHQYS-VRVGH-YDKLSKGEMKQLIIRELPNTLK-----NTKDQATIDKIFQDLDANKDGQVNFNEFVSLIVSVLVTAHEDIH
S100A12|Callorhinus ursinus|XP_025714672.1 S100A12 M F hphobic aromatic --------------------------------------------------------------------------------------------------------------------------------------------------MTKLEDHLEGIINVFHQYS-VRVGH-YDKLSNSEMTQLIRRELPNTLR-----NTKEQATIDKIFQELGADKDGEVSFKKFVSLIVNVLLTAHDDIH
S100A12|Odobenus rosmarus divergens|XP_004403098.1 S100A12 M F hphobic aromatic --------------------------------------------------------------------------------------------------------------------------------------------------MTKLEDHMEGIINVFHQYS-VRVGH-YDKLSKGEMKQLITRELPNTLQ-----NTKEQATIDKIFQDLDADKDGEVTFKEFVSLIVSVLVTAHDDIH
S100A12|Ceratotherium simum simum|XP_004436108.1 S100A12 L F hphobic aromatic --------------------------------------------------------------------------------------------------------------------------------------------------MTKLEDHLEGIINVFHQYS-VRVGH-YDTLSKGELKKLIKRELPNTLE-----NTKDQATIDKIFQNLDADKDGEVNFKEFIILVAEVLETAHENIHI
S100A12|Rousettus aegyptiacus|XP_036091158.1 S100A12 L F hphobic aromatic --------------------------------------------------------------------------------------------------------------------------------------------------MTKLEDHLEGIINVFHQYS-VRVGN-FDTLSKRELKQLLTRELANSLK-----NTKDQPTIDRIFQDLDADRDGQVNFNEFASLVAQVLKTAHVNIHI
S100A12|Galeopterus variegatus|XP_008565858.1 S100A12 L F hphobic aromatic --------------------------------------------------------------------------------------------------------------------------------------------------MTKLEDHMEGIINIFHQYS-VRVGH-FDTLSKGELKQLLTRELANTIK-----NTKDKPTIDKIFQDLDSDRDGQVNFKEFISLVASVLETAHENIHI
S100A12|Equus asinus|XP_014720468.1 S100A12 L F hphobic aromatic --------------------------------------------------------------------------------------------------------------------------------------------------MTKLEDHLEGIINVFHQYS-VRVGH-FDTLSKRELKQLITRELANSIK-----NTKDKATIDQIFQDLDADKDGQVNFNEFVVLVCKVLETAHEDIHI
S100A12|Suricata suricatta|XP_029789124.1 S100A12 L F hphobic aromatic --------------------------------------------------------------------------------------------------------------------------------------------------MTKLEDHLEGIINVFHQYS-VRVGH-YDTLSKGELKQLITKQLPNSLK-----NTKDQATIDKIFQDLDADKDGQVNFEEFVVLVCRVLQTAHENIHI
S100A12|Vicugna pacos|XP_006214797.1 S100A12 L F hphobic aromatic --------------------------------------------------------------------------------------------------------------------------------------------------MTKLEDHLEGIINIFHQYS-VRVGH-YDTLSKRELKQLITQLGNTLK-----NTKDQATIDKIFQDLDADKDGQVNFEEFIALVARVLLTAHEEIHI
S100A12|Ornithorhynchus anatinus|XP_028911694.1 S100A12 L F hphobic aromatic -----------------------------------------------------------------------------------------------------------------------------------------------------MSLESHLEGVVNVFHQYS-VRVGD-FDQLSKKELKQLIEKELPNYKK-----KPKNPEAMDALFKDLDQNRDQEINFGEFMRFITAVLIATHDYIH
S100A12|Tachyglossus aculeatus|XP_038624872.1 S100A12 L F hphobic aromatic -----------------------------------------------------------------------------------------------------------------------------------------------------MSLESHLEGVVNVFHTYS-VRVGD-FDQLSKKELKQLIEKELPNYKK-----KPKDAEAMAALFKDLDQNRDQEVSFGEFMRFITTVLIATHEHIH
S100A12|Penelope pileata|NXC49113.1 S100A12 L F hphobic aromatic LNKDKELTFEEFTIVLAKVCDDAHRIIHKDDR--------------------------------------------------------------------------------------------SSRAWSVTQGCQTQGPLSELEKSMDVIIDVFHQYS-RREGD-RDTLTKRELKLMIEKQLANYLK-----HVKSQATIDQIFKDLDSNKDGLSFGEVMRLIVHVTVATHEHLH
S100A12|Callipepla squamata|OXB53812.1 S100A12 L F hphobic aromatic LNKDKELTFEEFTIVLAKVTDDAHRIIHKDDR------------------------------------------------------------------------------------------YTCCDPDLILFIRFSQGCQTQGPLSELEKAIDVIIDVFHQYS-RREGD-RDTLTRKELKLLIEKQLANYLK-----HVKNQASIDQIFKDLDVNKDDQLSFGEVMLLIVRVIVATHEHLH
S100A12|Aegotheles bennettii|NWX11416.1 S100A12 L M hphobic hphobic LNKDKEVSFEEFTIILSKLADDAHRIINHGSDR-----------------------------------------------------------------------------------------------SLDQAAQRGCGSQEQLSDLEKSMDVIIDAFHQYS-RREGD-RDTLTKKELKLLIEKQLANYLR-----HVKNKATIDEIMKDLDINKDEQISFCEMMLLITRVTIATHEHLH
S100A12|Chelonia mydas|XP_007067898.1 S100A12 L F hphobic aromatic ---------------------------------------------------------------------------------------------------------------------------------------MSQVSQTLIKGEFSELEKAIETIINIFHQYS-VRVGH-WDTLTKKELRQLIDRHLVNFLK-----TRKDHSAIDSLFSELDRNKDQQLSFGEFMVLITRVTIATHEHIH
```

**Video S1: Morph between the crystal structure of hS100A9 and the NMR structure of hS100A9/M63F**

Available for download as a mov file:
https://github.com/harmslab/stability-constraint-ms-figures/blob/a214271747be3ffc95203aa3f50010673e26c8bc/video_s1.mov
